# periscope: sub-genomic RNA identification in SARS-CoV-2 Genomic Sequencing Data

**DOI:** 10.1101/2020.07.01.181867

**Authors:** Matthew D Parker, Benjamin B Lindsey, Shay Leary, Silvana Gaudieri, Abha Chopra, Matthew Wyles, Adrienn Angyal, Luke R Green, Paul Parsons, Rachel M Tucker, Rebecca Brown, Danielle Groves, Katie Johnson, Laura Carrilero, Joe Heffer, David G Partridge, Cariad Evans, Mohammad Raza, Alexander J Keeley, Nikki Smith, Ana Da Silva Filipe, James G Shepherd, Chris Davis, Sahan Bennett, Alain Kohl, Elihu Aranday-Cortes, Lily Tong, Jenna Nichols, Emma C Thomson, The COVID-19 Genomics UK (COG-UK) consortium, Dennis Wang, Simon Mallal, Thushan I de Silva

## Abstract

We have developed periscope, a tool for the detection and quantification of sub-genomic RNA (sgRNA) in SARS-CoV-2 genomic sequence data. The translation of the SARS-CoV-2 RNA genome for most open reading frames (ORFs) occurs via RNA intermediates termed “sub-genomic RNAs”. sgRNAs are produced through discontinuous transcription which relies on homology between transcription regulatory sequences (TRS-B) upstream of the ORF start codons and that of the TRS-L which is located in the 5’ UTR. TRS-L is immediately preceded by a leader sequence. This leader sequence is therefore found at the 5’ end of all sgRNA. We applied periscope to 1,155 SARS-CoV-2 genomes from Sheffield, UK and validated our findings using orthogonal datasets and *in vitro* cell systems. Using a simple local alignment to detect reads which contain the leader sequence we were able to identify and quantify reads arising from canonical and non-canonical sgRNA. We were able to detect all canonical sgRNAs at expected abundances, with the exception of ORF10. A number of recurrent non-canonical sgRNAs are detected. We show that the results are reproducible using technical replicates and determine the optimum number of reads for sgRNA analysis. In VeroE6 ACE2+/− cell lines, periscope can detect the changes in the kinetics of sgRNA in orthogonal sequencing datasets. Finally, variants found in genomic RNA are transmitted to sgRNAs with high fidelity in most cases. This tool can be applied to all sequenced COVID-19 samples worldwide to provide comprehensive analysis of SARS-CoV-2 sgRNA.

## Introduction

Understanding variation within sub-genomic RNA (sgRNA) synthesis within the human host may have important implications for the study of SARS-CoV-2 biology and evolution. Thanks to advances in sequencing technology and collaborative science, over 100,000 SARS-CoV-2 genomes have been sequenced worldwide to date.

The genome of SARS-CoV-2 comprises a single positive-sense RNA molecule of approximately 29kb in length. While the 1a and 1b polyproteins are translated directly from this genomic RNA (gRNA), all other proteins are translated from sgRNA intermediates (Stern and Kennedy 1980; Sola et al. 2015). sgRNAs are produced through discontinuous transcription during negative strand synthesis followed by positive-strand synthesis to form mRNA. The resulting sgRNAs contain a leader sequence derived from the 5’ untranslated region of the genome and a transcription regulating sequence (TRS) 5’ of the ORF. The template switch occurs during sgRNA synthesis due to a conserved core sequence within the TRS 5’ of each ORF (TRS-B) and the TRS within the leader sequence (TRS-L)(Zúñiga et al. 2004). The conserved core sequence leads to base pairing between the TRS-L and the nascent RNA molecule transcribed from the TRS-B resulting in a long-range template switch and incorporation of the 5’ leader sequence (Sola et al. 2015). SARS-CoV-2 produces at least nine canonical sgRNAs containing ORFs for four structural proteins (S; spike, E; envelope, M; membrane, N; nucleocapsid) and several accessory proteins (3a & b, 6, 7a & b, 8 and 10)(F. Wu et al. 2020; Zhou et al. 2020; Davidson et al. 2020). In SARS-CoV ORFs 3b and 7b are considered nested ORFs and not thought to be translated from their own sub-genomic RNA (Inberg and Linial 2004). Understanding variation within sgRNA synthesis within the human host may have important implications for the study of SARS-CoV-2 biology and evolution.

Beyond the regulation of transcription, sgRNA may also play a role in the evolution of coronaviruses and the template switching required for sgRNA synthesis may explain the high rate of recombination seen in coronaviruses (Simon-Loriere and Holmes 2011; H.-Y. Wu and Brian 2010). While the majority of sgRNA relate to known ORFs, novel, non-canonical sgRNA are also produced (Nomburg, Meyerson, and DeCaprio 2020; Finkel et al. 2020; Kim et al. 2020), although the biological function of this is unclear. sgRNAs have also been shown to modulate host cell translational processes (Patel et al. 2013).

The ARTIC Network (“Artic Network” 2020) protocol for the sequencing of the SARS-CoV-2 (Figure 1A) has been employed worldwide to characterise the genetic diversity of this novel coronavirus. The COVID-19 Genomics Consortium (COG-UK)(“An Integrated National Scale SARS-CoV-2 Genomic Surveillance Network” 2020) in the UK, alone, has produced 16826 ARTIC nanopore genome sequences (Correct 29th October 2020), while internationally GISAID contains thousands more similar datasets (8775 with “nanopore” in the metadata and 3660 list “artic”, 14th June 2020). This protocol involves the amplification of 98 overlapping regions of the SARS-CoV-2 genome in two pools of 49 amplicons to provide full sequence coverage when sequenced with Oxford Nanopore sequencing devices. All known SARS-CoV-2 ORF TRS sites are contained within one or more amplicons in this panel (Figure 1A). Other methods of enrichment and enrichment-free sequencing of the SARS-CoV-2 genome like bait based capture and subsequent short read Illumina sequencing or metagenomics, respectively, are also popular and hold promise for the detection of sgRNA.

**Figure 1.**
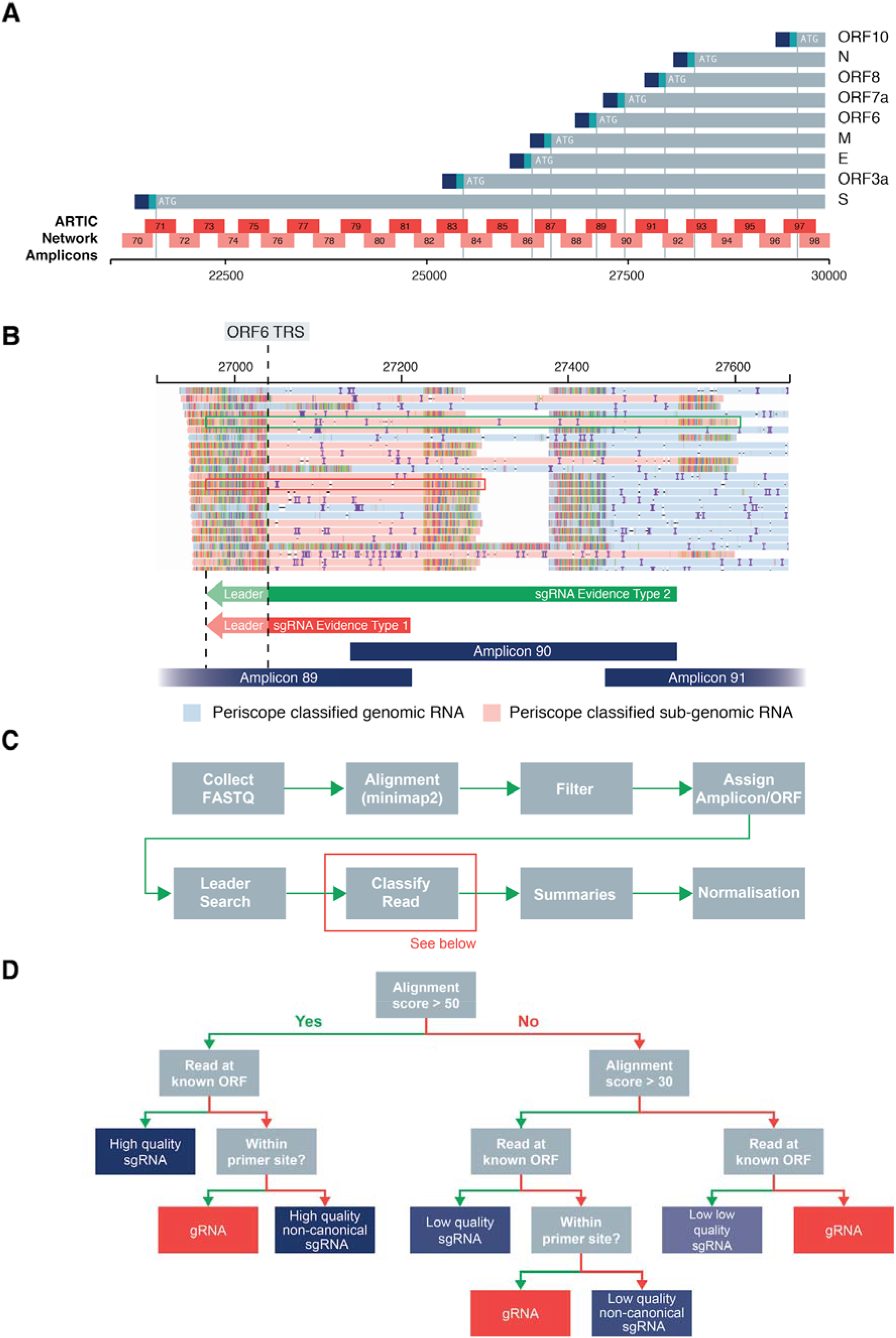
Periscope ARTIC Nanopore Algorithm Design Details. ***A.** ARTIC network amplicon layout with respect to ORF TRS positions of SARS-CoV-2. Blue and aqua at the end of each ORF signifies leader and TRS respectively. **B.** Read pileup at ORF6 TRS showing two types of reads which support the existence of sgRNAs. Type 1 (Red): Result from 3’->5’ amplification from the closest primer to the 3’ of the TRS site, and Type 2 (Green): These result from 3’->5’ amplification from the adjacent amplicons 3’ primer (i.e. the second closest 3’ primer). **C.** Overview of periscope workflow. **D.** Decision tree for read classification. Green arrow denotes a “Yes” for the step in question, i.e. if the read is at a known ORF start site a green arrow is used, if not a red arrow for “No” is used.*

It has previously been shown that sgRNA transcript abundances can be quantified from full RNASeq data by calculation of Reads Per Kilobase of transcript, per Million mapped reads (RPKM) or by using so-called “chimeric” fragments containing the leader and TRS (Irigoyen et al. 2016). From two independent repeats, the R^2^ between these two measurement methods was 0.99. Studies of SARS-CoV-2 sgRNA have used methods which specifically detect expressed RNA, such as direct RNA sequencing of cultured cells infected with SARS-CoV-2 (Davidson et al. 2020; Taiaroa et al. 2020; Kim et al. 2020) or more traditional total PolyA RNASeq (Finkel et al. 2020). We hypothesised that we could detect and quantify the levels of sgRNA to both identify novel non-canonical sgRNA and provide an estimate of ORF sgRNA expression in SARS-CoV-2 sequence data. Here we present a tool for these purposes and its application to 1155 ARTIC Nanopore-generated SARS-CoV-2 sequences derived from clinical samples in Sheffield, UK and validate our findings in data from independent SARS-CoV-2 sequences from Glasgow, UK, in addition to Illumina data generated with bait capture and metagenomic approaches.

## Results

### Evidence for Sub-Genomic RNA

We designed a tool, periscope (https://github.com/sheffield-bioinformatics-core/periscope), to re-analyse raw data from SARS-CoV-2 isolates to identify sgRNA based on the detection of the leader sequence at the 5’ end of reads as described previously (Leary et al. 2020).

#### ARTIC Network Nanopore Sequencing Data

The recommended bioinformatics standard operating procedure to process ARTIC network sequencing data to produce a consensus sequence involves selecting reads between 400 and 700 base pairs, and the trimming of the primer and adapter sequence. In most cases this removes reads which might provide evidence for sgRNA. Mapping raw data from this protocol reveals the presence of reads at ORF TRS sites which are sometimes shorter (Supplementary Figure S1 - sgRNA) than the full ARTIC Network amplicon and contain leader sequence at their 5’ end. We believe these reads are the result, in the case of Pool 1, priming from primer 1 of the pool, which is homologous to most of the leader sequence. We also see, in both pools, uni-directional amplification from the 3’ primer, which results in a truncated amplicon when the template is a sgRNA (Figure 1B). Interestingly, we also see longer reads which are the result of priming from the 3’ end of the adjacent amplicon (Figure 1B).

To separate gRNA from sgRNA reads, we employ the following workflow using snakemake(Köster and Rahmann 2012); Raw ARTIC Network Nanopore sequencing reads that pass QC are collected and aligned to the SARS-CoV-2 reference, reads are filtered out if they are unmapped or supplementary alignments. We do not perform any length filtering. Each read is assigned an amplicon. We search the read for the presence of the leader sequence (5’-AACCAACTTTCGATCTCTTGTAGATCTGTTCT-3’) using a local alignment. If we find the leader with a strong match it is likely that that read is from amplification of sgRNA. We assign reads to an ORF. Using all of this information we then classify each read into genomic, canonical sgRNA or non-canonical sgRNA (Figure 1D) and produce summaries for each amplicon and ORF including normalised pseudo-expression values.

#### Illumina Sequencing Data

Next we wanted to investigate whether we could employ a similar method to Illumina sequencing data. Illumina sequencing data for SARS-CoV-2 has been generated using three main approaches: amplicon based (ARTIC Network), bait capture based, or using metagenomics on *in vitro* samples. Here we describe sgRNA detection in both Illumina bait based capture and Illumina metagenomic data from *in vitro* exp. The reads from these techniques tend to have differing amounts of leader at the 5’ end of the reads (Supplementary Figure S2A). This is due to library preparation methods employed in these workflows. We therefore implemented a modified method for detecting sgRNA to ensure that we could capture as many reads originating from sgRNA as possible. In our Illumina implementation (Supplementary Figure S2C&D) we extract the soft-clipped bases from the 5’ end of reads and use these in a local alignment to the leader sequence. In addition to adjusting the leader detection method we also process mate pairs, ensuring both reads in the pair are assigned the same status.

### Detection of Sub-Genomic RNA

We were able to detect sgRNAs with a high leader alignment score from all canonical ORFs in multiple samples (Figure 2, Supplementary Table S1). As shown in Figure 2A, sgRNA from the N & M ORFs were the most abundant sgRNA (dependent on normalisation method), with N being found in 97.3% of Sheffield samples, consistent with published reports *in vitro* (Alexandersen, Chamings, and Bhatta 2020; Finkel et al. 2020; Kim et al. 2020). To demonstrate that the levels of sgRNA detected in the Sheffield dataset were not site specific, we applied periscope to an independent dataset of 55 ARTIC Network Nanopore sequenced SARS-CoV-2 samples from Glasgow, UK (Figure 2A).

**Figure 2.**
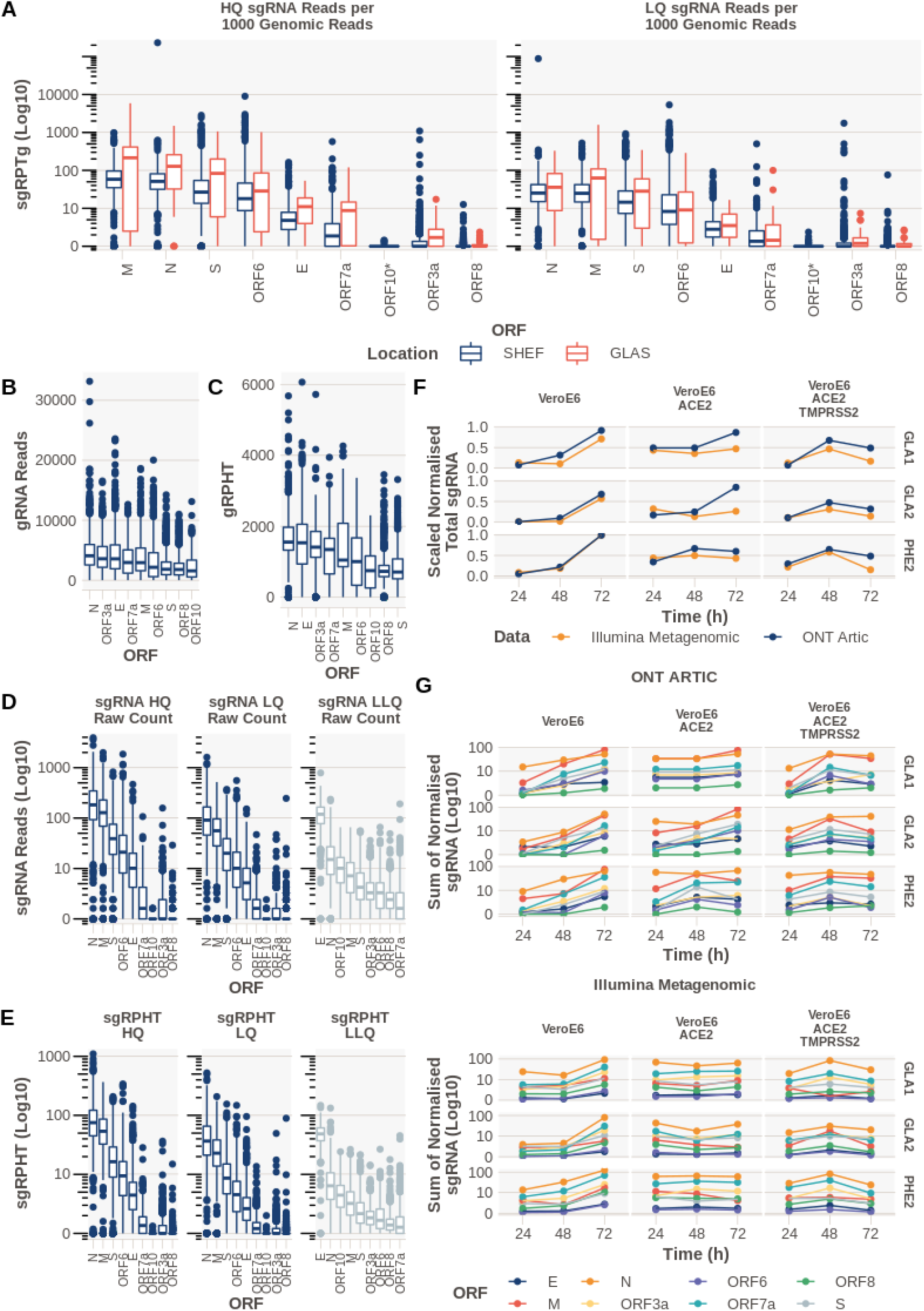
*In Vivo and In Vitro* Detection and Quantification of Canonical sub-genomic RNA in SARS-CoV-2. ***A.** The abundance of sgRNA detected for each ORF normalised per 1000 gRNAs from Oxford Nanopore (ONT) ARTIC data from both Sheffield (n=1155) and Glasgow (n=55); sgRPTg = sgRNA reads per 1000 gRNA reads. Ordered by median. *See Supplementary Figure S3 for ORF10 investigation. **B.** Number of reads supporting gRNA at each ORF. If multiple amplicons cover the ORF then this represents the sum of reads for those amplicons. **C.** gRNA reads normalised per hundred thousand mapped reads (gRPHT) at each ORF. **D.** Raw counts of sgRNAs. **E.** sgRNA normalised to total mapped reads. (sgRPHT = sgRNA reads per 100,000 mapped reads) **F&G.** In Vitro infection time course with 3 SARS-CoV-2 viral isolates (GLA1, GLA2 and PHE2) in either VeroE6 cells, VeroE6 expressing ACE2 or VeroE6 expressing ACE2 and TMPRSS2, with total RNA collected and sequenced at 24, 48 and 72 hours after infection, sequenced using either ONT ARTIC or Illumina Metagenomic approaches. **F.** The sum of all normalised (to total mapped reads to allow direct comparison across ONT ARTIC and illumina metagenomic methods) sgRNA in each technology scaled to 1. **G.** Normalised quantity (to total mapped reads) of each canonical sgRNA in each technology (ONT ARTIC; Top panel, Illumina Metagenomic; Bottom panel).*

Like previously published reports (Taiaroa et al. 2020; Alexandersen, Chamings, and Bhatta 2020; Finkel et al. 2020; Kim et al. 2020), we were unable to find strong evidence of sgRNA supporting the presence of ORF10 (Supplementary Table 2, Figure 2A) with only 0.95% of samples containing high (HQ) or low quality (LQ) sgRNA calls at this ORF. We aligned the 12 reads from these samples to a reference composing of ORF10 and leader (Supplementary Figure S3). On manual review of these results ten (4 HQ) of these reads are falsely classified as sgRNA. Two reads remain, one read from each of samples SHEF-C0840 and SHEF-C58A5. These reads could represent ORF10 sgRNA as they have an almost complete match to the leader and the remainder of the reads is a strong match to ORF10.

### Normalisation of Sub-Genomic Read Abundance

Beyond defining the presence of reads which are a result of amplification of sgRNA, we hypothesized that we could quantify the level of sgRNA present in a sample using either total mapped reads or gRNA reads from the same amplicon as denominators for normalisation. Normalisation of this kind would be analogous to traditional RNAseq analysis where reads per million (RPM) are calculated to allow comparisons between datasets where the number of reads affect the amount of each transcript detected. In the case of ARTIC Network Nanopore sequencing data, which involves polymerase chain reaction (PCR) of small (~400bp) overlapping regions of the SARS-CoV-2 genome, amplification efficiency of each amplicon should also be taken into account. Because we have a median of 258,210 mapped reads for samples from the Sheffield dataset (Supplementary Figure S4), we normalised both gRNA or sgRNA per 100,000 mapped reads (gRNA reads per hundred thousand, gRPHT, or sgRNA reads per hundred thousand, sgRPHT, respectively, Figures 2C and E).

In our second approach, because of differences in amplicon performance in the ARTIC PCR protocol that leads to coverage differences in the final sequencing data (Figure 2B&C), we determine the amplicon from which the sgRNA has originated, using methods from the ARTIC Network Field Bioinformatics package(“Artic Network” 2020). We then normalise the sgRNA per 1000 gRNA reads from the same amplicon. If a sgRNA has resulted from more than one amplicon (Figure 1B) the resulting normalised counts from each amplicon are summed giving us sgRNA reads per 1000 gRNA reads (sgRPTg) for every ORF (Figure 2A). Periscope outputs the results from both methods of normalisation so that the user can decide which is more appropriate in their case and determine whether the conclusions of their analysis are consistent across both approaches.

For Illumina data we applied one further normalisation technique to allow the normalisation of bait based capture and metagenomic data. Efficiency of capture varies between probes and designs. For metagenomic data, natural fluctuations in coverage due to sequence content can exist, therefore, to try and account for this we took the median coverage for the region around each canonical ORF start site (+/− 20bp) as the denominator in the normalisation of this data.

### Sub-genomic RNA Detection *in vitro*

The kinetics of sgRNA expression during the course of a SARS-CoV-2 infection is still not well understood. We applied periscope to data generated from an infection time course (Figure 2F&G). We used both Illumina metagenomic and Nanopore ARTIC sequencing data from an *in vitro* model of SARS-CoV-2 infection. Wildtype (WT) VeroE6, VeroE6 expressing *ACE2* and VeroE6 expressing both *ACE2* and *TMPRSS2* were infected with three different SARS-CoV-2 viral isolates; PHE2 (wild type; WT), GLA1 (D614G) and GLA2 (N439K & D614G) (Supplementary Table S9) and RNA collected for sequencing at 24, 48 and 72 hours. We normalised both datasets to the total mapped reads (per 100,000) to allow direct comparison. Importantly the scaled normalised total sgRNA level detected using periscope on both sequencing technologies is strikingly similar (Figure 2F), indicating the pattern of expression is maintained between technologies. ORF N remains one of the highest expressed ORFs in both datasets (Figure 2G). The addition of *ACE2* and a combination of *ACE2* and *TMPRSS2* results in a clear difference in the kinetics of all sgRNA over WT. This data suggests that the peak of sgRNA expression is expedited by the addition of *ACE2* and *TMRPSS2*.

### Technical Replicates & Batch Effects

To assess the reproducibility of sgRNA analysis using ARTIC Network Nanopore sequencing data and periscope, we analysed two samples that were subject to four technical replicates each; cDNA was independently prepared from the same swab extracted RNA, and subject to independent amplification using the recommended ARTIC Network PCR and sequenced (Figures 3A & 3B). The pearson correlation coefficient (R) is >=0.88 for all normalised sgRNA abundances between replicates from the same sample.

**Figure 3.**
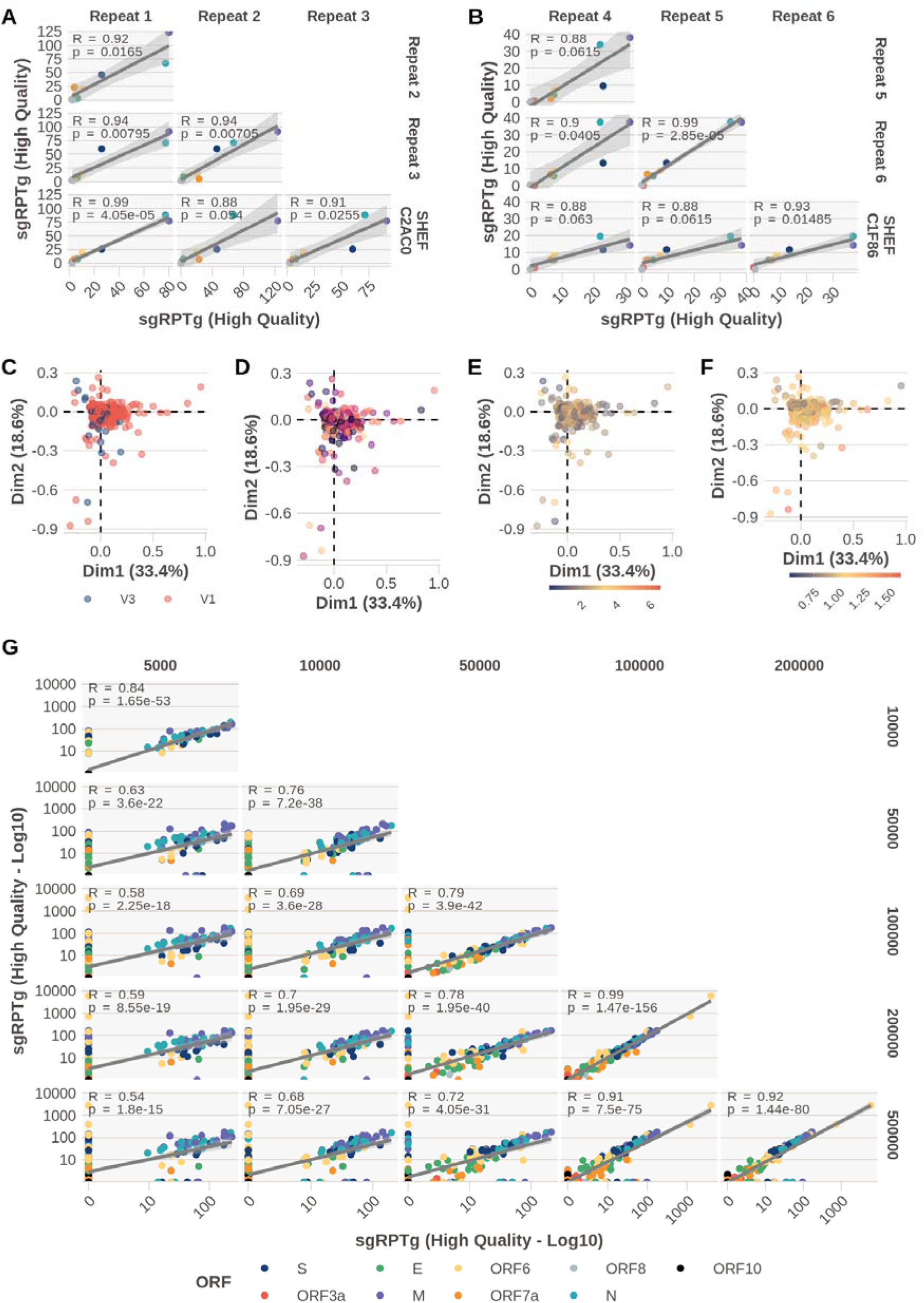
Technical Replicates, Detection Limit, and Batch Effects. ***A & B.** Four technical replicates of two samples additional to the Sheffield cohort. Pearson correlation coefficients between sgRNA normalised per 1000 gRNA reads (sgRPTg) p values adjusted with Bonferroni. (ORFS coloured according to legend in **(G)) C, D, E, F.** Unsupervised principal component analysis coloured by, ARTIC primer version V1 or V3 **(C)**, sequencing run **(D)** where the colour denotes a different run, total mapped read count (scale = 100k reads) **(E)**, or normalised E gene cycle threshold (CT) value **(F)**. **G.** Downsampling of reads from 23 high coverage (>1million mapped reads) samples. The number of reads provided as input to periscope was downsampled with seqtk(Li 2012) to 5,10,50,100,200, and 500 thousand reads.*

Next, we treated our sgRNA abundance values like an RNASeq dataset and asked whether other factors could be influencing expression. To do this we used an unsupervised principal component analysis (PCA; Figure 3C-F) and coloured samples by the different categorical variables that could affect expression (batch effects), like sequencing run, the ARTIC primer version, the number of mapped reads and ORF E gene cycle threshold (e.g. CT, Diagnostic test, normalised to RNAse P CT value, see materials and methods) as this is an indicator of the amount of virus present in an isolate and a proxy for quality (Figure 3F). There are no significant clusters between any of the above variables and the expression values in the PCA analysis.

### Lower Limit of Detection

The number of reads generated from any sequencing experiment is likely to vary between samples and between runs. The median mapped read count in the Sheffield dataset is 258,210, but varies between 9,105 and 3,260,686 (Supplementary Figure S4). In our experience we generally see much lower total amounts of sgRNA compared to their genomic counterparts, therefore, its detection is likely to suffer when a sample has lower amounts of reads (Supplementary Figure S5). To determine the effect of lower coverage on the detection of sgRNA, we downsampled 23 samples which had > 1million mapped reads to lower read counts with seqtk(Li 2012). We chose high (500k reads), medium (200k and 100k), and low read counts (50k,10k and 5k) and ran periscope on this downsampled data (Supplementary File S3). In the absence of a ground truth, we performed pairwise correlation on the abundance of sgRNA between downsampled datasets (Figure 3G). If coverage did not affect the abundance estimates, then all coverage levels would show a high correlation coefficient (pearson) when compared to each other (R>0.7, adjusted p<0.01). As expected, lower counts of 5,000 and 10,000 reads do not correlate with those generated from 100,000, 200,000 and 500,000 reads (R^2^ < 0.7). Samples with 50,000 reads seem to perform well when compared to 100,000, 200,000 and 500,000 reads with an R^2^ of 0.94, 0.89, and 0.89 respectively.

### Non-Canonical Sub-Genomic RNA

In addition to estimating sgRNA for known ORFs, we can use periscope to detect novel, non-canonical sgRNA (Figure 4). We previously applied periscope to detect one such novel sgRNA (N*) which is a result of the creation of a new TRS site by a triplet variation at position 28881 to 28883 which results in production of a truncated N ORF (Leary et al. 2020). To classify sgRNA as non-canonical, supporting reads must fulfil two criteria; First, the start position does not fall in a known TRS-B region (+/− 20bp from the leader junction), and second, in ONT data the start position must not fall within +/-5bp from a primer sequence. We chose to implement the second criteria because we noticed a pattern of novel sgRNAs being detected at amplicon edges due to erroneous leader matches to the primer sites.

**Figure 4.**
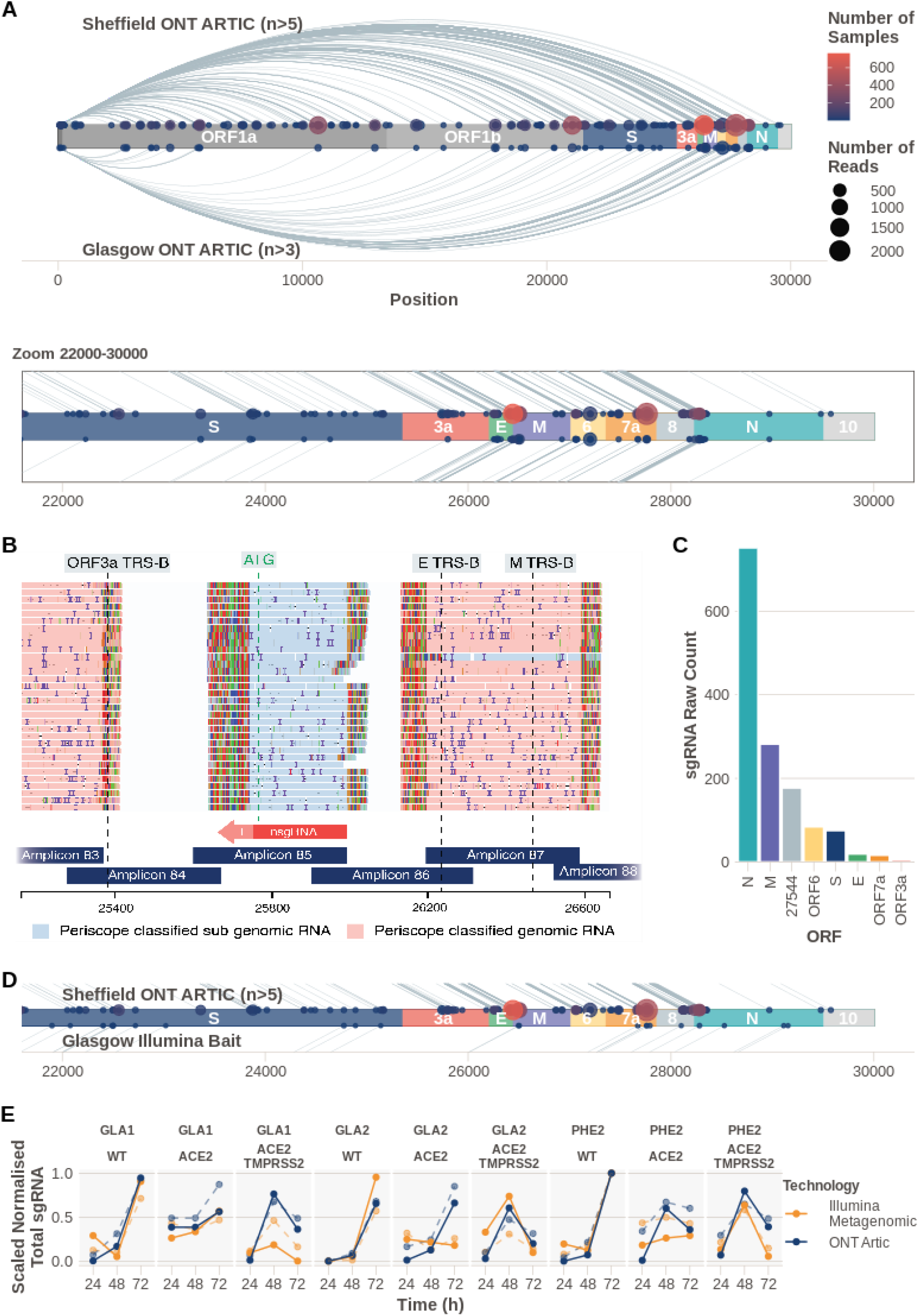
Non-Canonical Sub-Genomic RNA. *We classified reads as supporting non-canonical sgRNA as described in Figure 1D. **A.** Plot showing the number of samples with each non-canonical sgRNA detected in ARTIC Nanopore data. Size of the point represents the number of reads and the color indicates the number of samples that non-canonical sgRNA was found in. Lines connecting points represent the sgRNA product of discontinuous transcription. Those detected in Sheffield samples are above the genome schematic, and Glasgow below. Inset shows zoomed in region between nucleotides 22,000 and 30,000. **B.** Non-canonical sgRNA with strong support in SHEF-C0118 at position 25,744. **C.** Raw sgRNA levels (HQ&LQ) in SHEF-C0118 show high relative amounts of this non-canonical sgRNA at position 25,744. **D.** Zoom of region 22-30kb of the SARS-CoV-2 genome showing non-canonical sgRNA in the Sheffield ONT dataset (top) compared to the non-canonical sgRNA detected in the Illumina bait capture data from Glasgow. **E.** Non-canonical sgRNA levels (solid lines) compared to canonical (dashed lines) in an in vitro model of SARS-CoV-2 infection measured with both Illumina metagenomic sequencing (orange) and ONT Artic (Blue). TotalsgRNA levels are normalised per 100,000 mapped reads and scaled within each dataset for comparison.*

We found evidence of non-canonical sgRNAs supported by two or more reads in 913 samples (Supplementary Figure S6). Unsurprisingly a large number of these recurrent non-canonical sgRNAs cluster around known TRS-B sites (Figure 4A). Interestingly though, we see enrichment of non-canonical sgRNAs at other sites throughout the genome (Figure 4A and Supplementary Figure S6).

In particular, SHEF-C0118 contains 177 reads (HQ & LQ) which support a non-canonical sgRNA at position 25,744 between ORFs 3a and E (Figure 4B&C). The number of reads for this non-canonical sgRNA at 25,744 is high compared to canonical sgRNA from the same sample(Figure 4C, Supplementary Tables S3 & S4). 26 samples contain 1 read supporting this non-canonical sgRNA with 4 samples with > 1 read. In another example, there are 377 samples that have evidence of >=1 read for a non-canonical sgRNA at position 10,639 (HQ or LQ, Supplementary Figure S7A&B; of these 155 have evidence for >= 2 reads +/− 5bp from 10,639). In this case, there is a TRS-*like* sequence close to the leader in this non-canonical sgRNA; ACGAAC -> ACG**G**AC. Two samples have significant support with 103 HQ reads each (SHEF-CE04A, SHEF-CA0D5). It is possible that this represents an independent ORF1b sgRNA. Furthermore, there are 226 samples that have evidence for a non-canonical sgRNA at position 5,785 (>= 1 read; Supplementary Figure S7C&D; 62 with >=2 reads, +/-5 from 5,785), where there is no core TRS sequence present and there does not appear to be a productive start codon.

#### Non-Canonical Sub-Genomic RNAs Across Centres & Technologies

To demonstrate the detection of non-canonical sgRNA was not a phenomenon of the Sheffield dataset and ARTIC Nanopore method alone, we repeated the above analysis on both Nanopore and Illumina data from Glasgow, UK (Figure 4A&D, Supplementary Figure S8E).

The non-canonical sgRNA at 5,785 is found in 15/55 samples from this dataset and 7 of those have multiple read support (Supplementary Table S6). This sgNA is also found with 3 reads in the Illumina sample CVR201. The non-canonical sgRNA found at 10,639 is found in 13/55 samples and 5 of those have multiple read support (Supplementary Table S7). 10,639 is also found in Illumina sample CV196 with 1 read 1 supporting. 25,744 is supported by 1 HQ sgRNA read in 1 sample (CVR2185) from the Glasgow ONT dataset.

To determine if there were any differences in non-canonical sgRNA over time, we applied the same analysis to the *in vitro* system described earlier (Figure 4E & Supplementary Figure S9). Normalised and scaled non-canonical (solid lines) shows that non-canonical sgRNA has comparable kinetics to canonical sgRNA (dashed lines) which appear to be dependent on the virus, cell line and time since infection. This dataset demonstrates that although proportions of non-canonical sgRNA are similar (Supplementary Figure S5), Illumina metagenomic data appears to be more sensitive (Supplementary Figure S5, S9, S10) with a greater number of non-canonical sgRNAs detected.

### Variants in Sub-Genomic RNA

An advantage of having reads from both gRNA and sgRNA is the ability to examine how genomic variants are represented in sgRNAs. For variants found in our isolates by the ARTIC Network nanopolish (Simpson 2018) pipeline we interrogated the bases called (pysam (Gilman et al. 2019) pileup) at the variant position in gRNA and sgRNA to determine if there were detectable differences (this tool is integrated into periscope). As we can only discern gRNA and sgRNA from a small subset of amplicons, the chance that a variant falls within these amplicons is low, but where the two do coincide, variants called in gRNA were supported in reads from sgRNA.

In a small subset of samples we have identified variants in the TRS sequence of some ORFs. Notably 6 samples have a variant at 27,046C>T (ACGAAC to ACGAA**T**)in the TRS of ORF6, this variant is present in both the gRNA and sgRNA reads. 4 of these samples have low expression of ORF6 compared to the rest of the cohort (Figure 5E), although numbers are too low to compute statistical significance.

**Figure 5.**
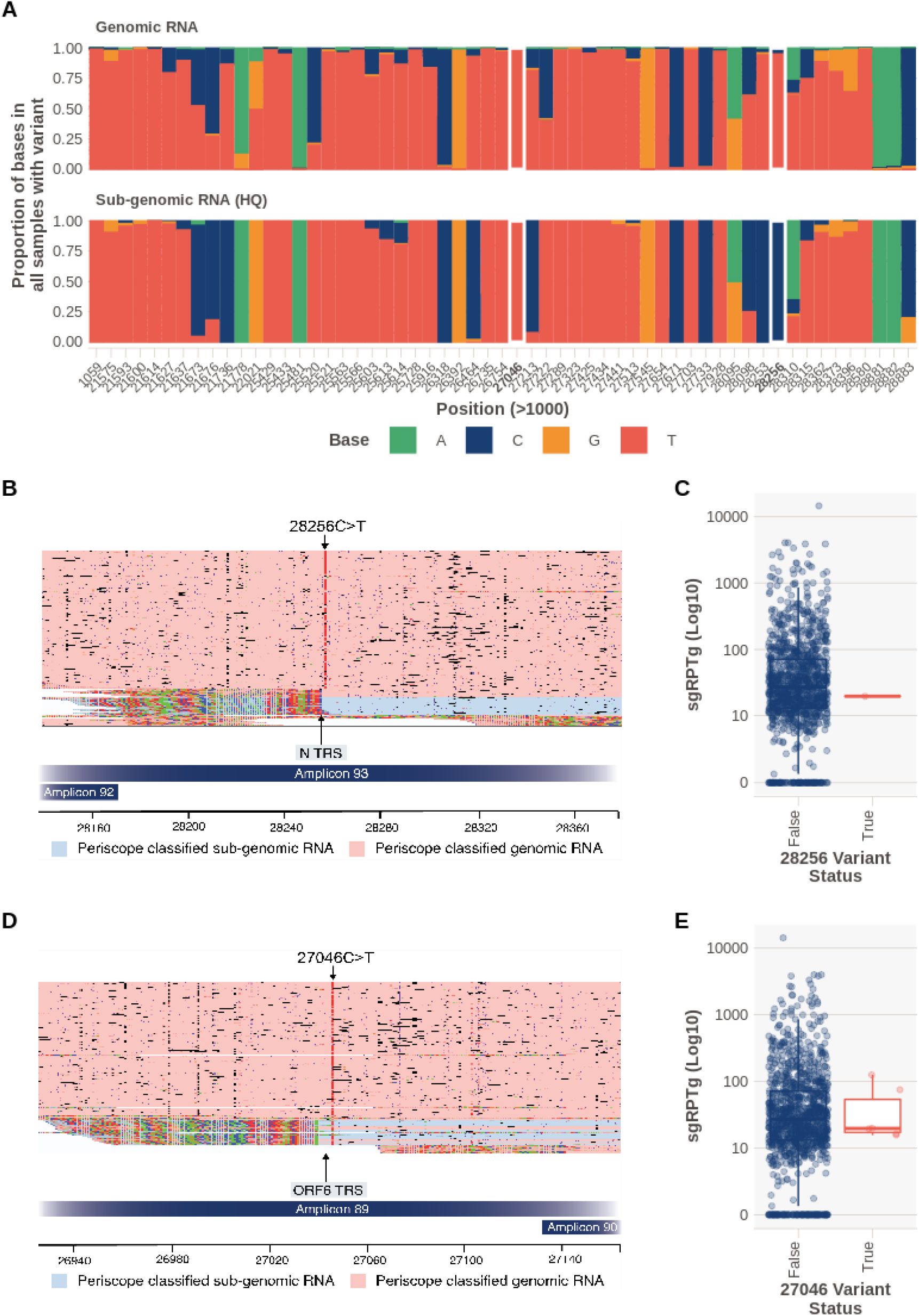
Variants in Sub-Genomic RNA. ***A.** Base frequencies at each of the variant positions called by ARTIC in each sample, (multiple samples can be represented at one position), split by read class. White rectangles represent variants detailed in **(B)** and **(C)**. **B.** SHEF-C0F96 has a 28,256C>T variant, of high quality which sits in the ORF N TRS sequence. This variant is not present in sgreads **C.** Normalised sgRNA expression (sgRNA reads per thousand genomic; sgRPTg) for the N ORF in samples with the variant and without. N expression is one of the lowest in the cohort. **D.** SHEF-C0C35 has 27,046C>T variant of high quality which sits in the TRS sequence. This variant is present in both gRNA and sgRNA. **E.** ORF6 expression levels in samples with 27046C>T.*

In addition, we have one sample with a variant in the N ORF TRS (SHEF-C0F96, 28256C>T. CTAAACGAAC to TTAAACGAAC) which is found only in gRNA but not sgRNA reads. Interestingly, this sample has low expression of most ORFs (ORF N shown in Figure 5C) and has 233,127 reads which is around the median of the cohort. However, this mutation falls outside the core TRS and read counts for sgRNA are low so this result should be treated with caution. It is possible that this represents a sequencing error in the gRNA which is not found in the sgRNA due to the context around that position changing as a result of the inclusion of the leader sequence.

## Discussion

We have developed periscope, a tool that can be used on nearly all publically available SARS-CoV-2 sequence datasets worldwide to detect and quantify sgRNA. Here we applied periscope to 1,155 SARS-CoV-2 sequences from Sheffield, UK and 3 datasets from Glasgow; 55 ARTIC network Nanopore sequences, 5 bait captured Illumina sequenced samples and Illumina metagenomic data from an *in vitro* SARS-CoV-2 infection model. The development of periscope was initially motivated to aid in the the detection of a novel, non-canonical, sgRNA generated (N*) from a de-novo TRS site as a result of the triplet mutation 28,881G>A, 28,882G>A, and 28,883G>C found in a large number of worldwide SARS-CoV-2 isolates (Leary et al. 2020). By searching for reads containing the SARS-CoV-2 leader sequence, incorporated into all sgRNA at their 3’ ends by the SARS-CoV-2 RNA dependent RNA polymerase, we were able to detect sgRNA representing all annotated canonical open reading frames of SARS-CoV-2. ORF10 sgRNA, however, was supported by only two reads in all 1,155 samples in the Sheffield dataset (Supplementary Table S1). Using the *in vitro* SARS-CoV-2 infection system Illumina dataset we identified a further eight reads in total supporting an ORF10 sgRNA. Seven of these reads were present at 72 hours after infection, the remaining read was found at 48 hours. The inability to find significant support for ORF10 mirrors previous findings (Alexandersen, Chamings, and Bhatta 2020; Finkel et al. 2020; Davidson et al. 2020; Kim et al. 2020) and is perhaps unsurprising if ORF10 is indeed non-essential (Pancer et al. 2020). The abundance of other sgRNAs is in line with previously published reports of protein levels in SARS-CoV-2, with M and N showing the highest expression levels after normalisation (Bouhaddou et al. 2020; Finkel et al. 2020). In the Sheffield dataset, the median proportion of total sgRNA is 1.2% (Supplementary Figure S4), which is in broad agreement with published reports, based on ORF E, that sgRNA represented 0.4% of total viral RNA (Wölfel et al. 2020). These findings were mirrored in an equivalent dataset of 55 ONT ARTIC samples from Glasgow. We were able to show that sgRNA analysis using periscope is reproducible, with strong correlations between sgRNA abundance levels between technical replicates.

It has been suggested that sgRNA abundance estimates from amplicon based sequencing data are largely a function of the quality of the RNA in the initial sample, defined in one study by average read length (Alexandersen, Chamings, and Bhatta 2020). The advantage of the ARTIC Network protocol over the AmpliSeq IonTorrent protocol used by the aforementioned study is that the ARTIC Network protocol has a short, consistent amplicon length (mean is 389 and standard deviation is 11.2, Supplementary Figure S1 - gRNA). The assay is designed inherently, to deal with samples with degraded RNA. Since sgRNA reads are a product of these amplicons we do not believe degradation plays a significant role in the determination of abundance levels in our dataset. Furthermore, using E gene CT value as a surrogate for viral load, we find only a weak correlation with the total amount of sgRNA detected (Spearman Rank Correlation, rho=0.268, Supplementary Figure S13A) and this was mainly driven by outliers. Furthermore, sequence coverage across the genome (which is affected by low viral load and poor quality RNA) is also not correlated with sgRNA amount (Spearman Rank Correlation, rho=-0.01188594, Supplementary Figure S13B). Finally on comparison with amplification free approaches like metagenomics we see a strikingly similar pattern of sgRNA expression.

To demonstrate the utility of periscope for the investigation of important biological questions, we applied it to sequencing data from an *in vitro* infection time course using VeroE6 cells which were either wildtype, over-expressing *ACE2* or over-expressing *ACE2* and *TMPRSS2 (Hoffmann et al. 2020)*. These cells were challenged with three different viral isolates (GLA1, GLA2, and PHE2) and total RNA extracted for sequencing by Illumina metagenomics and ONT ARTIC at 24, 48 and 72hrs post infection. The total sgRNA abundance measured using periscope on these orthogonal methods is strikingly similar and reveals an interesting pattern of sgRNA expression. *ACE2* & *TMPRSS2* co-expressing cells have the most obviously altered sgRNA kinetics, with the peak level of sgRNA occuring at 48hrs followed by a reduction at 72hrs, which is in contrast with wildtype cells where sgRNA is still accumulating after 48hrs. This may indicate an expedited course of active replication allowed by greater cellular permissibility with ACE2 & TMPRSS2, followed by attenuated replication in a closed *in vitro* model. Of note, a greater quantity of reads from the Illumina metagenomic data are classified as sgRNA when compared to ONT ARTIC data (Supplementary Figure S4), and there are some differences between the quantities of each ORF when considered individually.

Non-canonical sgRNAs are readily detected by periscope and we present examples where periscope was able to detect high abundances of specific non-canonical sgRNAs in a number of isolates which could indicate some functional significance. It has previously been observed that non-canonical sgRNAs are not formed due to a TRS-like homology (Nomburg, Meyerson, and DeCaprio 2020). The non-canonical sgRNA at position 25,744 in SHEF-C0118 is particularly interesting due to its high relative abundance compared to the canonical sgRNAs in the same sample. There does not appear to be a canonical TRS sequence in close proximity to the leader junction, but there exists a motif which has two mismatches to the canonical TRS; `AAGAAT`. An ATG downstream of the leader in these reads would result in an N-terminal truncated 3a protein. Interestingly, two forms of 3a protein in SARS-CoV have been noted in the literature (Huang et al. 2006). Alternatively, SARS-CoV contains a nested ORF within the 3a sgRNA, 3b (Supplementary Figure S11), but the homolog of this protein is truncated early in SARS-CoV-2, however, others note that a protein from this truncated 3b, of only 22 amino acids in length, could have an immune regulatory function (Konno et al. 2020). This non-canonical sgRNA could indicate production of this novel 3b protein in SARS-CoV-2 independent from the 3a sgRNA, a phenomenon that has been shown to occur in SARS-CoV (Hussain et al. 2005). We cannot explain why, in this sample, this non-canonical sgRNA is present in such high abundance. There are no genomic variants that contribute to a TRS sequence, for example. We also find evidence of highly recurrent non-canonical sgRNAs which have weaker evidence like those at 10,639, which could represent an independent sgRNA for ORF 1b. Others, like those at 5,785 have no apparent related ORF.

Some studies have detected the presence of an sgRNA for ORF7b (Finkel et al. 2020; Kim et al. 2020), we also explored this possibility (Supplementary Figure S11 C,D&E). We are able to detect non-canonical sub-genomic RNAs just 3’ of the predicted start codon of ORF7b (27,760 and 27,761) and at least ten bases downstream of a predicted TRS-B site (Yang et al. 2020). These sgRNAs have string support in both the Sheffield (Supplementary Figure S11D), 668/1155 samples (>=1 high quality sgRNA, median of 2 reads per sample, and a maximum of 133), and Glasgow datasets, 44/55 samples (>=1 high quality sgRNA, median of 1.5 reads per sample, and a maximum of 12). Raw reads from SHEF-BFF12 show that the leader body junction in these sgRNAs does indeed exclude the start codon and in fact includes four additional bases not present in the genome (Supplementary Figure S11E). It is possible that this sgRNA encodes a protein from another ATG site downstream of the leader-body junction.

We were able to detect a number of these recurrent non-canonical sgRNAs in the data from Glasgow, demonstrating that these non-canonical sgRNAs are unlikely to be sequencing artefacts and may represent favoured sites for non-canonical sgRNA generation during SARS-CoV-2 replication for as yet unexplained reasons. We speculate that the diversity of non-canonical sgRNA seen in our dataset, which is most comparable to total RNASeq (Wyler et al. 2020), than direct RNAseq is a function of a) the number of samples analysed, and b) the varied and unknown length of the infection at the time of sampling. These findings illustrate that, although much is not known about the expression of non-canonical sgRNA, periscope could help define and quantify these non-canonical transcripts in order to explore their relevance in SARS-CoV-2 pathogenesis.

We were able to integrate the sgRNAs for variants which were called in gRNA. As expected in most cases, sgRNAs contain the same variants present in the gRNA. We found one case where a variant found in gRNA for ORF N was not present in the ORF N sgRNA. It is possible that this is due to sequencing errors, because the surrounding bases for the variant in gRNA vs sgRNA differ, therefore, base calling could be affected by this change in context.

The COVID-19 Genomics Consortium (COG-UK)(“An Integrated National Scale SARS-CoV-2 Genomic Surveillance Network” 2020) in the UK, alone, has 16,826 ARTIC Nanopore and 69,969 Illumina sequences (Correct 29th October 2020) while internationally GISAID contains thousands more similar datasets (8,775 with “nanopore” in the metadata and 3,660 list “ARTIC” as of 14th June 2020). The application of periscope could therefore provide significant insights into the sgRNA architecture of SARS-CoV-2 at an unprecedented scale. Furthermore, periscope can be provided new primer/amplicon locations for PCR based genomic analysis protocols and we have shown that it can be applied to metagenomic sequencing methods without prior species-specific genome amplification. Periscope could also be applied to sequencing data from other viruses where discontinuous transcription is the method of gene expression.

Periscope offers an opportunity to further understand the regulation of the SARS-CoV-2 genome by identifying and quantifying sgRNA. Applying it to the vast amount of SARS-CoV-2 sequencing datasets that have been generated world wide during this unprecedented public health crisis could uncover critical insights into the role of sgRNA in SARS-CoV-2 pathogenesis.

## Methods

### Sheffield SARS-CoV-2 Sample Collection and Processing

1155 samples from 1155 SARS-CoV-2 positive individuals were obtained from either throat or combined nose/throat swabs. Nucleic acids were extracted from 200μl of each sample using MagnaPure96 extraction platform (Roche Diagnostics Ltd, Burgess Hill, UK). SARS-CoV-2 RNA was detected using primers and probes targeting the E gene and the RdRp genes of SARS-CoV-2 and the human gene RNAseP, to allow normalisation, for routine clinical diagnostic purposes, with thermocycling and fluorescence detection on ABI Thermal Cycler (Applied Biosystems, Foster City, United States) using previously described primer and probe sets (Corman et al. 2020).

### Sheffield SARS-CoV-2 Isolate Amplification and Sequencing

Nucleic acids from positive cases underwent long-read whole genome sequencing (Oxford Nanopore Technologies (ONT), Oxford, UK) using the ARTIC Network protocol (accessed the 19th of April, https://artic.network/ncov-2019, https://www.protocols.io/view/ncov-2019-sequencing-protocol-bbmuik6w). In most cases 23 isolates and one negative control were barcoded per 9.4.1D nanopore flow cell. Following basecalling, data were demultiplexed using ONT Guppy (--require-both-ends). Reads were filtered based on quality and length (400 to 700bp), then mapped to the Wuhan reference genome (MN908947.3) and primer sites trimmed. Reads were then downsampled to 200x coverage in each direction. Variants were called using nanopolish (Simpson 2018).

### *In Vitro* SARS-CoV-2 Infection Model

Two derivatives and an unaltered lineage of the VeroE6 African green monkey kidney cell line were used in this study. One VeroE6 cell line was manipulated to overexpress the angiotensin-converting enzyme 2 (Ace2) receptor. In addition to Ace2 overexpression, a third line was also induced to overexpress the serine protease Transmembrane protease, serine 2 (TMPRSS2). These cell lines are referred to as VeroE6, VeroE6-ACE2 and VeroE6-ACE2-TMPRSS2, respectively. Methods for the production and the validation of these cell lines are fully described in Rhind et al 2020 (accepted PLOS Biology). SARS-CoV-2 infection of these three cell lines were set up with three viral isolates (PHE2, GLA1 and GLA2 - Supplementary Table S9) and supernatant harvested at 24, 48 and 72 hours for viral RNA extraction and sequencing.

### Illumina Metagenomic Sequencing

This protocol was applied to virus isolates propagated *in vitro*. Extracted nucleic acid was incubated with DNAseI (Thermo Fisher, Part Number AM2222) for 5 minutes at 37^°^C. After DNAse treatment, the samples were purified using Agencourt RNA Clean AMPure XP Beads (Beckman Coulter, A63987), following the manufacturer’s guidelines, and quantified using the Qubit dsDNA HS Kit (Thermo Scientific, Part Number Q32854). cDNA was synthesised using SuperScript III (Thermo Scientific, Part Number 18080044) and NEBNext Ultra II Non-Directional RNA Second Strand Synthesis Module (New England Biolabs, Part Number E6111L), as per the manufacturer’s guidelines.

Samples were further processed utilising the Kapa LTP Library Preparation Kit for Illumina Platforms (Kapa Biosystems, Part Number KK8232). Briefly, the cDNA was end repaired and the protocol followed through to adapter ligation. At this stage the samples were uniquely indexed using the NEBNext Multiplex Oligos for Illumina 96 Unique Dual Index Primer Pairs (New England Biolabs, Part Number E6442S), with 15 cycles of PCR performed.

All amplified libraries were quantified by Qubit dsDNA HS Kit and run on the Agilent 4200 Tapestation System (Agilent, Part Number G2991AA) using the High Sensitivity D5000 Screentape (Agilent, Part Number 5067-5592) and High Sensitivity D5000 Reagents (Agilent, Part Number 5067-5593). Libraries were sequenced on an Illumina NextSeq 550 (Illumina, Part Number SY-415-1002).

### Sub-Genomic RNA Detection

Periscope consists of a Python based snakemake(Köster and Rahmann 2012) workflow which runs a python package that processes and classifies reads based on their configuration (Figure 1C).

#### Pre-Processing

##### Nanopore

Pass reads for single isolates are concatenated and aligned to MN908947.3 with minimap2 (v2.17) (Li 2018) (-ax map-ont-k 15). It should be noted that adapters or primers are not trimmed. Bam files are sorted and indexed with samtools(Li et al. 2009).

##### Illumina

Paired end reads, ideally prior to trimming, are aligned to MN908974.3 with bwa mem (v0.7.17) (Li and Durbin 2009) with the “Y” flag set to use soft clipping for supplementary alignments. Bam files are sorted and indexed with samtools.

#### Periscope - Leader Identification & Read Classification

##### Nanopore

Reads from the minimap2 aligned bam file are then processed with pysam(Gilman et al. 2019). If a read is unmapped or represents a supplementary alignment then it is discarded. Each read is then assigned an amplicon using the “find_primer” method of the ARTIC field bioinformatics package. We search for the leader sequence (5’-AACCAACTTTCGATCTCTTGTAGATCTGTTCT-3’) with biopython (Cock et al. 2009) local pairwise alignment (localms) with the following settings, match +2, mismatch −2, gap −10 and extension −0.1 with score_only set to true to speed up computation. The read is then assigned an ORF using a pybedtools(Dale, Pedersen, and Quinlan 2011) and a bed file consisting of all known ORFs +/-10 of the predicted leader/genome transition.

We classify reads as a “High Quality” (HQ) sgRNA (Figure 1D) if the alignment score is > 50 and the read is at a known ORF. If the read starts at a primer site then it is classified as gRNA, if not, then it is classified as a “High Quality” non-canonical sgRNA supporting read. If the alignment score is > 30 but <= 50 and the read is at a known ORF then it is classified as a “Low Quality” (LQ) sgRNA. If the read is within a primer site, it is labelled as a gRNA, if not, then it is a “Low Quality” sgRNA. Finally, any reads with a score of <= 30 and are at a known ORF are then classified as a “Low Low Quality” sgRNA, otherwise, they are labelled as gRNA. The following tags are added to the reads for manual review of the periscope calls; XS: Alignment score, XA: Amplicon, AC: Read class, and XO: The read ORF. Reads are binned into qualitative categories (HQ, LQ, LLQ etc) because we noticed that some sgRNAs were not classified as such due to a lower match to the leader. After manual review, they are deemed bona fide sgRNA. This quality rating negates the need to alter alignment score cut-offs continually to find the best balance between sensitivity and specificity. Restricting to HQ data means that sensitivity is reduced but specificity is increased, including LQ calls will decrease specificity but increase sensitivity.

##### Illumina

Reads from the bwa mem aligned bam file are processed with pysam. If a read is unmapped or represents a supplementary alignment then it is discarded.

The presence of soft clipping at the 5’ end of the reads is an indicator that that read could contain the leader sequence so we extract all of the soft clipped bases from the 5’ end, additionally including three further bases to account for homology between leader and genome at the N ORF (these bases would therefore not be soft clipped at this ORF). If there are less than six extracted bases in total we do not process that read further as this is not enough to determine a robust match to the leader sequence. With a match score of 2 and a mismatch score of −2 (Gap opening penalty; −20, Extension, −0.1) soft clipped bases are aligned to the leader sequence with localms. Soft clipped bases that include the full >=33 bp of the leader would give an alignment score of 66. Allowing for two mismatches this gives a “perfect” score of 60. If the number of soft clipped bases is less then we adjust the “perfect” score in the following way:

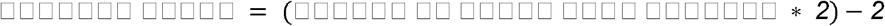

This allows for one mismatch. The position of the alignment is then checked, for these bases to be classed as the leader, the alignment of the soft clipped bases must be at the 3’ end. If this is true and: 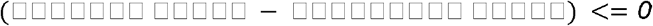 then the read is classified as sgRNA.

#### Periscope - Sub-genomic RNA Normalisation

##### Amplicon Data

Once reads have been classified, the counts are summarised and normalised. Two normalisation schemes are employed:

1. **Normalisation to Total Mapped Reads** Total mapped reads per sample are calculated using pysam idxstats and used to normalise both genomic, sgRNA and non-canonical sgRNA reads. Reads per hundred thousand total mapped reads are calculated per quality group.

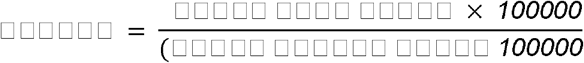
2. **Normalisation to Genomic Reads from the Corresponding Amplicon** Counts of sgRNA, non-canonical sgRNA, and gRNA are recorded on a per amplicon basis and normalisation occurs within the same amplicon per 1000 gRNA reads. If multiple amplicons contribute to the count of sgRNA or non-canonical sgRNA then the normalised values are summed.

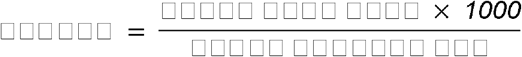

Periscope outputs several useful files which are described in more detail in the Supplementary Material, briefly these are; periscope’s processed bam file with associated tags, a per amplicons counts file, and a summarised counts file for both canonical and non-canonical ORFs.

##### Bait Based Capture & Metagenomic Data

Again two schemes are employed, as above, to the total amount of mapped reads and additionally:

**3. Normalisation to Local Coverage** For both canonical and non-canonical sgRNA normalisation we calculate the median coverage around either; a) for canonical ORFs the TRS site +/− 20bp or b) the leader/genome junction +/− 20bp and normalise the total sgRNA per 1000 reads of coverage.

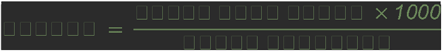

### Sub-Genomic Variant Analysis

A python script is provided “variant_expression.py” that takes the periscope bam file and a VCF file of variants (usually from the ARTIC analysis pipeline). For each position in the VCF (pyvcf(Casbon 2012)) file, it extracts the counts of each base in each class of read (i.e. genomic, sgRNA and non-canonical sgRNA) and outputs these counts as a table. This tool also provides a useful plot (Supplementary Figure S5) of the base counts at each position for each class.

### Analysis and Figure Generation

Further analysis was completed in R 3.5.2(Schulte et al. 2012) using Rstudio 1.1.442(Racine 2012), in general data was processed using dplyr (v0.8.3), figures were generated using ggplot2 (v3.3.1), both part of the tidyverse(Wickham et al. 2019) family of packages (v1.2.1). Plots themed with the ftplottools package (v0.1.5). GGally (v2.0.0) ggpairs was used for the matrix plots for downsampling and repeats. Where multiple hypothesis tests were carried out, multiple testing correction was carried out using Bonferroni. Reads were visualised in IGV(Robinson et al. 2011) and annotated with Adobe Illustrator. Code for figure generation can be found attached as a supplementary file and in the github repository: https://github.com/sheffield-bioinformatics-core/periscope-publication.

#### Principal Component Analysis (PCA)

PCA was carried out to determine if any of the experimental variables were responsible for the differences in expression values between samples. Reads from ORF1a and ORF10 amplicons were removed from the analysis and expression values were normalised within each ORF:

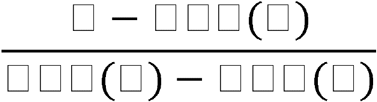

and then row means subtracted. The “PCA” function of the R package FactoMineR(Pagès 2014), (v1.41) was then used to carry out the PCA without further scaling and all other settings as default. The resulting PCA was plotted using the “fviz_pca_ind” function. Plots were coloured according to the variable in question.

### Data Cleaning

Data was uploaded to ENA after removing human reads with dehumaizer (https://github.com/SamStudio8/dehumanizer)

### Periscope Requirements

Periscope is a wrapper for a snakemake(Köster and Rahmann 2012) workflow with a package written in python to implement read filtering and classification, and is provided with a conda environment definition. It has been tested on a Dell XPS, core i9, 32gb ram, 1Tb SSD running ubuntu 18.04 and was able to process ~100,000 reads per minute. Periscope installation requires conda (For example Miniconda, https://docs.conda.io/en/latest/miniconda.html, version 3.7 or 3.8). To run periscope, you will need the path to your raw fastq files from your ARTIC Network Nanopore sequencing run or Illumina paired end reads (unfiltered), and other variables defined in the Supplementary Material and on the github README.

### Data Access

All raw sequencing data generated in this study have been submitted to the European Nucleotide Archive under study ID: PRJEB40972.

Periscope is freely available under GNU General Public License v3.0 from the Sheffield Bioinformatics Core github account (https://github.com/sheffield-bioinformatics-core/periscope) and the source code is contained within Supplementary File 18 (supplementary_file_18_periscope-0.0.8a.tar.gz)

The data and code required to recreate the analysis contained within this manuscript can be be found on github (https://github.com/sheffield-bioinformatics-core/periscope-publication) as well as supplementary files published along with this manuscript; Supplementary files 1-16 and Supplementary FIle 17 (supplementary_file_17_figure_generation.Rmd).

## Ethics Approval and Consent

Individuals presenting with active COVID-19 disease were sampled for SARS CoV-2 sequencing at Sheffield Teaching Hospitals NHS Foundation Trust, UK using samples collected for routine clinical diagnostic use. This work was performed under approval by the Public Health England Research Ethics and Governance Group for the COVID-19 Genomics UK consortium (R&D NR0195).

## Acknowledgements

We would like to thank the Mark Dunning of the Sheffield Bioinformatics Core for useful discussions around the normalisation techniques. We thank all partners of and contributors to the COG-UK consortium, who are listed at https://www.cogconsortium.uk/about/ and in the Supplementary Material.

## Author Contributions

MDP developed periscope, analysed and interpreted data and was a major contributor to manuscript preparation. BL analysed and interpreted data and contributed to manuscript preparation. DW interpreted data and contributed to manuscript preparation. SL,SG, SM, AC originally conceived the method for sgRNA detection and contributed to manuscript preparation. MW, LRG, PP, DG, RT, RB, LC, AA, AK, KJ and NS collated, processed and sequenced samples, and reviewed the manuscript. JH is a developer of our LIMS system to ensure efficient sample processing, and reviewed the manuscript. MR, CE and DGP collected and organised clinical samples and metadata and reviewed the manuscript. AdSF, JS, CD, AK, EAC, LT, JN, ECT generated SARS-CoV-2 sequencing data from clinical isolates and cell lines in Glasgow, and contributed to manuscript preparation and data interpretation. TdS conceived the study and contributed to manuscript preparation and oversight. All authors read and approved the final manuscript.

## Disclosure Declaration

All authors declare there are no conflicts of interest.

## Funding

Sequencing of SARS-CoV-2 samples was undertaken by the Sheffield COVID-19 Genomics Group as part of the COG-UK CONSORTIUM and supported by funding from the Medical Research Council (MRC) part of UK Research & Innovation (UKRI), the National Institute of Health Research (NIHR) and Genome Research Limited, operating as the Wellcome Sanger Institute. MDP and DW were funded by the NIHR Sheffield Biomedical Research Centre (BRC - IS-BRC-1215-20017). TIdS is supported by a Wellcome Trust Intermediate Clinical Fellowship (110058/Z/15/Z). AK is supported by the MRC (MC_UU_12014/8). AdSF and LT are supported by the MRC (MC_UU_12014/12).

## Notes

https://github.com/sheffield-bioinformatics-core/periscope

https://github.com/sheffield-bioinformatics-core/periscope-publication

### Competing Interest Statement

The authors have declared no competing interest.

### Summary of Updates

There has been significant additions to the manuscript: * Addition of 55 ARTIC Nanopore sequences from Glasgow * Addition of in vitro experiments comparing sub-genomic RNA measurement with both ARTIC Nanopore and Illumina metagenomics from Glasgow * Illumina bait capture from Glasgow Other changes include more examination of non-canonical sgRNA and particularly around ORF3b and ORF7b, updates to figures, addition of all summary tables and code for figure generation.

