## Supplementary Material for "periscope: sub-genomic RNA identification in SARS-CoV-2 Genomic Sequencing Data"

#### Contents

**COG-UK Consortium Author List** 3
**Supplementary Files** **11**
**Periscope README** **14**
Requirements 14
Installation 14
Execution 14
Output Files 15
Examining Base Frequencies of Called Variants in periscope 16
**Supplementary Methods** **17**
Glasgow Oxford Nanopore Amplicon Sequencing 17
Glasgow Bait Capture & Subsequent Illumina Sequencing 18
**Supplementary Tables** **20**
Supplementary Table S1 - sgRNAs detected for each canonical ORF 20
Supplementary Table S2 - Samples with predicted sgRNA for ORF10 21
Supplementary Table S3 - Non-canonical sgRNA at position 25,744 in sample SHEF-C0118 has strong support 22
Supplementary Table S4 - Canonical sgRNA in SHEF-C0118 23
Supplementary Table S5 - Non-canonical sgRNA at 10,639 in SHEF-CE04A has strong support 23
Supplementary Table S6 - Non-canonical sgRNA at 5,785 in Glasgow Nanopore samples 24
Supplementary Table S7 - Non-canonical sgRNA at 10,639 in Glasgow Nanopore samples 25
Supplementary Table S8 - Canonical sub-genomic RNA in SHEF-CE04A 26
Supplementary Table S9 - Changes with respect to MN908947.3 in viral isolates used in in vitro SARS-CoV-2 infection model 27
**Supplementary Figures** **28**

Supplementary Figure S1 - Length of reads in SHEF ONT ARTIC data broken down by periscope assigned class in a representative sample (SHEF-BFDBE) 28
Supplementary Figure S2 - SARS-CoV-2 Illumina Data 29
Supplementary Figure S3 - Manual review of periscope bam files for ORF10 31
Supplementary Figure S4 - Mapped read counts 32
Supplementary Figure S5 - Sub-Genomic RNA Proportions 33
Supplementary Figure S6 - Number of samples with at the most frequently represented non-canonical sub-genomic RNAs 34
Supplementary Figure S7 - Highly expressed non-canonical sgRNAs at 10,369 and 5,785 35
Supplementary Figure S8 - sgRNA Levels in bait capture Illumina samples (n=5) 37
Supplementary Figure S9 - Non-canonical sub-genomic RNA detected in an vitro infection model 39
Supplementary Figure S10 - Read level detail of Non-canonical sub-genomic RNA in Illumina metagenomic sequencing of an in vitro infection model 40
Supplementary Figure S11 - Non-canonical sub-genomic RNAs (High quality) that could represent ORF3b and ORF7b 41
Supplementary Figure S12 - Example of periscope output for variant analysis 43
Supplementary Figure S13 - Normalised E Ct and consensus coverage are not correlated with the raw amount of sub-genomic RNA detected 44

### COG-UK Consortium Author List

**Funding acquisition, leadership, supervision, metadata curation, project administration, samples, logistics, Sequencing, analysis, and Software and analysis tools:**

Thomas R Connor [33, 34], and Nicholas J Loman [15].

**Leadership, supervision, sequencing, analysis, funding acquisition, metadata curation, project administration, samples, logistics, and visualisation:**

Samuel C Robson [68].

**Leadership, supervision, project administration, visualisation, samples, logistics, metadata curation and software and analysis tools:**

Tanya Golubchik [27].

**Leadership, supervision, metadata curation, project administration, samples, logistics sequencing and analysis:**

M. Estee Torok [8, 10].

**Project administration, metadata curation, samples, logistics, sequencing, analysis, and software and analysis tools:**

William L Hamilton [8, 10].

**Leadership, supervision, samples logistics, project administration, funding acquisition sequencing and analysis:**

David Bonsall [27].

**Leadership and supervision, sequencing, analysis, funding acquisition, visualisation and software and analysis tools:**

Ali R Awan [74].

**Leadership and supervision, funding acquisition, sequencing, analysis, metadata curation, samples and logistics:**

Sally Corden[33] .

**Leadership supervision, sequencing analysis, samples, logistics, and metadata curation:** Ian Goodfellow [11].

**Leadership, supervision, sequencing, analysis, samples, logistics, and Project administration:**

Darren L Smith [60, 61].

**Project administration, metadata curation, samples, logistics, sequencing and analysis:**

Martin D Curran [14], and Surendra Parmar [14].

**Samples, logistics, metadata curation, project administration sequencing and analysis:**

James G Shepherd [21].

**Sequencing, analysis, project administration, metadata curation and software and analysis tools:**

Matthew D Parker [38,] and Dinesh Aggarwal [1, 2, 3].

**Leadership, supervision, funding acquisition, samples, logistics, and metadata curation:**

Catherine Moore [33].

**Leadership, supervision, metadata curation, samples, logistics, sequencing and analysis:**

Derek J Fairley[6, 88], Matthew W Loose [54], and Joanne Watkins [33].

**Metadata curation, sequencing, analysis, leadership, supervision and software and analysis tools:**

Matthew Bull [33] , and Sam Nicholls [15].

**Leadership, supervision, visualisation, sequencing, analysis and software and analysis tools:**

David M Aanensen [1, 30].

**Sequencing, analysis, samples, logistics, metadata curation, and visualisation:**

Sharon Glaysher [70] .

**Metadata curation, sequencing, analysis, visualisation, software and analysis tools:**

Matthew Bashton [60], and Nicole Pacchiarini [33].

**Sequencing, analysis, visualisation, metadata curation, and software and analysis tools**:

Anthony P Underwood[1, 30].

**Funding acquisition, leadership, supervision and project administration:**

Thushan I de Silva[38], and Dennis Wang [38].

**Project administration, samples, logistics, leadership and supervision**:

Monique Andersson[28] , Anoop J Chauhan [70], Mariateresa de Cesare [26], Catherine Ludden [1,3] , and Tabitha W Mahungu [91].

**Sequencing, analysis, project administration and metadata curation:**

Rebecca Dewar [20], and Martin P McHugh [20].

**Samples, logistics, metadata curation and project administration:**

Natasha G Jesudason [21], Kathy K Li MBBCh [21], Rajiv N Shah [21], and Yusri Taha [66].

**Leadership, supervision, funding acquisition and metadata curation:**

Kate E Templeton [20]**.**

**Leadership, supervision, funding acquisition, sequencing and analysis:**

Simon Cottrell [33], Justin O'Grady [51], Andrew Rambaut [19], and Colin P Smith[93].

**Leadership, supervision, metadata curation , sequencing and analysis:**

Matthew T.G. Holden [87], and Emma C Thomson [21].

**Leadership, supervision, samples, logistics and metadata curation**:

Samuel Moses [81, 82].

**Sequencing, analysis, leadership, supervision, samples and logistics:**

Meera Chand [7], Chrystala Constantinidou [71], Alistair C Darby [46], Julian A Hiscox [46], Steve Paterson [46], and Meera Unnikrishnan [71]**.**

**Sequencing, analysis, leadership and supervision and software and analysis tools:**

Andrew J Page [51], and Erik M Volz [96].

**Samples, logistics, sequencing, analysis and metadata curation:**

Charlotte J Houldcroft [8], Aminu S Jahun [11], James P McKenna [88], Luke W Meredith [11], Andrew Nelson [61], Sarojini Pandey [72], and Gregory R Young [60].

**Sequencing, analysis, metadata curation, and software and analysis tools:**

Anna Price [34], Sara Rey [33], Sunando Roy [41], Ben Temperton[49], and Matthew Wyles [38].

**Sequencing, analysis, metadata curation and visualisation:**

Stefan Rooke[19], and Sharif Shaaban [87].

**Visualisation, sequencing, analysis and software and analysis tools:**

Helen Adams [35], Yann Bourgeois [69], Katie F Loveson [68], Áine O'Toole [19], and Richard Stark [71].

**Project administration, leadership and supervision:**

Ewan M Harrison [1, 3], David Heyburn [33], and Sharon J Peacock [2, 3]

**Project administration and funding acquisition:**

David Buck [26], and Michaela John[36]

**Sequencing, analysis and project administration:**

Dorota Jamrozy [1], and Joshua Quick [15]

**Samples, logistics, and project administration:**

Rahul Batra [78], Katherine L Bellis [1, 3], Beth Blane [3], Sophia T Girgis [3], Angie Green [26], Anita Justice [28], Mark Kristiansen [41], and Rachel J Williams [41].

**Project administration, software and analysis tools:**

Radoslaw Poplawski[15].

**Project administration and visualisation:**

Garry P Scarlett [69].

**Leadership, supervision, and funding acquisition:**

John A Todd [26], Christophe Fraser [27], Judith Breuer [40,41], Sergi Castellano [41], Stephen L Michell [49], Dimitris Gramatopoulos [73], and Jonathan Edgeworth [78].

**Leadership, supervision and metadata curation:**

Gemma L Kay [51].

**Leadership, supervision, sequencing and analysis:**

Ana da Silva Filipe [21], Aaron R Jeffries [49], Sascha Ott [71], Oliver Pybus [24], David L Robertson [21], David A Simpson [6], and Chris Williams [33].

**Samples, logistics, leadership and supervision:**

Cressida Auckland [50], John Boyes [83], Samir Dervisevic [52], Sian Ellard [49, 50], Sonia Goncalves[1], Emma J Meader [51], Peter Muir [2], Husam Osman [95], Reenesh Prakash [52], Venkat Sivaprakasam [18], and Ian B Vipond [2].

**Leadership, supervision and visualisation**

Jane AH Masoli [49, 50].

**Sequencing, analysis and metadata curation**

Nabil-Fareed Alikhan [51], Matthew Carlile [54], Noel Craine [33], Sam T Haldenby [46], Nadine Holmes [54], Ronan A Lyons [37], Christopher Moore [54], Malorie Perry [33] , Ben Warne [80], and Thomas Williams [19].

**Samples, logistics and metadata curation:**

Lisa Berry [72], Andrew Bosworth [95] , Julianne Rose Brown [40], Sharon Campbell [67], Anna Casey [17], Gemma Clark [56], Jennifer Collins [66], Alison Cox [43, 44] , Thomas Davis [84], Gary Eltringham [66], Cariad Evans [38, 39] , Clive Graham [64], Fenella Halstead [18], Kathryn Ann Harris [40], Christopher Holmes [58], Stephanie Hutchings [2] , Miren Iturriza-Gomara [46], Kate Johnson [38, 39], Katie Jones [72], Alexander J Keeley [38], Bridget A Knight [49, 50], Cherian Koshy[90], Steven Liggett [63], Hannah Lowe [81] , Anita O Lucaci [46] , Jessica Lynch [25, 29] , Patrick C McClure [55], Nathan Moore [31] , Matilde Mori [25, 29, 32] , David G Partridge [38, 39] , Pinglawathee Madona [43, 44] , Hannah M Pymont [2], Paul Anthony Randell [43, 44] , Mohammad Raza [38, 39] , Felicity Ryan [81] , Robert Shaw [28], Tim J Sloan [57], and Emma Swindells [65] .

**Sequencing, analysis, Samples and logistics:**

Alexander Adams [33], Hibo Asad [33], Alec Birchley [33] , Tony Thomas Brooks [41], Giselda Bucca [93], Ethan Butcher [70], Sarah L Caddy [13], Laura G Caller [2, 3, 12] , Yasmin Chaudhry [11], Jason Coombes [33], Michelle Cronin [33], Patricia L Dyal [41], Johnathan M Evans [33], Laia Fina [33], Bree Gatica-Wilcox [33], Iliana Georgana [11], Lauren Gilbert [33] , Lee Graham [33], Danielle C Groves [38], Grant Hall [11], Ember Hilvers [33], Myra Hosmillo [11], Hannah Jones [33], Sophie Jones [33], Fahad A Khokhar [13] , Sara Kumziene-Summerhayes [33], George MacIntyre-Cockett [26], Rocio T Martinez Nunez [94], Caoimhe McKerr [33], Claire McMurray [15], Richard Myers [7], Yasmin Nicole Panchbhaya [41], Malte L Pinckert [11] , Amy Plimmer [33] , Joanne Stockton [15] , Sarah Taylor [33] , Alicia Thornton [7] , Amy Trebes [26] , Alexander J Trotter [51] ,Helena Jane Tutill [41] ,Charlotte A Williams [41] , Anna Yakovleva [11] and Wen C Yew [62].

**Sequencing, analysis and software and analysis tools:**

Mohammad T Alam [71], Laura Baxter [71], Olivia Boyd [96] , Fabricia F. Nascimento [96], Timothy M Freeman [38], Lily Geidelberg [96], Joseph Hughes [21], David Jorgensen [96], Benjamin B Lindsey [38], Richard J Orton [21] , Manon Ragonnet-Cronin [96] Joel Southgate [33, 34,] and Sreenu Vattipally [21].

**Samples, logistics and software and analysis tools:**

Igor Starinskij [23].

**Visualisation and software and analysis tools:**

Joshua B Singer [21] , Khalil Abudahab [1, 30], Leonardo de Oliveira Martins [51] , Thanh Le-Viet [51] ,Mirko Menegazzo [30] ,Ben EW Taylor [1, 30], and Corin A Yeats [30].

**Project Administration:**

Sophie Palmer [3], Carol M Churcher [3] , Alisha Davies [33], Elen De Lacy [33], Fatima Downing [33], Sue Edwards [33] , Nikki Smith [38] , Francesc Coll [97] , Nazreen F Hadjirin [3] and Frances Bolt [44, 45].

**Leadership and supervision:**

Alex Alderton[1], Matt Berriman[1], Ian G Charles [51], Nicholas Cortes [31], Tanya Curran [88], John Danesh[1], Sahar Eldirdiri [84], Ngozi Elumogo [52], Andrew Hattersley [49, 50], Alison Holmes [44, 45], Robin Howe [33], Rachel Jones [33], Anita Kenyon [84], Robert A Kingsley [51], Dominic Kwiatkowski [1, 9], Cordelia Langford[1], Jenifer Mason[48], Alison E Mather [51], Lizzie Meadows [51], Sian Morgan [36], James Price [44, 45], Trevor I Robinson [48], Giri Shankar [33] , John Wain [51], and Mark A Webber [51].

**Metadata curation:**

Declan T Bradley [5, 6], Michael R Chapman [1, 3, 4] , Derrick Crooke [28] , David Eyre [28], Martyn Guest [34] , Huw Gulliver [34], Sarah Hoosdally [28], Christine Kitchen [34], Ian Merrick [34], Siddharth Mookerjee [44, 45], Robert Munn [34] , Timothy Peto [28], Will Potter [52], Dheeraj K Sethi [52], Wendy Smith [56] , Luke B Snell [75, 94], Rachael Stanley [52] , Claire Stuart [52] and Elizabeth Wastenge[20].

**Sequencing and analysis:**

Erwan Acheson [6] , Safiah Afifi [36] , Elias Allara [2, 3] , Roberto Amato [1], Adrienn Angyal [38], Elihu Aranday-Cortes [21] , Cristina Ariani [1], Jordan Ashworth [19], Stephen Attwood [24], Alp Aydin [51], David J Baker [51], Carlos E Balcazar [19], Angela Beckett [68] Robert Beer [36], Gilberto Betancor [76], Emma Betteridge [1] , David Bibby [7] , Daniel Bradshaw[7] , Catherine Bresner [34], Hannah E Bridgewater [71] , Alice Broos [21], Rebecca Brown [38] , Paul E Brown [71], Kirstyn Brunker [22] , Stephen N Carmichael [21] , Jeffrey K. J. Cheng [71], Dr Rachel Colquhoun [19], Gavin Dabrera [7] , Johnny Debebe [54], Eleanor Drury [1], Louis du Plessis [24] , Richard Eccles [46], Nicholas Ellaby [7], Audrey Farbos [49], Ben Farr [1], Jacqueline Findlay [41] , Chloe L Fisher [74], Leysa Marie Forrest [41], Sarah Francois [24], Lucy R. Frost [71], William Fuller[34] , Eileen Gallagher [7], Michael D Gallagher [19] , Matthew Gemmell [46], Rachel AJ Gilroy [51], Scott Goodwin [1], Luke R Green [38], Richard Gregory [46] , Natalie Groves [7], James W Harrison [49], Hassan Hartman [7] , Andrew R Hesketh [93],Verity Hill [19], Jonathan Hubb [7], Margaret Hughes[46] , David K Jackson [1] , Ben Jackson [19], Keith James [1] ,Natasha Johnson [21] ,Ian Johnston [1], Jon-Paul Keatley [1], Moritz Kraemer [24], Angie Lackenby [7], Mara Lawniczak [1] , David Lee [7], Rich Livett [1], Stephanie Lo [1], Daniel Mair [21], Joshua Maksimovic [36], Nikos Manesis [7] , Robin Manley [49], Carmen Manso [7], Angela Marchbank [34] , Inigo Martincorena [1] , Tamyo Mbisa [7], Kathryn McCluggage [36], JT McCrone [19], Shahjahan Miah [7] , Michelle L Michelsen [49], Mari Morgan [33], Gaia Nebbia [78],Charlotte Nelson [46] ,Jenna Nichols [21] ,Paola Niola [41] , Kyriaki Nomikou [21] ,Steve Palmer [1] , Naomi Park [1], Yasmin A Parr [1] , Paul J Parsons [38] , Vineet Patel [7] , Minal Patel [1] ,Clare Pearson [2, 1], Steven Platt [7] ,Christoph Puethe [1], Mike Quail [1],Jayna Raghwani [24] , Lucille Rainbow [46] ,Shavanthi Rajatileka [1], Mary Ramsay [7] , Paola C Resende Silva [41, 42], Steven Rudder 51, Chris Ruis [3] , Christine M Sambles [49], Fei Sang [54], Ulf Schaefer[7], Emily Scher [19], Carol Scott [1] ,Lesley Shirley [1], Adrian W Signell [76], John Sillitoe [1] ,Christen Smith [1] ,Dr Katherine L Smollett [21] ,Karla Spellman [36] ,Thomas D Stanton [19], David J Studholme [49] ,Grace Taylor-Joyce [71] ,Ana P Tedim [51], Thomas Thompson [6], Nicholas M Thomson [51], Scott Thurston[1] , Lily Tong [21], Gerry Tonkin-Hill [1], Rachel M Tucker [38] , Edith E Vamos [4], Tetyana Vasylyeva[24], Joanna Warwick-Dugdale [49] , Danni Weldon [1], Mark Whitehead [46], David Williams [7], Kathleen A Williamson [19],Harry D Wilson [76],Trudy Workman [34], Muhammad Yasir[51], Xiaoyu Yu [19], and Alex Zarebski [24].

**Samples and logistics:**

Evelien M Adriaenssens [51], Shazaad S Y Ahmad [2, 47] , Adela Alcolea-Medina [59, 77], John Allan [60], Patawee Asamaphan [21], Laura Atkinson [40], Paul Baker [63], Jonathan Ball [55], Edward Barton[64], Mathew A Beale[1], Charlotte Beaver[1], Andrew Beggs [16], Andrew Bell [51], Duncan J

Berger [1], Louise Berry. [56], Claire M Bewshea [49], Kelly Bicknell [70], Paul Bird [58], Chloe Bishop [7], Tim Boswell [56], Cassie Breen [48], Sarah K Buddenborg[1], Shirelle Burton-Fanning [66], Vicki Chalker [7], Joseph G Chappell [55], Themoula Charalampous [78, 94], Claire Cormie[3], Nick Cortes[29, 25], Lindsay J Coupland [52], Angela Cowell [48], Rose K Davidson [53], Joana Dias [3], Maria Diaz [51], Thomas Dibling[1], Matthew J Dorman[1], Nichola Duckworth[57], Scott Elliott[70], Sarah Essex[63], Karlie Fallon [58], Theresa Feltwell [8], Vicki M Fleming [56], Sally Forrest [3], Luke Foulser[1], Maria V Garcia-Casado[1], Artemis Gavriil [41], Ryan P George [47], Laura Gifford [33], Harmeet K Gill [3], Jane Greenaway [65], Luke Griffith[53], Ana Victoria Gutierrez[51], Antony D Hale [85], Tanzina Haque [91], Katherine L Harper [85], Ian Harrison [7], Judith Heaney [89], Thomas Helmer [58], Ellen E Higginson[3], Richard Hopes [2], Hannah C Howson-Wells [56], Adam D Hunter [1], Robert Impey [70], Dianne Irish-Tavares [91], David A Jackson[1], Kathryn A Jackson [46], Amelia Joseph [56], Leanne Kane [1], Sally Kay [1], Leanne M Kermack [3], Manjinder Khakh [56], Stephen P Kidd [29, 25,31], Anastasia Kolyva [51], Jack CD Lee [40], Laura Letchford [1], Nick Levene [79], Lisa J Levett [89], Michelle M Lister [56], Allyson Lloyd [70], Joshua Loh [60], Louissa R Macfarlane-Smith [85], Nicholas W Machin [2, 47], Mailis Maes [3], Samantha McGuigan [1], Liz McMinn [1], Lamia Mestek-Boukhibar [41], Zoltan Molnar [6], Lynn Monaghan [79], Catrin Moore [27], Plamena Naydenova [3], Alexandra S Neaverson [1], Rachel Nelson [1], Marc O Niebel [21], Elaine O'Toole[48], Debra Padgett [64], Gaurang Patel [1], Brendan AI Payne [66], Liam Prestwood [1], Veena Raviprakash [67], Nicola Reynolds[86], Alex Richter [16], Esther Robinson [95], Hazel A Rogers[1], Aileen Rowan [96], Garren Scott [64], Divya Shah [40], Nicola Sheriff [67], Graciela Sluga, Emily Souster[1], Michael Spencer-Chapman[1], Sushmita Sridhar [1, 3], Tracey Swingler [53], Julian Tang[58], Graham P Taylor[96], Theocharis Tsoleridis [55], Lance Turtle[46], Sarah Walsh [57], Michelle Wantoch [86], Joanne Watts [48], Sheila Waugh [66], Sam Weeks[41], Rebecca Williams[31], Iona Willingham[56], Emma L Wise [25, 29, 31], Victoria Wright [54], Sarah Wyllie [70], and Jamie Young [3].

**Software and analysis tools:**

Amy Gaskin[33], Will Rowe [15], and Igor Siveroni [96].

**Visualisation:**

Robert Johnson [96].

**Affiliations:**

**1** Wellcome Sanger Institute, **2** Public Health England, **3** University of Cambridge, **4** Health Data Research UK, Cambridge, **5** Public Health Agency, Northern Ireland ,**6** Queen's University Belfast **7** Public Health England Colindale, **8** Department of Medicine, University of Cambridge, **9** University of Oxford, **10** Departments of Infectious Diseases and Microbiology, Cambridge University Hospitals NHS Foundation Trust; Cambridge, UK, **11** Division of Virology, Department of Pathology, University of Cambridge, **12** The Francis Crick Institute, **13** Cambridge Institute for Therapeutic Immunology and Infectious Disease, Department of Medicine, **14** Public Health England, Clinical Microbiology and Public Health Laboratory, Cambridge, UK, **15** Institute of Microbiology and Infection, University of Birmingham, **16** University of Birmingham, **17** Queen Elizabeth Hospital, **18** Heartlands Hospital, **19** University of Edinburgh, **20** NHS Lothian, **21** MRC-University of Glasgow Centre for Virus Research, **22** Institute of Biodiversity, Animal Health & Comparative Medicine, University of Glasgow, **23** West of Scotland Specialist Virology Centre, **24** Dept Zoology, University of Oxford, **25** University of Surrey, **26** Wellcome Centre for Human Genetics, Nuffield Department of Medicine, University of Oxford, **27** Big Data Institute, Nuffield Department of Medicine, University of Oxford, **28** Oxford University Hospitals NHS Foundation Trust, **29** Basingstoke Hospital, **30** Centre for Genomic Pathogen Surveillance, University of Oxford, **31** Hampshire Hospitals NHS Foundation Trust, **32** University of Southampton, **33** Public Health Wales NHS Trust, **34** Cardiff University, **35** Betsi Cadwaladr University Health Board, **36** Cardiff and Vale University Health Board, **37** Swansea University, **38**

University of Sheffield, **39** Sheffield Teaching Hospitals, **40** Great Ormond Street NHS Foundation Trust, **41** University College London, **42** Oswaldo Cruz Institute, Rio de Janeiro **43** North West London Pathology, **44** Imperial College Healthcare NHS Trust, **45** NIHR Health Protection Research Unit in HCAI and AMR, Imperial College London, **46** University of Liverpool, **47** Manchester University NHS Foundation Trust, **48** Liverpool Clinical Laboratories, **49** University of Exeter, **50** Royal Devon and Exeter NHS Foundation Trust, **51** Quadram Institute Bioscience, University of East Anglia, **52** Norfolk and Norwich University Hospital, **53** University of East Anglia, **54** Deep Seq, School of Life Sciences, Queens Medical Centre, University of Nottingham, **55** Virology, School of Life Sciences, Queens Medical Centre, University of Nottingham, **56** Clinical Microbiology Department, Queens Medical Centre, **57** PathLinks, Northern Lincolnshire & Goole NHS Foundation Trust, **58** Clinical Microbiology, University Hospitals of Leicester NHS Trust, **59** Viapath, **60** Hub for Biotechnology in the Built Environment, Northumbria University, **61** NU-OMICS Northumbria University, **62** Northumbria University, **63** South Tees Hospitals NHS Foundation Trust, **64** North Cumbria Integrated Care NHS Foundation Trust, **65** North Tees and Hartlepool NHS Foundation Trust, **66** Newcastle Hospitals NHS Foundation Trust, **67** County Durham and Darlington NHS Foundation Trust, **68** Centre for Enzyme Innovation, University of Portsmouth, **69** School of Biological Sciences, University of Portsmouth, **70** Portsmouth Hospitals NHS Trust, **71** University of Warwick, **72** University Hospitals Coventry and Warwickshire, **73** Warwick Medical School and Institute of Precision Diagnostics, Pathology, UHCW NHS Trust, **74** Genomics Innovation Unit, Guy's and St. Thomas' NHS Foundation Trust, **75** Centre for Clinical Infection & Diagnostics Research, St. Thomas' Hospital and Kings College London, **76** Department of Infectious Diseases, King's College London, **77** Guy's and St. Thomas' Hospitals NHS Foundation Trust, **78** Centre for Clinical Infection and Diagnostics Research, Department of Infectious Diseases, Guy's and St Thomas' NHS Foundation Trust, **79** Princess Alexandra Hospital Microbiology Dept. , **80** Cambridge University Hospitals NHS Foundation Trust, **81** East Kent Hospitals University NHS Foundation Trust, **82** University of Kent, **83** Gloucestershire Hospitals NHS Foundation Trust, **84** Department of Microbiology, Kettering General Hospital, **85** National Infection Service, PHE and Leeds Teaching Hospitals Trust, **86** Cambridge Stem Cell Institute, University of Cambridge, **87** Public Health Scotland, 88 Belfast Health & Social Care Trust, **89** Health Services Laboratories, **90** Barking, Havering and Redbridge University Hospitals NHS Trust, **91** Royal Free NHS Trust, **92** Maidstone and Tunbridge Wells NHS Trust, **93** University of Brighton, **94** Kings College London, **95** PHE Heartlands, **96** Imperial College London, **97** Department of Infection Biology, London School of Hygiene and Tropical Medicine.

#### Supplementary Files

**supplementary_file_f1_cannonical_orf_counts.csv**
All canonical ORF counts for all 1155 samples.

**supplementary_file_f2_novel_orf_counts.csv**
All non-canonical ORF counts for all 1155 samples.

**supplementary_file_f3_downsampling.csv**
All canonical ORF counts for all samples used in the downsampling experiment.

**supplementary_file_f4_repeats.csv**
All canonical ORF counts for all samples used in the replicates experiment.

**supplementary_file_f5_illumina_in_vitro.csv**
Periscope counts for Illumina metagenomic data.

**supplementary_file_f6_ont_in_vitro.csv**
Periscope counts for ONT ARTIC *in vitro* data.

**supplementary_file_f7_glasgow_ont.csv**
Canonical periscope counts for Glasgow ONT ARTIC data.

**supplementary_file_f8_sample_run_info.csv**
Run information for the samples in the study for PCA analysis.

**supplementary_file_f9_sample_ect.csv**
E cycle threshold values for samples in the study.

**supplementary_file_f10_novel_orf_counts_glasgow.csv**
Non-canonical sgRNA counts for Glasgow ONT ARTIC data.

**supplementary_file_f11_glasgow_bait_capure_novel.csv**
Non-canonical sgRNA counts for Glasgow bait capture Illumina data.

**supplementary_file_f12_illumina_in_vitro_novel.csv**
Non-canonical sgRNA counts for the *in vitro* dataset sequenced using an Illumina metagenomic approach.

**supplementary_file_f13_ont_in_vitro_novel.csv**
Non-canonical sgRNA counts for the *in vitro* dataset sequenced using an ONT ARTIC approach.

**supplementary_file_f14_glasgow_bait_capure.csv**

Canonical periscope counts for Glasgow ONT ARTIC data.

**supplementary_file_f15_read_lengths.csv**
Read lengths for a representative sample classed into either sgRNA or gRNA.

**supplementary_file_f16_base_counts.csv**
Counts of bases at each variant position.

**Supplementary_file_f17_coverage.csv**
Consensus coverage of samples used in the study.

**supplementary_file_f18_figure_generation.Rmd**
All R markdown code used in generation of figures contained within this manuscript.

**supplementary_file_f19_periscope-0.0.8a.tar.gz**
periscope source code.

Common column names contained within these files:

| Column Name | Contents |
|---|---|
| gRNA_count | Raw count of genomic reads |
| sgRNA_HQ_count, sgRNA_LQ_count, sgRNA_LLQ_count | Raw count of sub-genomic reads classified as high quality, low quality or low low quality |
| gRPHT | Genomic reads per 100,000 mapped reads |
| sgRPTg_HQ, sgRPTg_LQ, sgRPTg_LLQ | High, low and low low quantity sub-genomic RNA reads per 1000 genomic |
| sgRPTg_ALL | Sum of all sub-genomic reads normalised per 1000 genomic |
| sgRPHT_HQ, sgRPHT_LQ, sgRPHT_LLQ | High, low and low low quantity sub-genomic RNA reads per 100,000 mapped reads |
| sgRPHT_ALL | Sum of all sub-genomic reads normalised per 100,000 mapped |
| nsgRNA_HQ_count, nsgRNA_LQ_count, nsgRNA_LLQ_count | Raw count of non-canonical sub-genomic reads classified as high quality, low quality or low low quality |
| nsgRPTg_HQ, nsgRPTg_LQ, nsgRPTg_LLQ | High, low and low low quantity non-canonical sub-genomic RNA reads per 1000 genomic |
| nsgRPTg_ALL | Sum of all non-canonical sub-genomic |

| | |
|---|---|
| | reads normalised per 1000 genomic |
| nsgRPHT_HQ, nsgRPHT_LQ, nsgRPHT_LLQ | High, low and low low quantity non-canonical sub-genomic RNA reads per 100,000 mapped reads |
| nsgRPHT_ALL | Sum of all non-canonical sub-genomic reads normalised per 100,000 mapped |
| sgRPTL | Sub-genomic reads per 1000 reads of local coverage |

### Periscope README

Below we have provided instructions for the installation, execution and interpretation of periscope.

#### Requirements

periscope runs on MacOS, unix and unix subsystem for windows 10.

You will need:

- conda
- Your raw fastq files from the ARTIC protocol
- Periscope installation

In our hands periscope takes around 1-5 minutes per million reads on a single core on a Dell XPS core i9 with 32Gb ram and 1Tb SSD.

#### Installation

```
git clone https://github.com/sheffield-bioinformatics-core/periscope.git
&& cd periscope
conda env create -f environment.yml
conda activate periscope
python setup.py install
```

#### Execution

```
conda activate periscope

periscope
    --fastq-dir <PATH_TO_DEMUXED_FASTQ> (ont only)
    OR
    --fastq <FULL_PATH_OF_FASTQ_FILE(s)> (space separated list of fastq files,
```

```
you MUST use this for Illumina data)
    --output-prefix <PREFIX>
    --sample <SAMPLE_NAME>
    --artic-primers <ASSAY_VERSION; V1,V2,V3 or 2kb>
    --resources <PATH_TO_PERISCOPE_RESOURCES_FOLDER>
    --technology <SEQUENCING TECH; ont or illumina>
    --threads <THREADS_FOR_MAPPING>
```

#### Output Files

| **Filename** | **Description** |
| --- | --- |
| `<OUTPUT_PREFIX>.fastq` | A merge of all files in the fastq directory specified as input. |
| `<OUTPUT_PREFIX>_periscope_counts.csv` | The counts of genomic, sgRNA and normalisation values for *known* ORFs |
| `<OUTPUT_PREFIX>_periscope_amplicons.csv` | The amplicon by amplicon counts, this file is useful to see where the counts come from. Multiple amplicons may be represented more than once where they may have contributed to more than one ORF. (ONT only) |
| `<OUTPUT_PREFIX>_periscope_novel_counts.csv` | The counts of genomic RNA, sgRNA and normalisation values for *non-canonical* ORFs |
| `<OUTPUT_PREFIX>.bam &`<br>`<OUTPUT_PREFIX>.bam.bai` | Minmap2 or bwa mem mapped reads and index with no adjustments made. |
| `<OUTPUT_PREFIX>_periscope.bam &`<br>`<OUTPUT_PREFIX>_periscope.bam.bai &`<br>`<OUTPUT_PREFIX>_sorted_periscope.bam &`<br>`<OUTPUT_PREFIX>_sorted_periscope.bam.bai` | This is the original input bam file and index created by periscope with the reads specified in the fastq-dir. This file, however, has tags which represent the results of periscope:<br>● XS is the alignment score<br>● XA is the amplicon number<br>● XC is the assigned class (gDNA or sgDNA)<br>These are useful for manual review in IGV or similar genome viewer. You can sort or colour reads by these tags to aid in manual review and figure creation. |

#### Examining Base Frequencies of Called Variants in periscope

This script will take the pass vcf from the ARTIC Network pipeline and examine the periscope bam file for the bases present at that position. It will split the counts by read class and output a plot showing contribution at each base at each site in the VCF.

```
conda activate periscope

gunzip <ARTIC_NETWORK_VCF>.pass.vcf.gz

<PATH_TO_PERISCOPE>/periscope/periscope/scripts/variant_expression.py \
    --periscope-bam <PATH_TO_PERISCOPE_OUTPUT_BAM> \
    --vcf <ARTIC_NETWORK_VCF>.pass.vcf \
    --sample <SAMPLE_NAME> \
    --output-prefix <OUPUT_PREFIX>
```

| **Filename** | **Description** |
|---|---|
| `<OUTPUT_PREFIX>_base_counts.csv` | Counts of each base at each position |
| `<OUTPUT_PREFIX>_base_counts.png` | Plot of each position and base composition |

### Supplementary Methods

#### Glasgow Oxford Nanopore Amplicon Sequencing

Sequencing libraries were prepared according to the ARTIC nCoV-2019 sequencing protocol version 2, described in detail at https://artic.network/ncov-2019. For amplicon generation cDNA was synthesised from extracted RNA using SuperScript IV (Thermo Scientific, Part Number 18090200) and random hexamer primers (Part number N8080127). PCR amplicons (approximately 400 bp) were generated from this cDNA using Q5® Hot Start High-Fidelity 2X Master Mix (New England Biolabs, Part Number M0494L), nCoV-2019 PrimalSeq sequencing primers (Version 3) and 25-35 cycles of the ARTIC nCoV-2019 recommended thermocycling conditions. The PCR amplicons were purified using Agencourt AMPure XP for PCR Purification (Beckman Coulter, Part Number A63881), following the manufacturer's guidelines and quantified using the Qubit dsDNA HS Kit (Thermo Scientific, Part Number Q32854). Generated amplicons were then used to prepare Oxford Nanopore sequencing libraries.

End repair of amplicons (200 fmol) was carried out using the NEBNext Ultra II End repair /dA-tailing Module (New England Biolabs, Part Number E7546L), then 20 fmol of end prepped amplicon was barcoded using the Native Barcoding Expansion 1-12 (PCR free) kit (Oxford Nanopore Technologies, Part Number EXP-NBD104) and NEBNext® Ultra™ II Ligation Module (New England Biolabs, Part Number E7595L). Barcoded amplicons were pooled together and purified using Agencourt AMPure XP for PCR Purification (Beckman Coulter, Part Number A63881), following the manufacturer's guidelines and quantified using the Qubit dsDNA HS Kit (Thermo Scientific, Part Number Q32854). Adapter mix II (Oxford Nanopore Technologies, Part Number EXP-NBD104) was ligated to the barcoded amplicons using the Ligation Sequencing kit (Oxford Nanopore Technologies, SQK-LSK109) and NEBNext Quick Ligation Module (New England Biolabs, Part Number E6056L). The final library was purified using Agencourt AMPure XP for PCR Purification (Beckman Coulter, Part Number A63881), following the manufacturer's guidelines and quantified using the Qubit dsDNA HS Kit (Thermo Scientific, Part Number Q32854). Approximately 50 fmol of the sequencing library pool was loaded onto a flow cell (R9.4.1) (Oxford Nanopore Technologies, Part Number FLO-MIN106D) where sequencing was conducted in MinKNOW version 19.12.6 and raw FAST5 files were basecalled using Guppy version 3.2.10 in high accuracy mode using a minimum quality score of 7.

#### Glasgow Bait Capture & Subsequent Illumina Sequencing

A mild DNAse treatment was done using DNAse1 (Thermo Fisher, Part Number AM2222) and samples were purified using RNA Clean AMPure XP Beads (Beckman Coulter, A63987). cDNA was synthesised using SuperScript III (Thermo Scientific, Part Number 18080044) and NEBNext Ultra II Non-Directional RNA Second Strand Synthesis Module (New England Biolabs, Part Number E6111L). Reagents from the Illumina Nextera Flex for Enrichment (Cat. No. 20025523 and 20025524) were used for library preparation and targeted enrichment. Briefly, The cDNA was tagmented and followed through to PCR with the indexed primers. 12 cycles of PCR were used as per protocol recommendations using the IDT for Illumina Nextera DNA Unique Dual Indexes Set A (Illumina Part Number 20027213). The amplified libraries were quantified by Qubit dsDNA HS Kit and run on the Agilent 4200 Tapestation System (Agilent, Part Number G2991AA) using the High Sensitivity D5000 Screentape (Agilent, Part Number 5067-5592) and High Sensitivity D5000 Reagents (Agilent, Part Number 5067-5593), to guide sample pooling. Baits from Illumina (Respiratory Virus Oligos Panel V2, Part Number 20044311) were hybridized with the sample pool overnight at 62oC, as recommended by Illumina. After capture and wash, 12 cycles of PCR were used to amplify captured DNA. The amplified pools were quantified by Qubit dsDNA HS Kit and run on the Agilent 4200 Tapestation System using the High Sensitivity D5000 Assay to determine the size of the pool in base pairs (bp). The sequencing of the enriched pools was carried out on Illumina's MiSeq System (Illumina, Part Number SY-410-1003) using a MiSeq Reagent Kit v3 600 cycle kit (Illumina, Part Number MS-102-3003) with more than 70% bases presenting a Q score superior to 30.

### Supplementary Tables

| ORF | Samples with >=1 (HQ+LQ Reads) | Percent of Total |
|---|---|---|
| E | 1046 | 90.6 |
| M | 1109 | 96.0 |
| N | 1124 | 97.3 |
| ORF10 | 11 | 1.0 |
| ORF3a | 673 | 58.3 |
| ORF6 | 1053 | 91.2 |
| ORF7a | 906 | 78.4 |
| ORF8 | 274 | 23.7 |
| S | 1071 | 92.7 |

**Supplementary Table S1 - sgRNAs detected for each canonical ORF**

Raw counts of the number of sgRNAs found across all 1,155 samples of the cohort with 1 or more HQ or LQ read.

| Sample | Genomic RNA | HQ Sub-Genomic | LQ Sub-Genomic |
|---|---|---|---|
| SHEF-C00C0 | 3793 | 0 | 2 |
| SHEF-C045B | 911 | 0 | 1 |
| SHEF-C046A | 1340 | 0 | 1 |
| SHEF-C0840 | 1829 | 1 | 0 |
| SHEF-C09F2 | 2462 | 0 | 1 |
| SHEF-C58A5 | 4611 | 1 | 0 |
| SHEF-C722D | 1536 | 0 | 1 |
| SHEF-C8408 | 4237 | 0 | 1 |
| SHEF-CF595 | 5173 | 1 | 0 |
| SHEF-D179A | 2768 | 1 | 0 |
| SHEF-D227A | 3802 | 0 | 1 |

**Supplementary Table S2 - Samples with predicted sgRNA for ORF10**

Samples with any HQ or LQ evidence of ORF10 sgRNA (amplicons 97 and 98). Twelve reads in total across the whole cohort putatively support a sgRNA for ORF10 .

| Read Start Position | Contributing Amplicon | Total Genomic Read Count | Total HQ sgRNA Read Count | Total LQ sgRNA Read Count | Total sgRNA Read Count |
|---|---|---|---|---|---|
| 25744 | 84,85 | 6019 | 120 | 33 | 153 |
| 25745 | 85 | 1389 | 4 | 5 | 9 |
| 25746 | 85 | 1389 | 1 | 2 | 3 |
| 25748 | 85 | 1389 | 2 | 1 | 3 |
| 25749 | 85 | 1389 | 1 | 0 | 1 |
| 25754 | 85 | 1389 | 2 | 0 | 2 |
| 25755 | 85 | 1389 | 1 | 0 | 1 |
| 25732 | 85 | 1389 | 0 | 1 | 1 |
| 25735 | 85 | 1389 | 0 | 1 | 1 |
| 25742 | 85 | 1389 | 0 | 1 | 1 |
| 25753 | 85 | 1389 | 0 | 1 | 1 |
| 25766 | 85 | 1389 | 0 | 1 | 1 |

**Supplementary Table S3 - Non-canonical sgRNA at position 25,744 in sample SHEF-C0118 has strong support**

| ORF | Contributing Amplicon | Total Genomic Read Count | Total HQ sgRNA Read Count | Total LQ sgRNA Read Count | Total sgRNA Read Count |
|---|---|---|---|---|---|
| S | 71,72 | 4939 | 44 | 31 | 75 |
| ORF3a | 83,84 | 8147 | 3 | 2 | 5 |
| E | 86,87,88 | 11183 | 12 | 7 | 19 |
| M | 87,88 | 6590 | 192 | 90 | 282 |
| ORF6 | 89 | 564 | 60 | 24 | 84 |
| ORF7a | 90,91 | 5580 | 0 | 16 | 16 |
| N | 93,94 | 8928 | 500 | 251 | 751 |
| **Non-Canonical sgRNA** | **84,85** | **6019** | **131** | **46** | **177** |

**Supplementary Table S4 - Canonical sgRNA in SHEF-C0118**

Showing only those ORF sgRNAs with supporting Reads.

| Read Start Position | Contributing Amplicon | Total Genomic Read Count | Total HQ sgRNA Read Count | Total LQ sgRNA Read Count | Total sgRNA Read Count |
|---|---|---|---|---|---|
| 10639 | 35,36 | 4698 | 103 | 15 | 118 |
| 10640 | 35 | 3583 | 0 | 1 | 1 |
| 10641 | 35 | 3583 | 2 | 3 | 5 |
| 10642 | 35,36 | 4698 | 1 | 4 | 5 |
| 10643 | 36 | 1115 | 1 | 0 | 1 |
| 10644 | 35 | 3583 | 1 | 0 | 1 |
| 10645 | 35 | 3583 | 2 | 2 | 4 |
| 10647 | 35 | 3583 | 0 | 1 | 1 |

**Supplementary Table S5 - Non-canonical sgRNA at 10,639 in SHEF-CE04A has strong support**

Counts of reads supporting the sgRNA at 10,639 in SHEF-CE04A.

| Sample | Read Start Position | Total Genomic Read Count | Total HQ sgRNA Read Count | Total LQ sgRNA Read Count |
|---|---|---|---|---|
| CVR2185 | 5785 | 12689 | 3 | 0 |
| CVR2187 | 5785 | 20464 | 1 | 2 |
| CVR2191 | 5785 | 5479 | 1 | 0 |
| CVR2231 | 5785 | 7561 | 2 | 0 |
| CVR2234 | 5785 | 8369 | 1 | 0 |
| CVR2239 | 5785 | 12261 | 2 | 0 |
| CVR2243 | 5785 | 21022 | 1 | 0 |
| CVR2251 | 5785 | 5907 | 1 | 1 |
| CVR2265 | 5785 | 13147 | 0 | 1 |
| CVR2306 | 5785 | 8442 | 4 | 1 |
| CVR2319 | 5785 | 4785 | 2 | 0 |
| CVR2322 | 5785 | 5926 | 1 | 0 |
| CVR2484 | 5785 | 3096 | 7 | 0 |
| CVR3941 | 5785 | 10157 | 2 | 0 |
| CVR3943 | 5785 | 5521 | 0 | 1 |

**Supplementary Table S6 - Non-canonical sgRNA at 5,785 in Glasgow Nanopore samples**

Fifteen samples in the Glasgow ONT ARTIC dataset have evidence for a non-canonical sgRNA at 5,785

| Sample | Read Start Position | Total Genomic Read Count | Total HQ sgRNA Read Count | Total LQ sgRNA Read Count |
|---|---|---|---|---|
| CVR2131 | 10639 | 9802 | 3 | 1 |
| CVR2133 | 10639 | 312 | 1 | 0 |
| CVR2166 | 10639 | 11494 | 1 | 1 |
| CVR2185 | 10639 | 4653 | 1 | 2 |
| CVR2187 | 10639 | 9927 | 3 | 0 |
| CVR2190 | 10639 | 1607 | 1 | 0 |
| CVR2191 | 10639 | 2107 | 1 | 0 |
| CVR2247 | 10639 | 11675 | 1 | 4 |
| CVR2265 | 10639 | 6284 | 7 | 1 |
| CVR2306 | 10639 | 3537 | 4 | 1 |
| CVR2319 | 10639 | 1772 | 5 | 2 |
| CVR2322 | 10639 | 2114 | 1 | 2 |
| CVR3940 | 10639 | 5916 | 1 | 0 |

**Supplementary Table S7 - Non-canonical sgRNA at 10,639 in Glasgow Nanopore samples**

Thirteen samples from the ONT ARTIC Glasgow dataset have evidence for a non-canonical sgRNA at 10,639.

| ORF | Contributing Amplicon | Total Genomic Read Count | Total HQ sgRNA Read Count | Total LQ sgRNA Read Count | Total sgRNA Read Count |
|---|---|---|---|---|---|
| S | 71,72 | 4945 | 64 | 16 | 80 |
| ORF3a | 84,85 | 6990 | 8 | 1 | 9 |
| M | 87 | 4456 | 176 | 43 | 219 |
| ORF6 | 89 | 1527 | 0 | 0 | 0 |
| N | 93 | 8083 | 116 | 38 | 154 |
| **Non-Canonical sgRNA** | **35,36** | **4698** | **110** | **26** | **136** |

**Supplementary Table S8 - Canonical sub-genomic RNA in SHEF-CE04A**

Showing only those ORF sgRNAs with supporting Reads

| Virus | SNP | Amino Acid Change | |
|---|---|---|---|
| | | Gene | Mutation |
| PHE2 | A2618G | nsp2 | I605V |
| | C8782T | nsp14 | I150T |
| | T18488C | S | T95I |
| | C21846T | E | L37H |
| | T23605G | ORF 8 | L84S |
| | T26354A | N | P365S |
| | T28144C | ORF 10 | I13M |
| | C29366T | | |
| | A29596G | | |
| GLA1 | C3037T | nsp12 | P323L |
| | C14408T | S | D614G |
| | A23403G | E | V5A |
| | A24388T | | |
| | T26258C | | |
| GLA2 | C3037T | nsp12 | P323L |
| | C14408T | nsp15 | V35A |
| | T19724C | S | N439K |
| | C22879A | S | D614G |
| | A23403G | ORF 10 | V6F |
| | G29573T | | |

**Supplementary Table S9 - Changes with respect to MN908947.3 in viral isolates used in *in vitro* SARS-CoV-2 infection model**

### Supplementary Figures

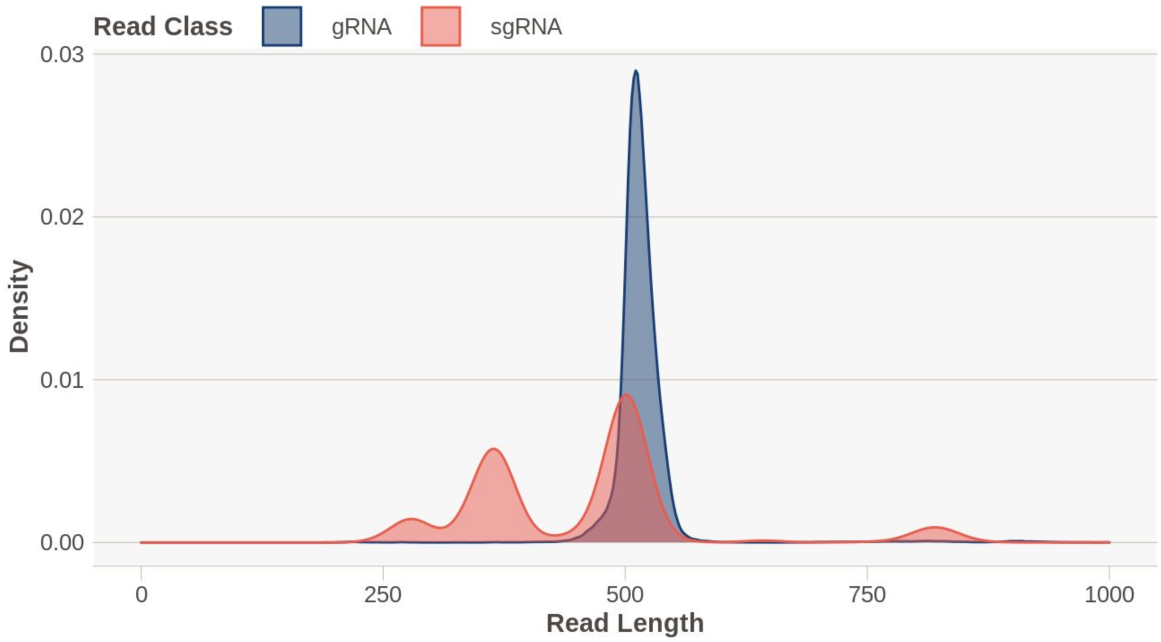


**Supplementary Figure S1 - Length of reads in SHEF ONT ARTIC data broken down by periscope assigned class in a representative sample (SHEF-BFDBE)**

At each of the amplicons responsible for the production of sgRNA supporting reads we examined the size of the two classes of reads. Genomic RNA is between 400 and 600bp, and in most cases sgRNA is shorter at around 200-300bp.

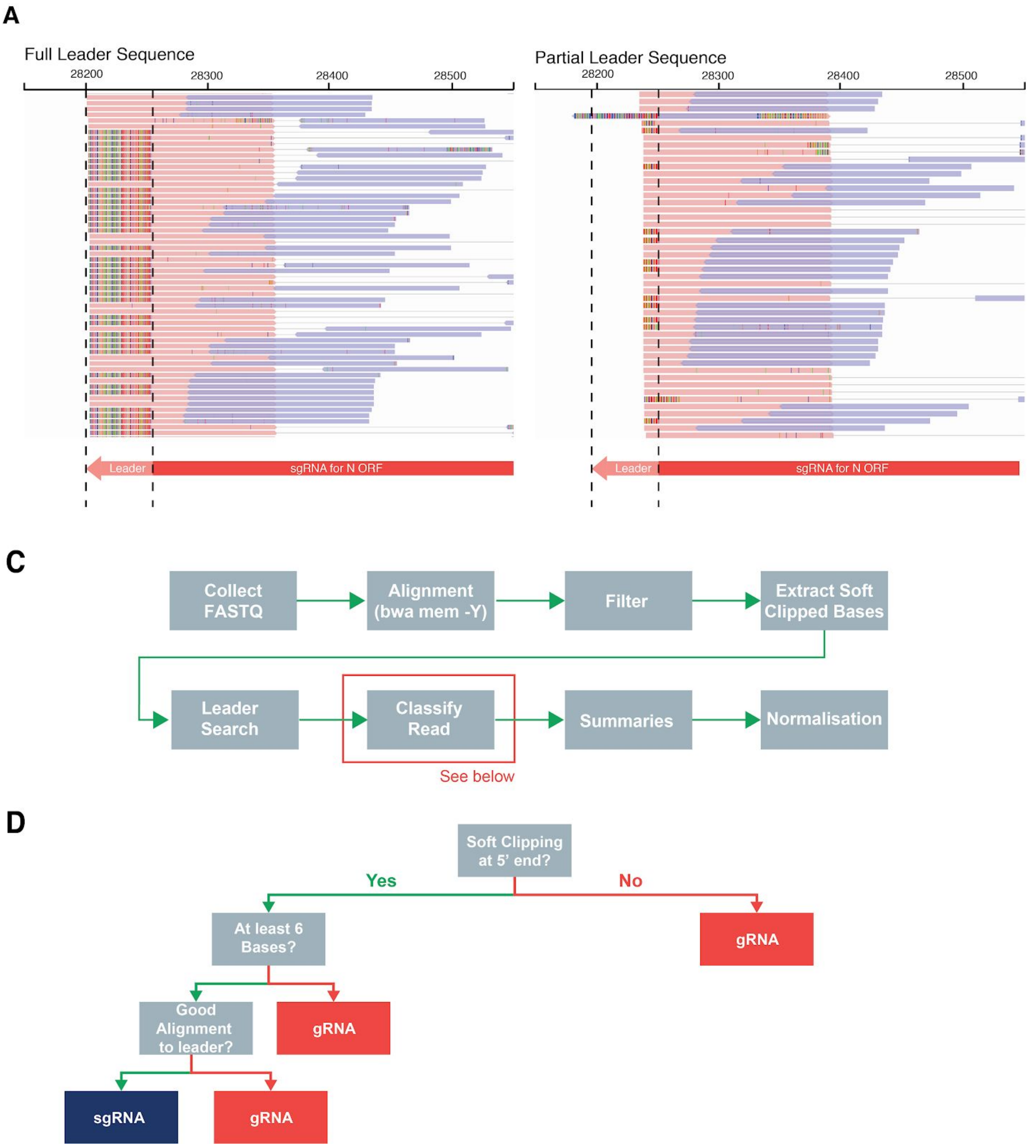


**Supplementary Figure S2 - SARS-CoV-2 Illumina Data**

**A**. Bait based capture and metagenomic sequencing of SARS-CoV-2 genomes by illumina methods results in variable lengths of leader sequence included in the read. **B.** Overall workflow for analysis of Illumina data is very similar to that of ARTIC Nanopore data, but adjusted in light the phenomenon described in (**A**). **C.** Classification of reads from illumina data involves extracting soft clipped bases from the 5' end of reads and performing a local alignment of these to the leader sequence. Only those reads that have >= 6 bases soft-clipped are considered for leader matching to reduce false positives.

A

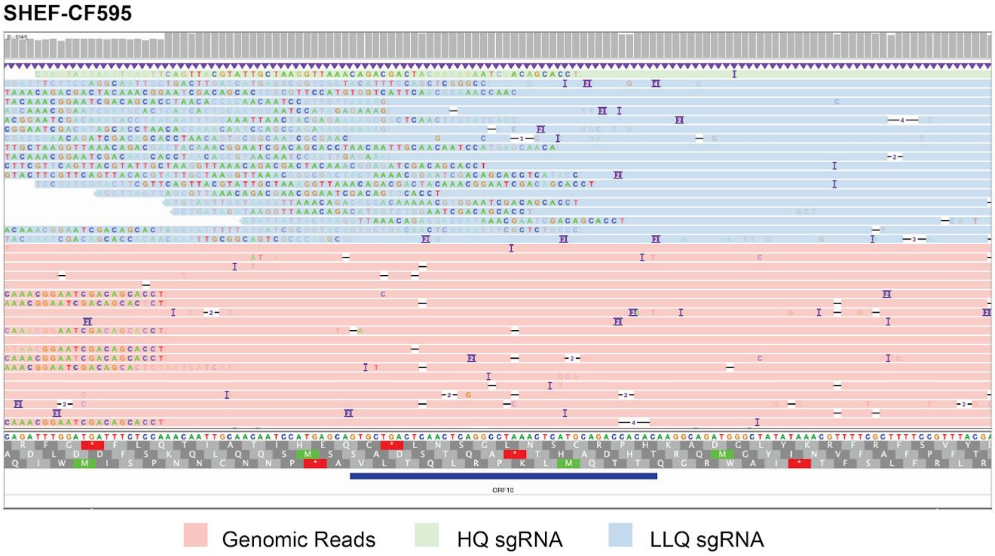


B

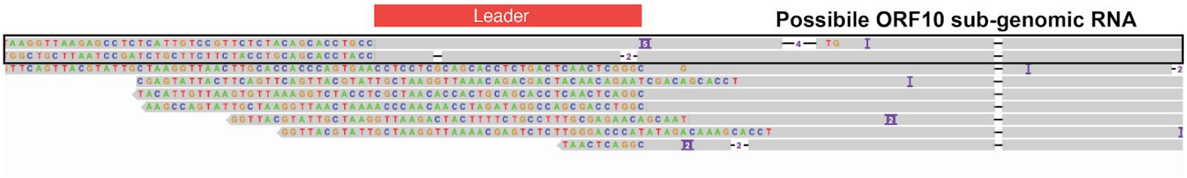


**Supplementary Figure S3 - Manual review of periscope bam files for ORF10**

**A.** SHEF-CF595 has 1 read classified as HQ sgRNA at the predicted ORF10 leader junction (light green). It is clear from this IGV screenshot that this read does not contain a valid leader sequence. **B.** All HQ and LQ sgRNA reads, 12 in total, aligned with minimap2 to a reference consisting of ORF10 and leader sequence, 3 reads failed to map. Two reads could be bona fide ORF10 sgRNAs (Highlighted in the black box with a good match to the leader) from samples SHEF-C0840 and SHEF-C58A5.

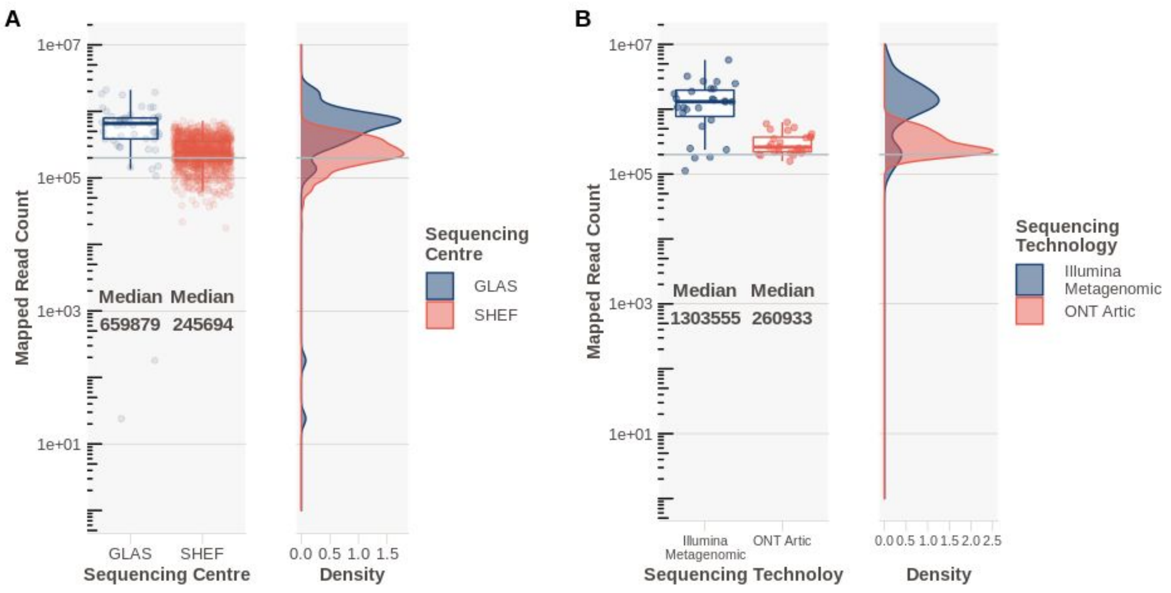


**Supplementary Figure S4 - Mapped read counts**

Reads were mapped with minimap2 (ONT) or bwa mem (Illumina) to MN908947.3, sorted with samtools and mapped reads counted with pysam. Grey horizontal line represents 200,000 mapped reads. Right plot in each panel is a density distribution of the points in the left panel. **A.** Data from SARS-CoV-2 clinical samples, sequenced with ONT Artic at Sheffield and Glasgow. **B.** Mapped read counts from SARS-CoV-2 *in vitro* infection models for Illumina metagenomic and ONT Artic data.

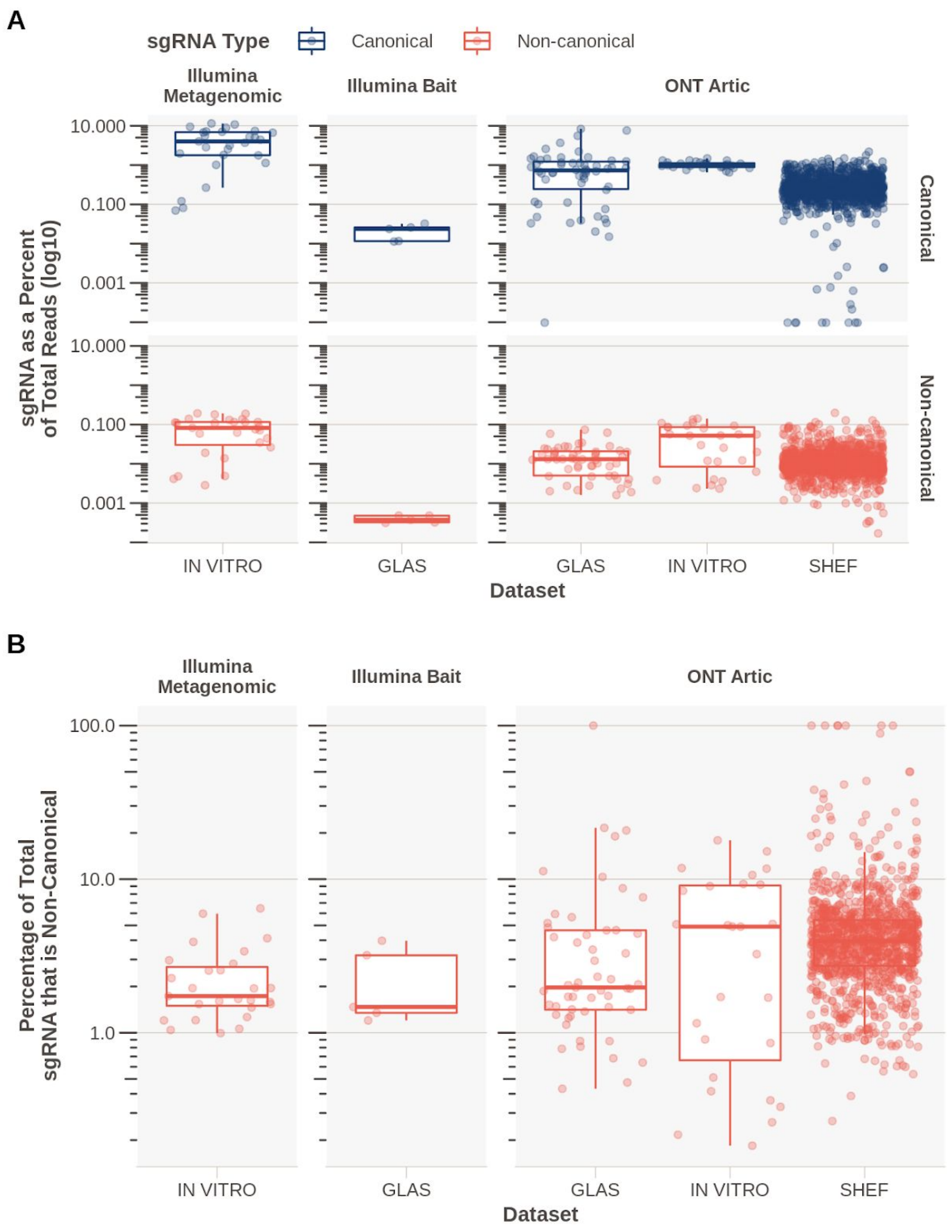


**Supplementary Figure S5 - Sub-Genomic RNA Proportions**

**A.** Total sgRNA reads split by canonical and non-canonical as a proportion of the total mapped reads in all featured datasets. **B.** Proportion of total sgRNA that is non-canonical.

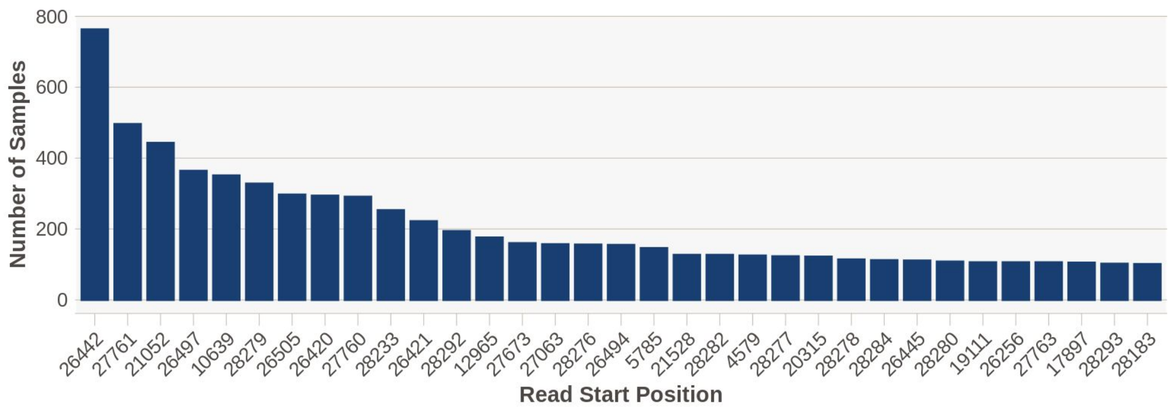


**Supplementary Figure S6 - Number of samples with at the most frequently represented non-canonical sub-genomic RNAs**

The number of samples each non-canonical sgRNA was found in. This is an exact position match, and includes sgRNAs that could be just outside the +/-20 of the TRS-B site. Sites with support in > 100 samples shown.

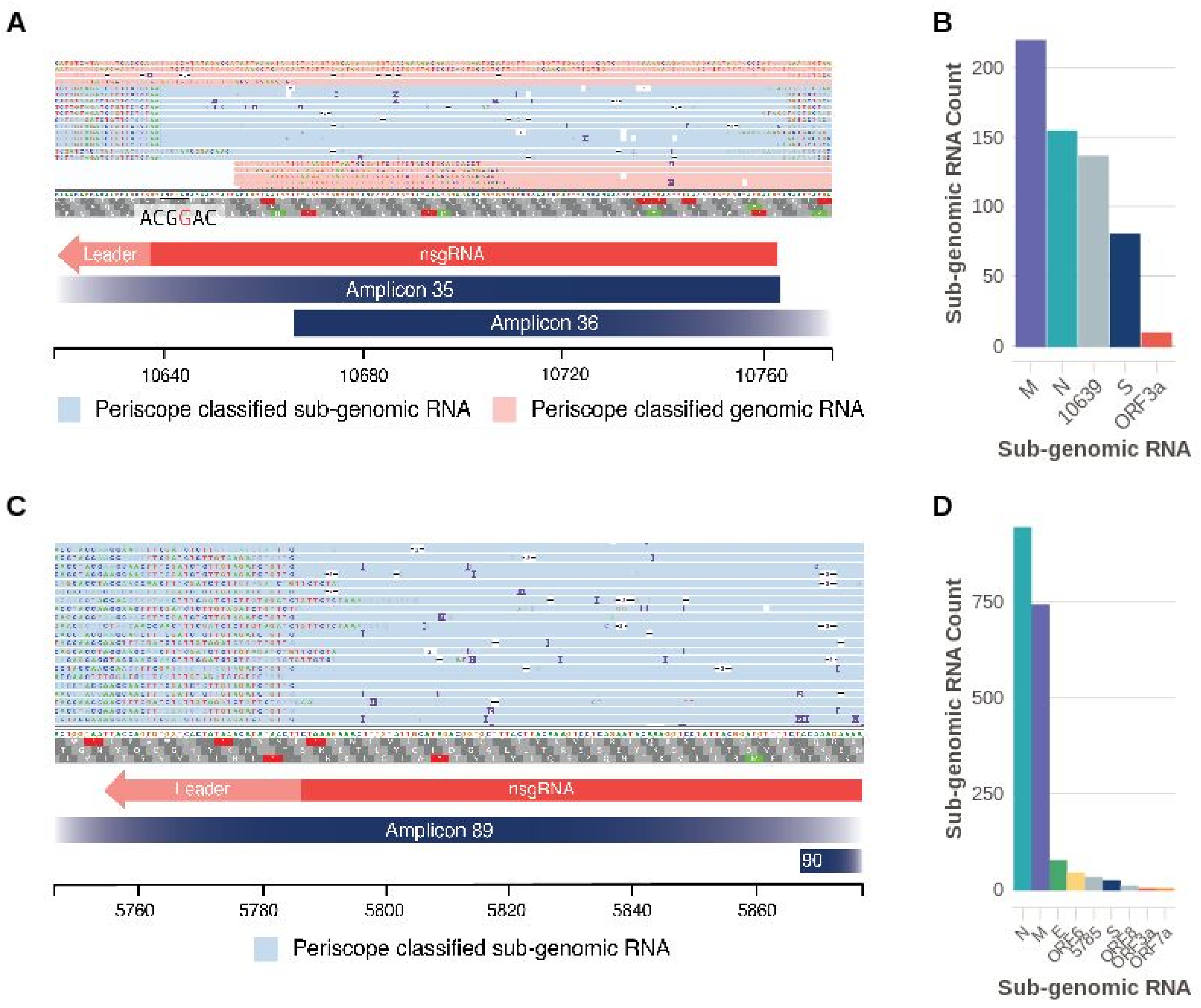


**Supplementary Figure S7 - Highly expressed non-canonical sgRNAs at 10,369 and 5,785**

**A.** Non-canonical sgRNA with strong support in SHEF-CE04A at 10,369 is also supported in additional 377 samples. In close proximity to the leader is a sequence which could be considered a “weak” TRS; ACGAAC -> ACG**G**AC. **B.** Raw sgRNA levels (HQ&LQ) in SHEF-CE04A show high relative amounts of this non-canonical sgRNA at 10639. (ORFs with sgRNA evidence shown) **C.** Non-canonical sgRNA at 5,785 (SHEF-BFD90 shown for illustration) found in 226 samples. **D**. Count of HQ&LQ raw sgRNAs in SHEF-D02E5.

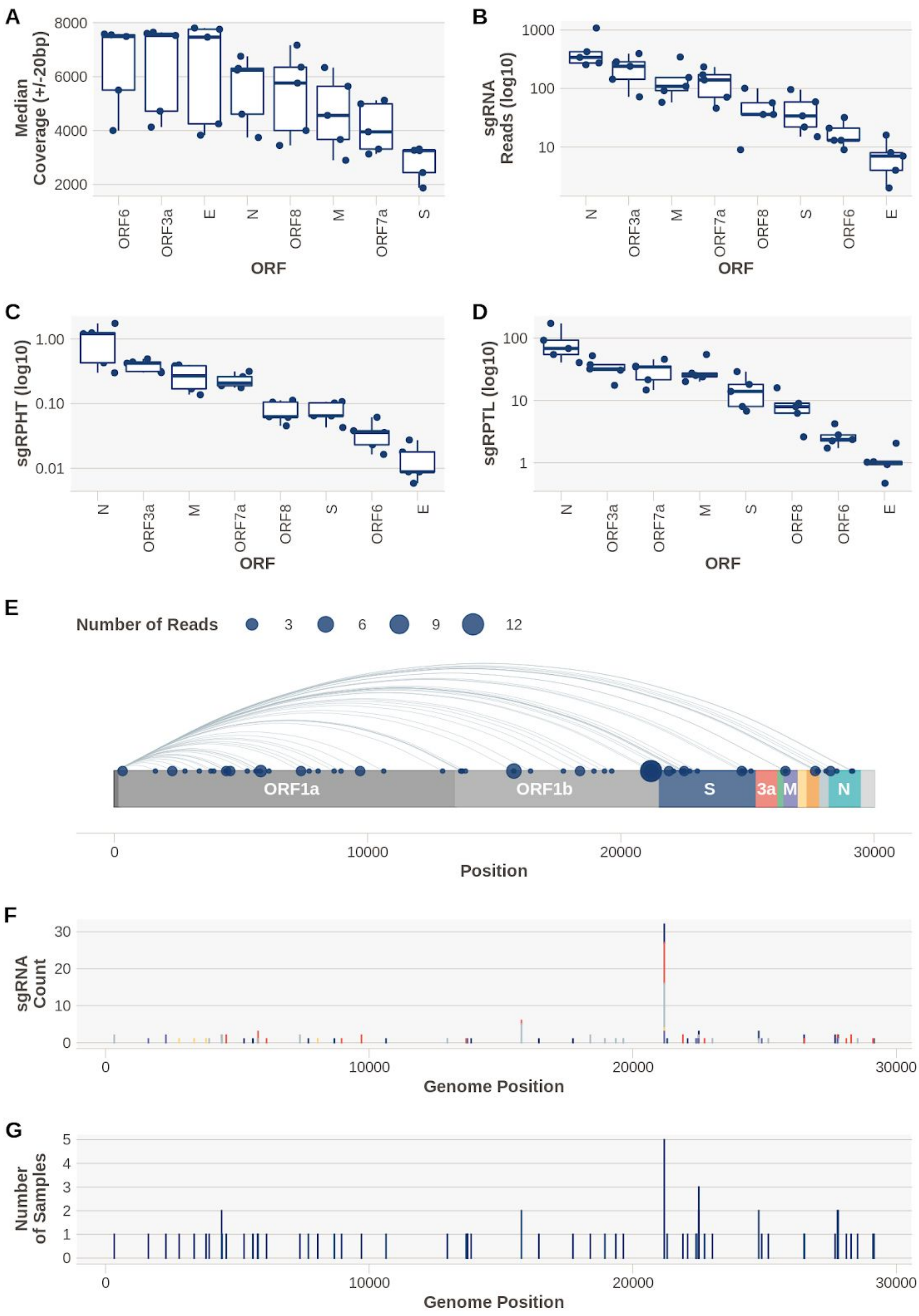

A
Median Coverage (+/-20bp)
8000
6000
4000
2000
ORF6
ORF3a
E
N
ORF8
M
ORF7a
S
ORF
B
sgRNA Reads (log10)
1000
100
10
N
ORF3a
M
ORF7a
ORF8
S
ORF6
E
ORF
C
sgRPHT (log10)
1.00
0.10
0.01
N
ORF3a
M
ORF7a
ORF8
S
ORF6
E
ORF
D
sgRPTL (log10)
100
10
1
N
ORF3a
ORF7a
M
S
ORF8
ORF6
E
ORF
E
Number of Reads
3
6
9
12
ORF1a
ORF1b
S
3a
M
N
0
10000
20000
30000
Position
F
sgRNA Count
30
20
10
0
0
10000
20000
30000
Genome Position
G
Number of Samples
5
4
3
2
1
0
0
10000
20000
30000
Genome Position

**Supplementary Figure S8 - sgRNA Levels in bait capture Illumina samples (n=5)**

**A.** Total coverage around canonical ORF TRS-B sites. **B.** Number of raw canonical sgRNA reads, no reads were found supporting ORF10. **C**. sgRNA reads normalised per 100,000 mapped reads showing that N is the most highly expressed sgRNA in this dataset. **D.** Normalising sgcounts to the local coverage (per 1000 reads +/- 20bp around TRS-B). **E.** Non-canonical sgRNA detected in this dataset. Size of point represents the total number of reads across all samples (n=5). **F.** Histogram showing the data in E, coloured by sample. **G.** Histogram of the number of samples each non-canonical sgRNA was detected in.

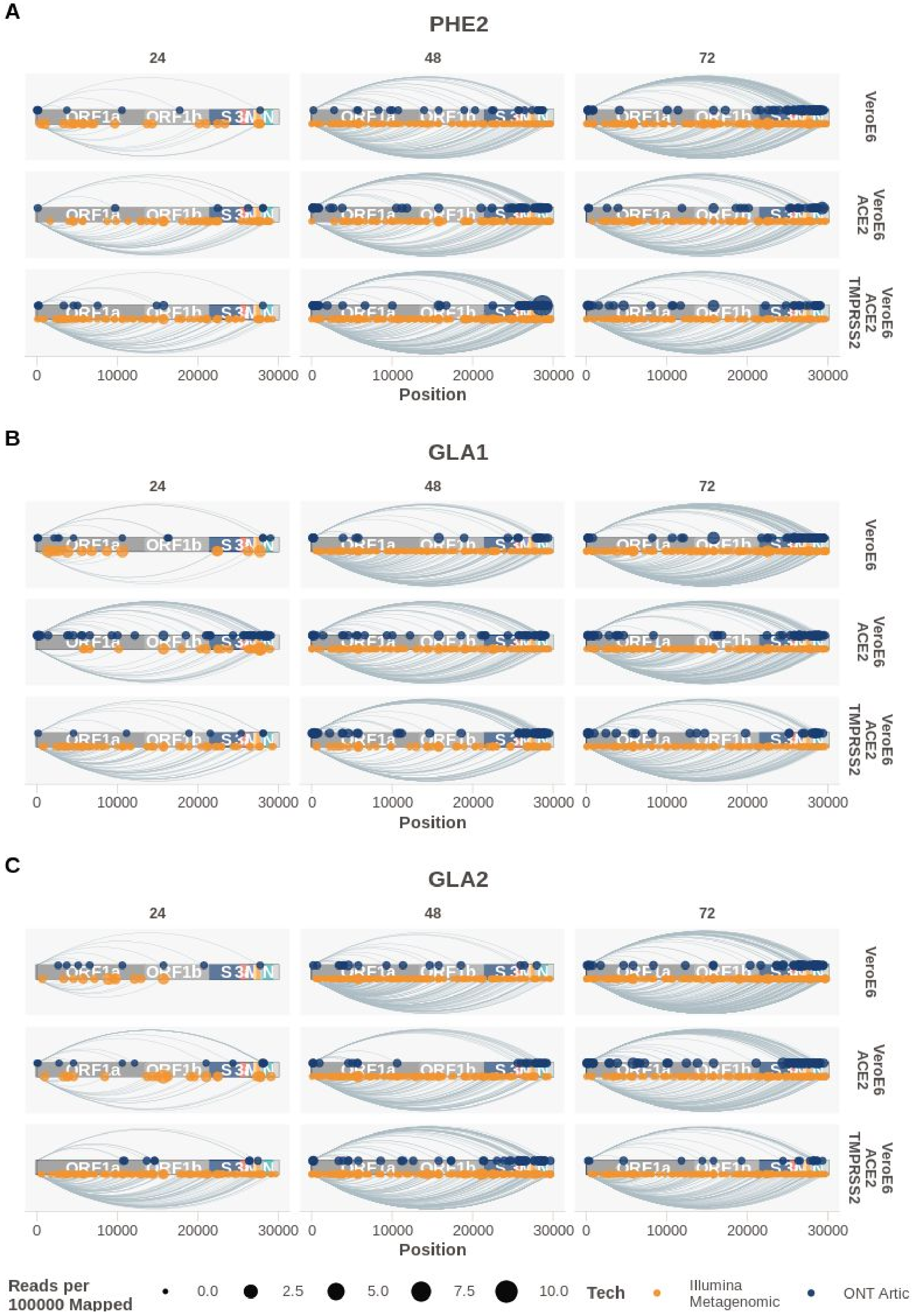
A
PHE2
B
GLA1
C
GLA2
24
48
72
VeroE6
VeroE6 ACE2
VeroE6 ACE2 TMPRSS2
ORF1a
ORF1b
S
3
M
N
0
10000
20000
30000
Position
Reads per 100000 Mapped
0.0
2.5
5.0
7.5
10.0
Tech
Illumina Metagenomic
ONT Artic

**Supplementary Figure S9 - Non-canonical sub-genomic RNA detected in an vitro infection model**

Results for WT VeroE6 and ACE2 & TMPRSS2 VeroE6 cells shown for viral isolates PHE2 (**A**), GLA1 (**B**), and GLA2 (**C**). Size of the point represents the number of reads per 100,000 mapped reads.

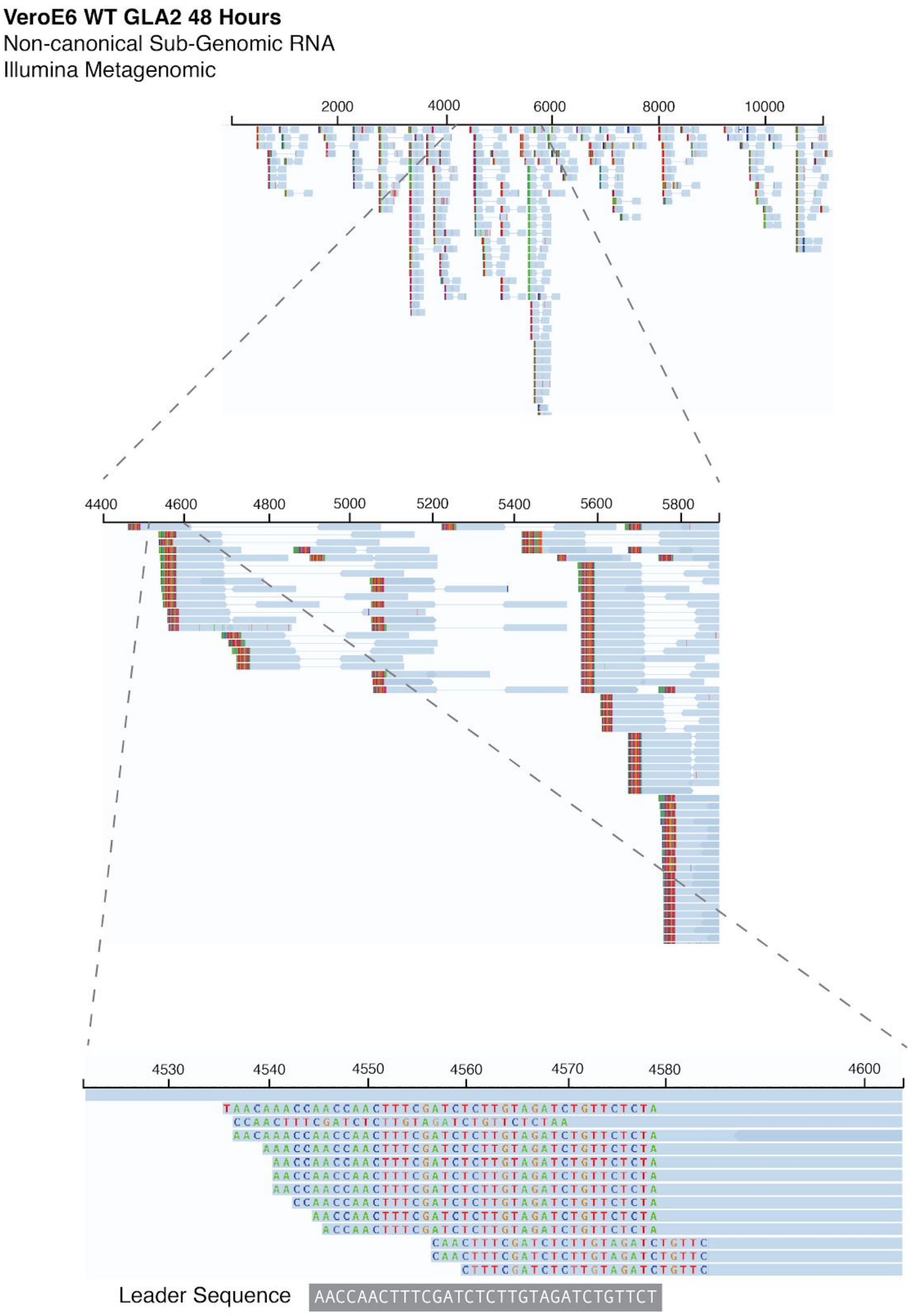


**Supplementary Figure S10 - Read level detail of Non-canonical sub-genomic RNA in Illumina metagenomic sequencing of an *in vitro* infection model**

IGV plots showing reads classified as non-canonical sgRNA by periscope in VeroE6 cells, after 48 hours of infection with SARS-CoV-2 (GLA2). Non-canoncial sgRNA can be seen throughout this region. These reads all contain leader sequence at their 5' end, as illustrated by the zoomed in panels.

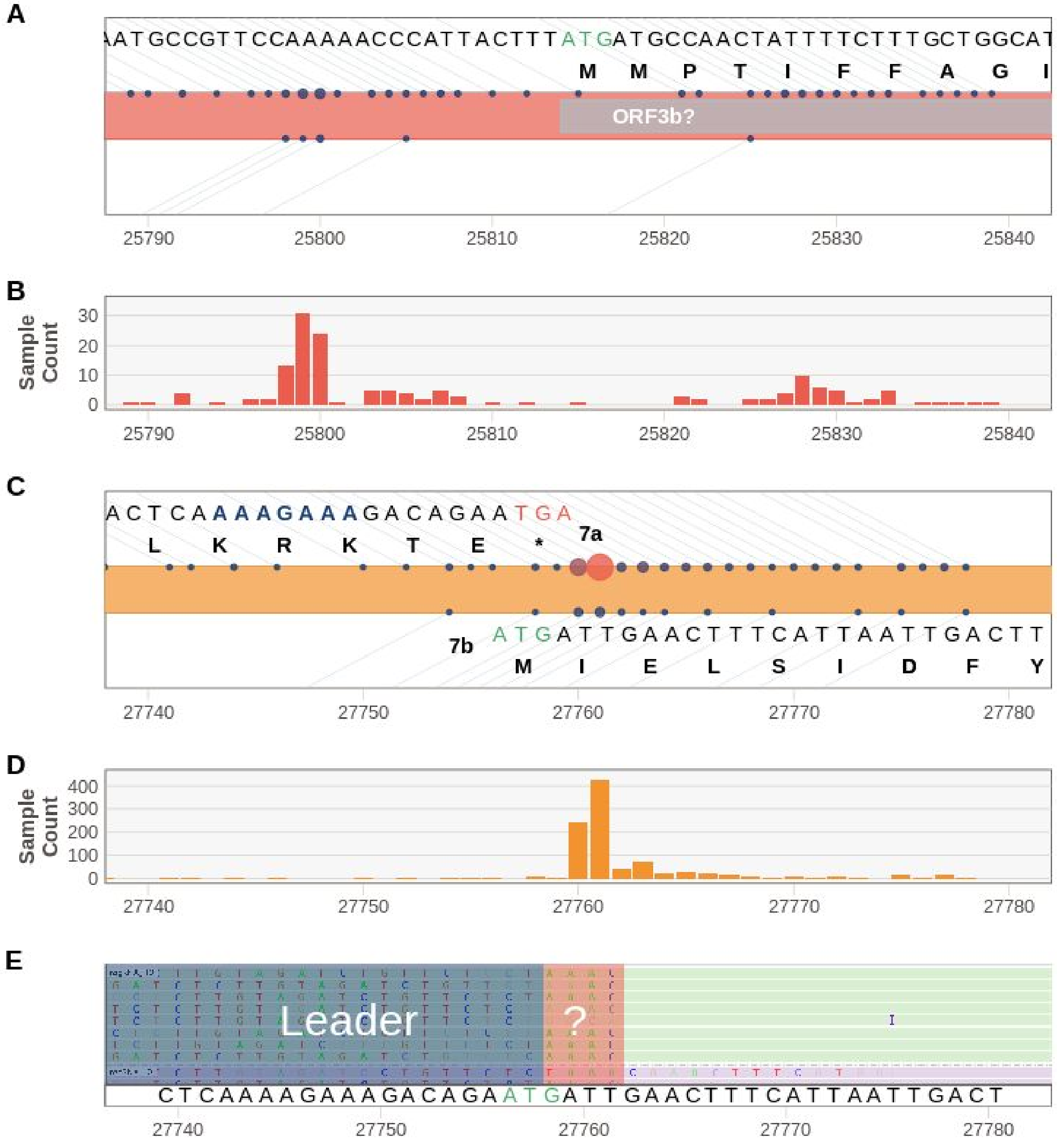


**Supplementary Figure S11 - Non-canonical sub-genomic RNAs (High quality) that could represent ORF3b and ORF7b**

**A.** All HQ non-canonical sgRNA between 25,790 and 25,840, Sheffield top, Glasgow bottom. It has been suggested a short 22 amino acid ORF 3b protein is a potent modulator of the interferon response (Konno et al. 2020) represented here in grey. **B.** Histogram of the number of samples from Sheffield with evidence of a non-canonical sgRNA at that position

(HQ). **C.** Non-canonical sgRNA between 27,740 and 27,780, Sheffield top, Glasgow bottom. Protein sequence shown for the C terminus of 7a and N terminus of 7b shown at the top and bottom respectively. Blue text indicates predicted TRS-B site for this ORF(Yang et al. 2020). **D.** Histogram of the number of samples from Sheffield with evidence of sgRNA at that position (HQ). **E.** Raw reads supporting the sgRNA at 27,761 showing the leader sequence, a mismatch of `AAAC`, followed by the genomic sequence for ORF7b. This shows that these sgRNAs do not include the predicted `ATG` for ORF7b.

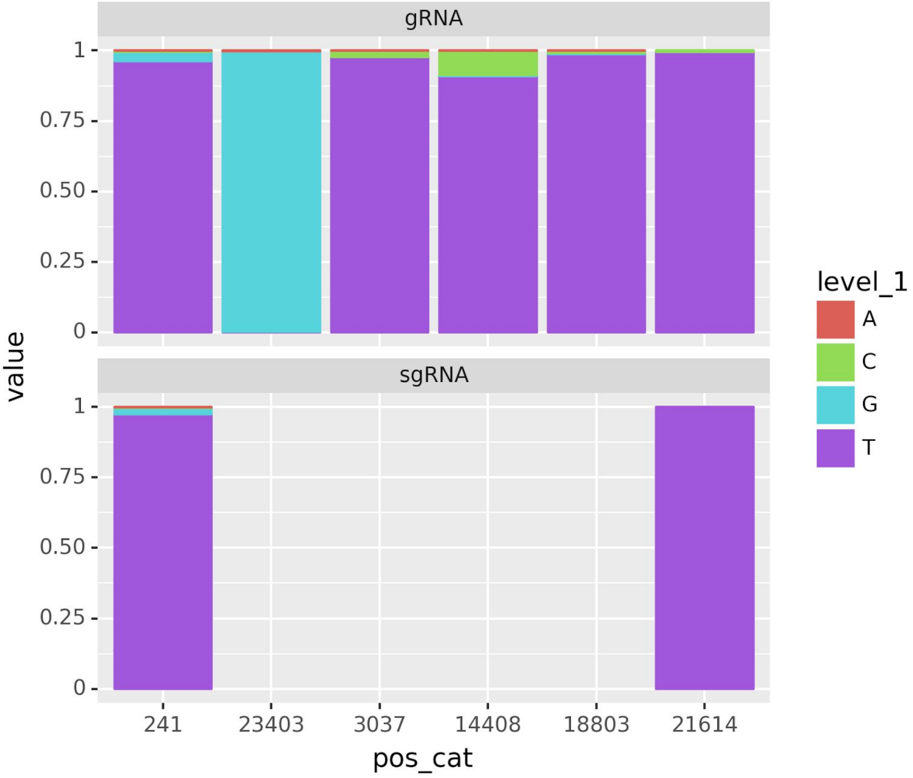


**Supplementary Figure S12 - Example of periscope output for variant analysis**

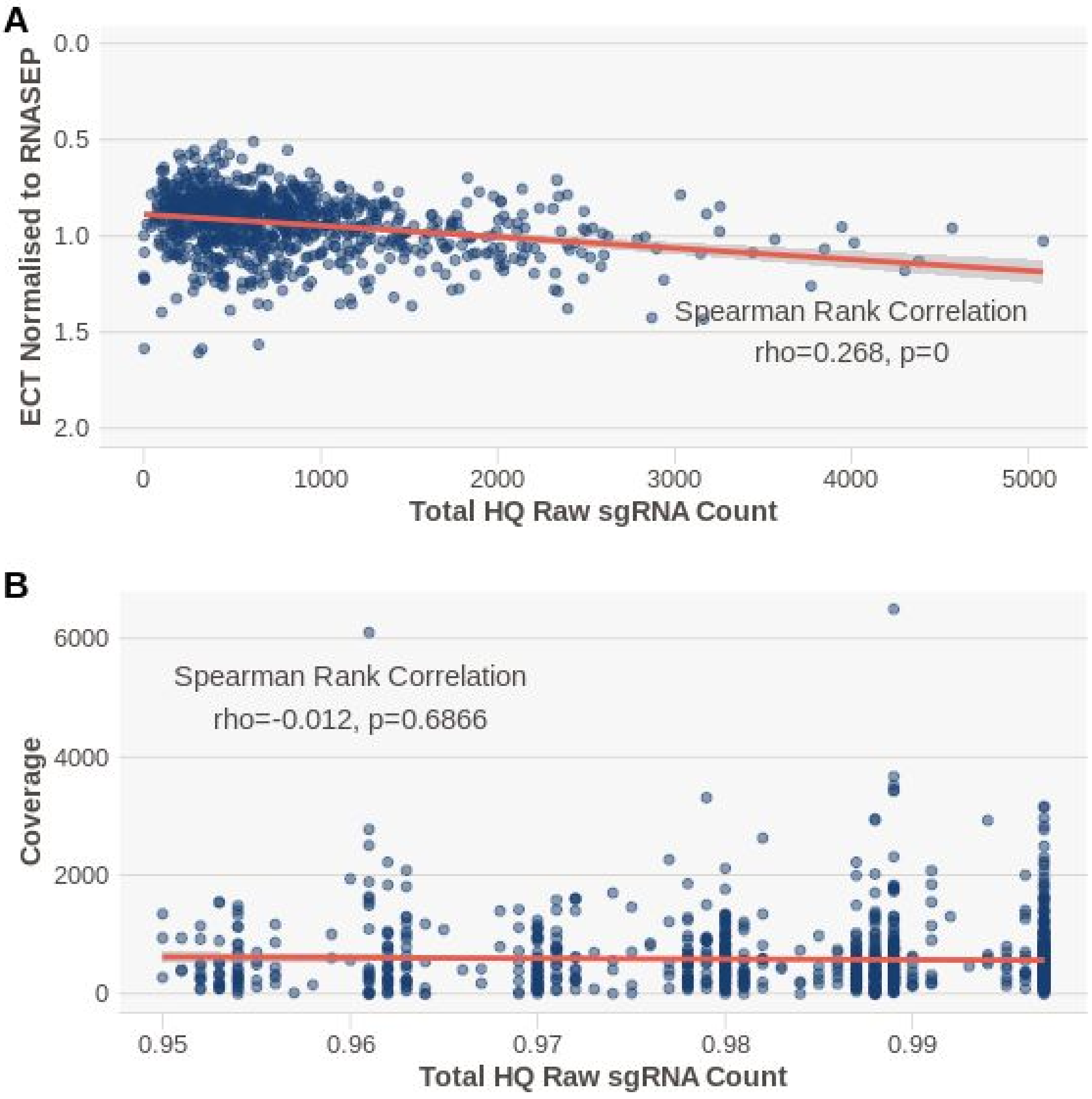


**Supplementary Figure S13 - Normalised E Ct and consensus coverage are not correlated with the raw amount of sub-genomic RNA detected**

**A.** Total raw HQ sgRNA counts are not correlated with normalised E Ct (E Ct/RNAseP). Y-axis reversed for ease of understanding as higher normalised E Ct = lower viral load. **B.** Total raw HQ sgRNA counts are not correlated with consensus genome coverage.
