## Supplementary File F19 for "periscope: sub-genomic RNA identification in SARS-CoV-2 Genomic Sequencing Data": environment.yml

```
name: periscope
channels:
  - conda-forge
  - bioconda
  - defaults
dependencies:
  - python=3.6
  - tqdm=4.46.0
  - plotnine=0.7.0
  - pandas=1.0.3
  - samtools=1.9
  - numpy=1.18.4
  - pysam=0.15.4
  - minimap2=2.17
  - bwa=0.7.17
  - bedtools=2.29.2
  - biopython=1.74
  - pybedtools=0.8.1
  - snakemake-minimal=5.13
  - pytest=5.4.2
  - pip=20.0.2
  - pip:
    - git+https://github.com/artic-network/fieldbioinformatics.git@3b6c1f3
```
