## Supplementary File F19 for "periscope: sub-genomic RNA identification in SARS-CoV-2 Genomic Sequencing Data": periscope.py

```
#!/usr/bin/env python3
from periscope import __version__
import argparse
import sys
import os
import snakemake
import glob
import logging

def main():

    parser = argparse.ArgumentParser(formatter_class=argparse.RawTextHelpFormatter,description='periscopre: Search for sgRNA reads in artic network SARS-CoV-2 sequencing data. A tool from Sheffield Bioinformatics Core/Florey Institute',usage='''periscope [options]''')
    parser.add_argument('--fastq-dir',dest='fastq_dir', help='the folder containing the raw pass demultiplexed fastqs from the artic protocol, if this is illumina data the tool expects a file labelled R1 and R2 in this dir.', default=None,required=False)
    parser.add_argument('--fastq',dest='fastq',help='if you already have a single fastq then you can use this flag instead, if illumina paired end separate fastq by space', nargs='+',required=False,default=[])
    parser.add_argument('--output-prefix',dest='output_prefix', help='Prefix of the output file',default="test")
    parser.add_argument('--score-cutoff',dest='score_cutoff', help='Cut-off for alignment score of leader (50)',default=50)
    parser.add_argument('--artic-primers', dest='artic_primers', help='artic network primer version used:\n* V1, V2, V3\n* 2kb (for the UCL longer amplicons)', default="V1")
    parser.add_argument('--threads', dest='threads', help='number of threads',
                        default="1")
    parser.add_argument('-r', '--resources', dest='resources', help="the path to the periscope resources directory - this is the place you cloned periscope into")
    parser.add_argument('-d', '--dry-run', action='store_true', help="perform a snakemake dryrun")
    parser.add_argument('-f', '--force', action='store_true', help="Overwrite all output", dest="force")
    parser.add_argument('--tmp',
                        help="pybedtools likes to write to /tmp if you want to write somewhere else define it here",
                        default="/tmp")
    parser.add_argument('--sample', help='sample id', default="SHEF-D2BD9")
    parser.add_argument('--technology', help='the sequencing technology used, either:\n*ont\n*illumina', default="ont")


    args = parser.parse_args()


    # if technology is illumina then we need to know where fastqs are because they could be paired end
    # this will work with any fastq input type
    if args.technology == "illumina":
        if len(args.fastq) == 0:
            print("If technology is illumina you must specify input fastqs wih --fastq flag. Do not use --fastq-dir", file=sys.stderr)
            exit(1)

    # check if fastq_dir exists
    print(args.fastq_dir)
    gzipped=False
    extension=".fastq"
    if args.fastq_dir:
        if not os.path.exists(args.fastq_dir):
            print("%s fastq directory must exist" % (args.fastq_dir), file=sys.stderr)
            exit(1)
        #here we should find out if fastqs are compressed or not - need to cat or zcat depending
        else:
            directory_listing = glob.glob(args.fastq_dir)
            if ".fq" in directory_listing:
                extension=".fq"
            if ".fq.gz" in directory_listing:
                gzipped=True
                extension=".fq.gz"
            if ".fastq.gz" in directory_listing:
                gzipped=True
                extension = ".fastq.gz"


    print(args.fastq)
    print(type(args.fastq))
    if len(args.fastq)>0:
        for fastq in args.fastq:
            if not os.path.exists(fastq):
                print("%s fastq file must exist" % (fastq), file=sys.stderr)
                exit(1)

    # check if version number is correct
    version = args.artic_primers
    if version not in ["V1", "V2", "V3","2kb"]:
        print("%s artic primer version incorrect" % (version), file=sys.stderr)
        exit(1)
    else:
        amplicons_bed="artic_amplicons_{}.bed".format(version)
        primers_bed="artic_primers_{}.bed".format(version)
        interest_bed = "artic_amplicons_of_interest.bed"


    # run snakemake pipeline 1st
    dir = os.path.join(os.path.dirname(__file__))
    scripts_dir= os.path.join(dir, 'scripts')


    config = dict(
        fastq_dir=args.fastq_dir,
        extension=extension,
        gzipped=gzipped,
        fastq=args.fastq,
        output_prefix=args.output_prefix,
        scripts_dir=scripts_dir,
        resources_dir=args.resources,
        amplicon_bed=amplicons_bed,
        interest_bed=interest_bed,
        primer_bed=primers_bed,
        orf_bed='orf_start.bed',
        score_cutoff=args.score_cutoff,
        reference_fasta='nCoV-2019.reference.fasta',
        sample=args.sample,
        threads=args.threads,
        tmp=args.tmp,
        technology=args.technology
    )


    snakefile = os.path.join(scripts_dir, 'Snakefile')
    print(snakefile)
    if not os.path.exists(snakefile):
        sys.stderr.write('Error: cannot find Snakefile at {}\n'.format(snakefile))
        sys.exit(-1)
    else:
        print("Found the snakefile")

    status = snakemake.snakemake(snakefile, printshellcmds=True,
                                 dryrun=args.dry_run, forceall=args.force, force_incomplete=True,
                                 config=config, cores=int(args.threads), lock=False
                                 )
    if status:  # translate "success" into shell exit code of 0
        exit(0)

    exit(1)
```
