## Supplementary File F19 for "periscope: sub-genomic RNA identification in SARS-CoV-2 Genomic Sequencing Data": calculate_amplicon_lengths.py

```
# bed format looks something like this
# MN908947.3	30	54	nCoV-2019_1_LEFT	nCoV-2019_1
# MN908947.3	385	410	nCoV-2019_1_RIGHT	nCoV-2019_1
import pprint as pp
from collections import OrderedDict
import argparse
import os
import sys


def closest(lst, K):
    return lst[min(range(len(lst)), key=lambda i: abs(lst[i] - K))]


def main(args):


    rights = {}
    rights_list=[]
    f = open(args.primer_bed, "r")
    for x in f:
        chr, start, end, name, name2 = x.rstrip().split("\t")
        if "RIGHT" in name:
            rights[int(end)]=name
            rights_list.append(int(end))
    f = open(args.orf_bed, "r")


    for x in f:
        chr,start,end,name=x.rstrip().split("\t")
        centre = (float(start)+float(end))/2
        closest_end = int(closest(rights_list,centre))
        distance = closest_end - centre
        if distance < 0:
            clean_list = [ x for x in rights_list if x is not closest_end ]
            closest_end = closest(clean_list,centre)
        else:
            clean_list=rights_list

        new_list=[ x for x in clean_list if x is not closest_end ]

        closest_end = int(closest(new_list, centre))
        distance = closest_end - centre
        if distance < 0:
            clean_list = [x for x in new_list if x is not closest_end]
            closest_end = closest(clean_list, centre)
            distance = closest_end - centre
            if distance < 0:
                clean_list = [x for x in clean_list if x is not closest_end]
                closest_end = closest(clean_list, centre)
                distance = closest_end - centre
        print(name+"\t"+str(distance))


    # f = open(args.input, "r")
    # result = OrderedDict()
    # for x in f:
    #     chr,start,end,name,name2=x.rstrip().split("\t")
    #     id = name.replace("_LEFT","").replace("_RIGHT","").replace("_alt2","")
    #     if id not in result:
    #         result[id]={}
    #         result[id]["start"]=start
    #     else:
    #         result[id]["end"]=end
    #
    # f = open(args.output, "w")
    # for i in result:
    #     f.write("\t".join(["MN908947.3",str(result[i]["start"]),str(result[i]["end"]),i])+"\n")
    # f.close()


if __name__ == '__main__':

    parser = argparse.ArgumentParser(formatter_class=argparse.RawTextHelpFormatter,description='periscopre: Prepare amplicon bed file, useful for visualisation\n\nYour bed file should look like this:\n\nMN908947.3	30	54	nCoV-2019_1_LEFT	nCoV-2019_1\nMN908947.3	385	410	nCoV-2019_1_RIGHT	nCoV-2019_1\n\nIt is important LEFT and RIGHT primers follow this naming scheme\n**AND** that the 5th column has <NAME>_1 or <NAME>_2 for the pool number')
    parser.add_argument('--primer_bed', help='primer bed file',default="The primers")
    parser.add_argument('--orf_bed', help='orf bed file', default="The orf starts")


    args = parser.parse_args()

    # if os.path.isfile(args.output):
    #     print("hello")
    #     # print("%s output file exists" % args.output, file=sys.stderr)
    #     exit(1)


    periscope = main(args)
```
