## Supplementary File F19 for "periscope: sub-genomic RNA identification in SARS-CoV-2 Genomic Sequencing Data": regenerate_counts_from_reads_tsv.py

```
import argparse
import periscope.scripts.search_for_sgRNA as p


class Read():

    def __init__(self,sample,read_id,position,read_length,orf,score,read_class,amplicon):
        self.sample = sample
        self.read_id = read_id
        self.reference_name = "MN908947.3"
        self.pos = position
        self.read_length = read_length
        self.orf = orf
        self.score = score
        self.read_class = read_class
        self.amplicon = amplicon


def reads_parser(read_file):

    with open(read_file, "r") as f:
        for line in f:
            sample,read_id,position,read_length,orf,score,read_class,amplicon=line.rstrip().split("\t")
            read=Read(sample,read_id,position,read_length,orf,score,read_class,amplicon)
            yield read
    # file looks like this
    # sample	read_id	position	read_length	orf	score	class	amplicon
    # SHEF-CAB89	acfa404a-736e-4c51-b809-648e0f5af7a4	14	497	None	46.0	gRNA	1


def main(args):
    orf_bed_object = p.open_bed(args.orf_bed)
    count=0
    for read in reads_parser(args.reads_file):
        count+=1
        print(count)
        print("old")
        print(read.orf)
        print("new")
        print(p.check_start(orf_bed_object,read))
        # if old is novel and change is to None - keep old ORF


if __name__ == '__main__':

    parser = argparse.ArgumentParser(description='periscopre: convert reads tsv into counts - useful for reprocessing with a different score')
    parser.add_argument('--output-prefix',dest='output_prefix', help='Prefix of the output file')
    parser.add_argument('--score-cutoff',dest='score_cutoff', help='Cut-off for alignment score of leader (45)',default=45)
    parser.add_argument('--orf-bed',dest="orf_bed", help='sample id')
    parser.add_argument('--sample', help='sample id')
    parser.add_argument('--reads-file',dest="reads_file", help='the reads file from your 1st periscope run')

    args = parser.parse_args()
    print(args.score_cutoff)

    main(args)
```
