## Supplementary File F19 for "periscope: sub-genomic RNA identification in SARS-CoV-2 Genomic Sequencing Data": regenerate_orf_summary_from_amplicons_tsv.py

```
import argparse
import periscope.scripts.search_for_sgRNA as p


class Amplicon():

    def __init__(self,sample,amplicon,raw_gRNA,total_reads,orf,raw_sgRNA,normalised_sgRNA,normalised_gRNA):
        self.sample = sample
        self.amplicon = amplicon
        self.raw_gRNA = raw_gRNA
        self.total_reads = total_reads
        self.orf = orf
        self.raw_sgRNA = raw_sgRNA
        self.normalised_sgRNA = normalised_sgRNA
        self.normalised_gRNA = normalised_gRNA


def amplicon_parser(amplicon_file):

    with open(amplicon_file, "r") as f:
        for line in f:
            if "sample" in line:
                continue
            sample,amplicon,raw_gRNA,total_reads,orf,raw_sgRNA,normalised_sgRNA,normalised_gRNA=line.rstrip().split("\t")
            amplicon=Amplicon(sample,amplicon,raw_gRNA,total_reads,orf,raw_sgRNA,normalised_sgRNA,normalised_gRNA)
            yield amplicon
    # file looks like this
    # sample	amplicon	raw_gRNA	total_reads	orf	raw_sgRNA	normalised_sgRNA	normalised_gRNA
    # SHEF-C764F	1	119	287	ORF1a	168	1.411764705882353	0.003114938617386069


def main(args):
    count=0
    orfs={"ORF1a": [1],"S" : [71],"ORF3a" : [84],"E" : [86,87],"M" : [87],"ORF6" : [89],"ORF7a" : [90,91],"ORF7b" : [91],"ORF8" : [92,93],"N" : [93],"N*" : [95],"ORF10" : [97,98]}
    orfs_amplicons = {}
    for amplicon in amplicon_parser(args.amplicon_file):
        if amplicon.orf == "NA":
            for orf in orfs:
                for amplicon_number in orfs[orf]:
                    if int(amplicon_number) == int(amplicon.amplicon):

                        amplicon.orf=orf

        if amplicon.orf in orfs:
            if amplicon.orf not in orfs_amplicons:
                orfs_amplicons[amplicon.orf]=[]
            orfs_amplicons[amplicon.orf].append(amplicon)

    # output new counts
    outfile_amplicons = args.output_prefix +"_periscope_amplicons_regenerated.tsv"
    outfile_counts = args.output_prefix + "_periscope_counts_regenerated.tsv"
    file_amplicons = open(outfile_amplicons, "w")
    file_counts = open(outfile_counts, "w")

    header_counts = ["sample","orf","normalised_sgRNA"]
    header_amplicons = ["sample","amplicon","raw_gRNA","total_reads","orf","raw_sgRNA","normalised_sgRNA","normalised_gRNA"]

    file_counts.write("\t".join(header_counts)+"\n")
    file_amplicons.write("\t".join(header_amplicons) + "\n")

    for orf in orfs_amplicons:
        line_counts=[]
        norm_orf = []
        for amplicon in orfs_amplicons[orf]:
            line_orf=[]
            line_orf.append(args.sample)
            line_orf.append(amplicon.amplicon)
            line_orf.append(amplicon.raw_gRNA)
            line_orf.append(amplicon.total_reads)
            line_orf.append(amplicon.orf)
            line_orf.append(amplicon.raw_sgRNA)
            line_orf.append(amplicon.normalised_sgRNA)
            line_orf.append(amplicon.normalised_gRNA)

            file_amplicons.write("\t".join(line_orf)+"\n")

            norm_orf.append(float(amplicon.normalised_sgRNA))

        line_counts.append(args.sample)
        line_counts.append(str(orf))
        # sum proportion of sgRNA_normalised from multiple amplicons
        line_counts.append(str(sum(norm_orf)))

        file_counts.write("\t".join(line_counts)+"\n")


    file_counts.close()
    file_amplicons.close()


if __name__ == '__main__':

    parser = argparse.ArgumentParser(description='periscopre: convert reads tsv into counts - useful for reprocessing with a different score')
    parser.add_argument('--output-prefix',dest='output_prefix', help='Prefix of the output file')
    parser.add_argument('--sample', help='sample id')
    parser.add_argument('--amplicon-file',dest="amplicon_file", help='the reads file from your 1st periscope run')

    args = parser.parse_args()

    main(args)
```
