## Supplementary File F19 for "periscope: sub-genomic RNA identification in SARS-CoV-2 Genomic Sequencing Data": search_for_sgRNA_illumina.py

```
#!/usr/bin/env python3
from periscope import __version__

from Bio import pairwise2
import pysam
import argparse
from pybedtools import *
from artic.vcftagprimersites import read_bed_file
import sys
from numpy import median
from tqdm import tqdm

class ClassifiedRead():
    def __init__(self,sgRNA: bool,orf: str,read: pysam.AlignedRead):
        self.sgRNA = sgRNA
        self.orf = orf
        self.pos = read.pos
        self.read = read

def get_mapped_reads(bam):
    mapped_reads = int(pysam.idxstats(bam).split("\n")[0].split("\t")[2])
    return mapped_reads

def check_start(bed_object,read):
    """
    find out where the read is in a bed file, in this case the ORF starts
    :param bed_object: bedtools object
    :param read: pysam read object
    :return: the orf
    """

    # reads with a pos of 0 make this fail so puting in try except works
    try:
        read_feature = BedTool(read.reference_name + "\t" + str(read.pos) + "\t" + str(read.pos), from_string=True)
        intersect = bed_object.intersect(read_feature)
        orf=intersect[0].name
        # if len(intersect) > 1:
        #     print("odd")
    except:
        orf=None
    # remove bedtools objects from temp
    cleanup()
    return orf


def supplementary_method(read):
   # we don't need the supplementary alignment
    if read.is_supplementary:
        return "supplementary"

    for tag in read.get_tags():
        if tag[0] == 'SA':
            supp_chrom,supp_pos,supp_strand,supp_cigar,supp_quality,unknown = tag[1].split(",")
            print(supp_pos)
            if 67 >= int(supp_pos) >= 0:
                return "supplementary_maps_to_leader"


def extact_soft_clipped_bases(read):
    """
            +-----+--------------+-----+
            |M    |BAM_CMATCH    |0    |
            +-----+--------------+-----+
            |I    |BAM_CINS      |1    |
            +-----+--------------+-----+
            |D    |BAM_CDEL      |2    |
            +-----+--------------+-----+
            |N    |BAM_CREF_SKIP |3    |
            +-----+--------------+-----+
            |S    |BAM_CSOFT_CLIP|4    |
            +-----+--------------+-----+
            |H    |BAM_CHARD_CLIP|5    |
            +-----+--------------+-----+
            |P    |BAM_CPAD      |6    |
            +-----+--------------+-----+
            |=    |BAM_CEQUAL    |7    |
            +-----+--------------+-----+
            |X    |BAM_CDIFF     |8    |
            +-----+--------------+-----+
            |B    |BAM_CBACK     |9    |
            +-----+--------------+-----+
    :param cigar:
    :return:
    """


    cigar = read.cigartuples
    if cigar[0][0] == 4:
        # there is softclipping on 5' end
        number_of_bases_sclipped = cigar[0][1]
        # get 3 extra bases incase of homology
        bases_sclipped = read.seq[0:number_of_bases_sclipped+3]


        search='AACCAACTTTCGATCTCTTGTAGATCTGTTCTC'


        # not enough bases to determine leader
        if number_of_bases_sclipped < 6:
            return False

        # determine perfect score for a match
        if number_of_bases_sclipped >= 33:
            # allow 3 mistamches
            perfect = 60.0
        else:
            # allow 1 mistamches
            perfect = (number_of_bases_sclipped * 2)-2

        align = pairwise2.align.localms(bases_sclipped, search, 2, -2, -20, -.1,  one_alignment_only=True  )
        align_score=align[0][2]
        align_right_position = align[0][4]

        logger.debug("Processing read: %s", read.query_name)
        logger.debug("%s perfect score: %s", read.query_name,str(perfect))
        logger.debug("%s actual score: %s", read.query_name, str(align_score))
        logger.debug("%s alignment: %s", read.query_name, align)

        # position of alignment must be all the way to the right
        if align_right_position >= len(search):
            # allow only one mismtach - Mismatches already taken care of above?
            # if perfect-align_score <= 2:
            if perfect-align_score <= 0:
                logger.debug("%s sgRNA: %s", read.query_name, True)
                return True
            else:
                # more than allowed number of mismatches
                logger.debug("%s sgRNA: %s,%s", read.query_name, False,'too many mismatches')
                return False
        else:
            # match is not at the end of the leader
            logger.debug("%s sgRNA: %s,%s", read.query_name, False,'match not at end of leader')
            return False
    else:
        # No soft clipping at 5' end of read
        logger.debug("%s sgRNA: %s,%s", read.query_name, False,'no soft-clipping')
        return False


def get_coverage(start,end,inbamfile):
    coverage = []
    for pileupcolumn in inbamfile.pileup("MN908947.3", int(start), int(end)):
            coverage.append(pileupcolumn.n)
    return median(coverage)


def open_bed(bed):
    """
    open bed file and return a bedtools object
    :param bed:
    :return:
    """
    bed_object = BedTool(bed)
    return bed_object


def setup_counts(primer_bed_object):
    """
    make the main counts dictionary, we populate this as we loop through the reads in teh bam file
    :param primer_bed_object: primer bed file object needed to get the pool name
    :return:
    """
    # set up dictionary for normalisation
    # need to get all regions in bed and make into a dict
    # { 71: { total_reads: x, genomic_reads: y, sg_reads: {orf:z,orf2:k},'normalised_sgRNA': {orf:i,orf2:t} } }
    total_counts = {}
    for primer in primer_bed_object:
        amplicon = int(primer["Primer_ID"].split("_")[1])
        if amplicon not in total_counts:
            total_counts[amplicon] = {'pool': primer["PoolName"], 'total_reads': 0, 'gRNA': [], 'sgRNA_HQ': {}, 'sgRNA_LQ':{}, 'sgRNA_LLQ':{}, 'nsgRNA_HQ':{}, 'nsgRNA_LQ':{}}
    return total_counts


def main(args):

    # read input bam file
    inbamfile = pysam.AlignmentFile(args.bam, "rb")

    # get bam header so that we can use it for writing later
    bam_header = inbamfile.header.copy().to_dict()
    # open output bam with the header we just got
    outbamfile = pysam.AlignmentFile(args.output_prefix + "_periscope.bam", "wb", header=bam_header)

    # get mapped reads
    mapped_reads = get_mapped_reads(args.bam)

    # open the orfs bed file
    orf_bed_object = open_bed(args.orf_bed)

    orf_coverage={}
    # get coverage for each orf
    logger.warning("getting coverage at canonical ORF sites")
    for row in orf_bed_object:

        median=get_coverage(row.start,row.end,inbamfile)

        orf_coverage[row.name]=median
    logger.warning("getting coverage at canonical ORF sites....DONE")

    # read input bam file again
    inbamfile = pysam.AlignmentFile(args.bam, "rb")

    # open the artic primer bed file
    primer_bed_object=read_bed_file(args.primer_bed)
    # set the output reads filename
    # outfile_reads = args.output_prefix + "_periscope_reads.tsv"
    # set the output counts file name

    # add headers to these files
    # file_reads = open(outfile_reads, "w")
    # file_reads.write("sample\tread_id\tposition\tread_length\torf\tscore\tclass\tamplicon\n")

    total_counts = setup_counts(primer_bed_object)
    # for every read let's decide if it's sgRNA or not
    logger.warning("Processing " + str(mapped_reads) + " reads")
    result={}
    count=0
    orfs={}
    reads={}
    for read in tqdm(inbamfile,total=mapped_reads):

        if read.seq == None:
            # print("%s read has no sequence" %
            #       (read.query_name), file=sys.stderr)
            continue
        if read.is_unmapped:
            # print("%s skipped as unmapped" %
            #       (read.query_name), file=sys.stderr)
            continue
        if read.is_supplementary:
            # print("%s skipped as supplementary" %
            #       (read.query_name), file=sys.stderr)
            continue
        if read.is_secondary:
            # print("%s skipped as secondary" %
            #       (read.query_name), file=sys.stderr)
            continue
        # print("------")
        # print(read.query_name)
        # # print(read.is_read1)
        # # print(read.get_tags())
        # print(read.cigar)
        leader_search_result = extact_soft_clipped_bases(read)
        if read.query_name not in reads:
            reads[read.query_name] = []
        orf=None
        for row in orf_bed_object:
            # see if read falls within ORF start location
            if row.end >= read.pos >= row.start:
                orf = row.name
        if orf == None:
            if leader_search_result == True:
                orf = "novel_" + str(read.pos)


        reads[read.query_name].append(

            ClassifiedRead(sgRNA=leader_search_result,orf=orf,read=read)


        )


    # now we have all the reads classified, deal with pairs
    logger.warning("Processing " + str(mapped_reads) + " reads....DONE")
    logger.info("dealing with read pairs")
    for id,pair in reads.items():

        # get the class and orf of the left hand read, this will be the classification and ORF for the pair - sometimes right read looks like it has subgenomic evidence - there are likely false positives

        left_read = min(pair, key=lambda x: x.pos)

        right_read = max(pair, key=lambda x: x.pos)

        read_class = left_read.sgRNA
        orf = left_read.orf

        left_read_object = left_read.read
        right_read_object = right_read.read

        left_read_object.set_tag('XO', 'orf')
        right_read_object.set_tag('XO', 'orf')

        if read_class == True:
            left_read_object.set_tag('XC', 'sgRNA')
            right_read_object.set_tag('XC', 'sgRNA')
        else:
            left_read_object.set_tag('XC', 'gRNA')
            right_read_object.set_tag('XC', 'gRNA')


        outbamfile.write(left_read_object)
        outbamfile.write(right_read_object)

        if orf == None:
            continue

        if read_class == False:
            continue

        if orf not in orfs:
            orfs[orf] = [left_read_object]
        else:
            orfs[orf].append(left_read_object)

    logger.info("dealing with read pairs....DONE")

    outbamfile.close()
    pysam.sort("-o", args.output_prefix + "_periscope_sorted.bam",  args.output_prefix + "_periscope.bam")
    pysam.index(args.output_prefix + "_periscope_sorted.bam")

    novel_count=0
    canonical = open(args.output_prefix+"_periscope_counts.csv","w")
    canonical.write(",".join(["sample","mapped_reads","orf","sgRNA_count","coverage", "sgRPTL","sgRPHT\n"]))

    novel = open(args.output_prefix+"_periscope_novel_counts.csv","w")
    novel.write(",".join(["sample","mapped_reads", "orf", "sgRNA_count", "coverage", "sgRPTL","sgRPHT\n"]))

    logger.info("summarising results")

    for orf in orfs:
        sgRPHT = len(orfs[orf]) / (mapped_reads / 10000)
        if "novel" not in orf:
            sgRPTL = len(orfs[orf])/(orf_coverage[orf]/1000)
            canonical.write(args.sample+","+str(mapped_reads)+","+orf+","+str(len(orfs[orf]))+","+str(orf_coverage[orf])+","+str(sgRPTL)+","+str(sgRPHT)+"\n")
        else:
            position = int(orf.split("_")[1])
            coverage=get_coverage(position-20,position+20,inbamfile)
            sgRPTL = len(orfs[orf])/(coverage/1000)
            novel.write(args.sample+","+str(mapped_reads)+","+orf+","+str(len(orfs[orf]))+","+str(coverage)+","+str(sgRPTL)+","+str(sgRPHT)+"\n")
            novel_count+=len(orfs[orf])

    canonical.close()
    novel.close()

    logger.info("summarising results....DONE")

    amplicons = open(args.output_prefix + "_periscope_amplicons.csv", "w")
    amplicons.write("not used yet")
    amplicons.close()


if __name__ == '__main__':


    parser = argparse.ArgumentParser(description='periscopre: Search for sgRNA reads in artic network SARS-CoV-2 sequencing data')
    parser.add_argument('--bam', help='bam file',default="The bam file of full artic reads")
    parser.add_argument('--output-prefix',dest='output_prefix',help="Path to the output, e.g. <DIR>/<SAMPLE_NAME>")
    parser.add_argument('--score-cutoff',dest='score_cutoff', help='Cut-off for alignment score of leader (50) we recommend you leave this at 50',default=50)
    parser.add_argument('--orf-bed', dest='orf_bed', help='The bed file with ORF start positions')
    parser.add_argument('--primer-bed', dest='primer_bed', help='The bed file with artic primer positions')
    parser.add_argument('--amplicon-bed', dest='amplicon_bed', help='A bed file of artic amplicons')
    parser.add_argument('--sample', help='sample id',default="SAMPLE")
    parser.add_argument('--tmp',help="pybedtools likes to write to /tmp if you want to write somewhere else define it here",default="/tmp")
    parser.add_argument('--progress', help='display progress bar', default="")


    logger = logging
    logger.basicConfig(format='%(asctime)s - %(message)s', level=logging.INFO)

    logger.info("staring periscope")

    args = parser.parse_args()

    set_tempdir(args.tmp)

    periscope = main(args)

    if periscope:
        print("all done", file=sys.stderr)
```
