## Supplementary File F19 for "periscope: sub-genomic RNA identification in SARS-CoV-2 Genomic Sequencing Data": search_for_sgRNA_ont.py

```
#!/usr/bin/env python3
from periscope import __version__

from Bio import pairwise2
import pysam
import argparse
from pybedtools import *
import datetime
from artic.align_trim import find_primer
from artic.vcftagprimersites import read_bed_file
import sys
import os
import pprint as pp
import snakemake
from collections import namedtuple
class PeriscopeRead(object):
    def __init__(self, read):
        self.read = read

import time
from tqdm import tqdm

def get_mapped_reads(bam):
    mapped_reads = int(pysam.idxstats(bam).split("\n")[0].split("\t")[2])
    return mapped_reads

def check_start(bed_object,read):
    """
    find out where the read is in a bed file, in this case the ORF starts
    :param bed_object: bedtools object
    :param read: pysam read object
    :return: the orf
    """
    # reads with a pos of 0 make this fail so puting in try except works
    # try:
    for row in bed_object:
        # see if read falls within ORF start location
        if row.end >= read.pos >= row.start:
            orf = row.name
            break
        else:
            orf=None

        # read_feature = BedTool(read.reference_name + "\t" + str(read.pos) + "\t" + str(read.pos), from_string=True)
        # intersect = bed_object.intersect(read_feature)
        # orf=intersect[0].name
        # if len(intersect) > 1:
        #     print("odd")
    # except:
    #     orf=None
    # remove bedtools objects from temp
    # cleanup()
    return orf

def search_reads(read,search):
    """
    given a pysam read object and a search string perform a localms alignment
    :param read: pysam read object
    :param search: DNA search string e.g. ATGTGCTTGATGC
    :return: dictionary containing the read_id, alignment score and the position of the read
    """
    align_score = pairwise2.align.localms(search, read.seq, 2, -2, -10, -.1,score_only=True)

    return {
        "read_id":  read.query_name,
        "align_score": align_score,
        "read_position": read.pos,
        "sequence": read.seq
    }


def find_amplicon(read,primer_bed_object):
    """
    use artic code to find primers called "find_primers"
    returns (1, 1, {'chrom': 'MN908947.3', 'start': 21357, 'end': 21386, 'Primer_ID': 'nCoV-2019_71_LEFT', 'PoolName': 'nCoV-2019_1', 'direction': '+'})
    in the case of sgRNAs this code fails to find the correct primer for the + direction. This is actualy a clue that
    this read is an sgRNA

    :param read:
    :param primer_bed_object:
    :return: the amplicon of the read
    """


    # get the left primer

    left_primer = find_primer(primer_bed_object, read.reference_start, '+')


    # get the right primer
    right_primer = find_primer(primer_bed_object, read.reference_end, '-')


    # get the left primer amplicon (we don't actually use this)
    left_amplicon = int(left_primer[2]['Primer_ID'].split("_")[1])
    # get the right primer amplicon
    right_amplicon = int(right_primer[2]['Primer_ID'].split("_")[1])

    # WARNING - LEFT_AMPLICON IS NOT RELIABLE FOR SG_RNA

    return dict(left_amplicon=left_amplicon,left_primer=left_primer,right_amplicon=right_amplicon,right_primer=right_primer)

def classify_read(read,score,score_cutoff,orf,amplicons):
    """
    classify read based on leader alignment score and other metrics
    :param score: the score
    :param score_cutoff: the user provided cut-off
    :return:
    """

    # some things I've learnt:
    # - if amplicons match it's more likely to by a gRNA but that doesn't hold true for reads that span amplicons - so score should still be 1st port of call
    # print(amplicons)
    quality=None
    if score > int(score_cutoff):
        quality = "HQ"
        if orf is not None:
            read_class = "sgRNA"
        else:
            read_class = "nsgRNA"

    elif score > 30:
        quality = "LQ"
        if orf is not None:
            read_class = "sgRNA"
        else:
            read_class = "nsgRNA"

    else:
        if orf is not None:
            quality = "LLQ"
            read_class = "sgRNA"
        else:
            read_class = "gRNA"

    # for those that have been classified as nsgRNA - do a final check - check not at amplicon edge
    # we see a lot of false positives at read ends

    if read_class == "nsgRNA":
        primer_start = amplicons["left_primer"][2]["start"]-5
        primer_end = amplicons["left_primer"][2]["end"]+5
        if primer_start <= read.pos <= primer_end:
            quality=None
            read_class="gRNA"

    if quality:
        return read_class+"_"+quality
    else:
        return read_class


def open_bed(bed):
    """
    open bed file and return a bedtools object
    :param bed:
    :return:
    """
    bed_object = BedTool(bed)
    return bed_object


def setup_counts(primer_bed_object):
    """
    make the main counts dictionary, we populate this as we loop through the reads in teh bam file
    :param primer_bed_object: primer bed file object needed to get the pool name
    :return:
    """
    # set up dictionary for normalisation
    # need to get all regions in bed and make into a dict
    # { 71: { total_reads: x, genomic_reads: y, sg_reads: {orf:z,orf2:k},'normalised_sgRNA': {orf:i,orf2:t} } }
    total_counts = {}
    for primer in primer_bed_object:
        amplicon = int(primer["Primer_ID"].split("_")[1])
        if amplicon not in total_counts:
            total_counts[amplicon] = {'pool': primer["PoolName"], 'total_reads': 0, 'gRNA': [], 'sgRNA_HQ': {}, 'sgRNA_LQ':{}, 'sgRNA_LLQ':{}, 'nsgRNA_HQ':{}, 'nsgRNA_LQ':{}}
    return total_counts


def calculate_normalised_counts(mapped_reads,total_counts,outfile_amplicon,orf_bed_object):
    """
    calculate normalised read counts on a per amplicon bases

    :param mapped_reads: total mapped reads
    :param total_counts: the total counts dictionary
    :param outfile_amplicon: the amplicon outfile
    :param orf_bed_object: the orf bed file object
    :return: the total counts dictionary with normalisation added
    """
    done=[]
    with open(outfile_amplicon, "w") as f:
        header = ["sample", "amplicon", "mapped_reads", "orf", "quality", "gRNA_count", "gRPTH", "sgRNA_count", "sgRPHT",
              "sgRPTg"]
        f.write(",".join(header)+"\n")
        for amplicon in total_counts:

            # total count of gRNA for amplicon
            amplicon_gRNA_count = len(total_counts[amplicon]["gRNA"])

            # gRNA total count per 100l mapped reads
            amplicon_gRPTH = amplicon_gRNA_count / (mapped_reads / 100000)

            total_counts[amplicon]["gRPHT"] = {}

            for quality in ["HQ", "LQ", "LLQ"]:
                total_counts[amplicon]["sgRPHT_" + quality] = {}
                total_counts[amplicon]["sgRPTg_" + quality] = {}
                for orf in total_counts[amplicon]["sgRNA_" + quality]:
                    total_counts[amplicon]["gRPHT"][orf] = amplicon_gRPTH

                    amplicon_orf_sgRNA_count = len(total_counts[amplicon]["sgRNA_" + quality][orf])

                    # normalised per 100k total mapped reads
                    amplicon_orf_sgRPHT = amplicon_orf_sgRNA_count / (mapped_reads / 100000)

                    total_counts[amplicon]["sgRPHT_" + quality][orf] = amplicon_orf_sgRPHT

                    # normalised per 1000 gRNA reads from this amplicon
                    try:
                        amplicon_orf_sgRPTg = amplicon_orf_sgRNA_count / (amplicon_gRNA_count / 1000)
                    except:
                        amplicon_orf_sgRPTg = "NA"


                    total_counts[amplicon]["sgRPTg_" + quality][orf] = amplicon_orf_sgRPTg

                    line = []
                    line.append(args.sample)
                    line.append(str(amplicon))
                    line.append(str(mapped_reads))
                    line.append(str(orf))
                    line.append(str(quality))
                    line.append(str(amplicon_gRNA_count))
                    line.append(str(amplicon_gRPTH))
                    line.append(str(amplicon_orf_sgRNA_count))
                    line.append(str(amplicon_orf_sgRPHT))
                    line.append(str(amplicon_orf_sgRPTg))
                    f.write(",".join(line)+"\n")

            for quality in ["HQ", "LQ"]:
                total_counts[amplicon]["nsgRPHT_" + quality] = {}
                total_counts[amplicon]["nsgRPTg_" + quality] = {}
                for orf in total_counts[amplicon]["nsgRNA_" + quality]:
                    total_counts[amplicon]["gRPHT"][orf] = amplicon_gRPTH

                    amplicon_orf_sgRNA_count = len(total_counts[amplicon]["nsgRNA_" + quality][orf])

                    # normalised per 100k total mapped reads
                    amplicon_orf_sgRPHT = amplicon_orf_sgRNA_count / (mapped_reads / 100000)

                    total_counts[amplicon]["nsgRPHT_" + quality][orf] = amplicon_orf_sgRPHT

                    # normalised per 1000 gRNA reads from this amplicon
                    amplicon_orf_sgRPTg = amplicon_orf_sgRNA_count / (amplicon_gRNA_count / 1000)

                    total_counts[amplicon]["nsgRPTg_" + quality][orf] = amplicon_orf_sgRPTg

                    line = []
                    line.append(args.sample)
                    line.append(str(amplicon))
                    line.append(str(mapped_reads))
                    line.append(str(orf))
                    line.append(str(quality))
                    line.append(str(amplicon_gRNA_count))
                    line.append(str(amplicon_gRPTH))
                    line.append(str(amplicon_orf_sgRNA_count))
                    line.append(str(amplicon_orf_sgRPHT))
                    line.append(str(amplicon_orf_sgRPTg))
                    f.write(",".join(line)+"\n")

                    # read_feature = BedTool("MN908947.3" + "\t" + str(int(orf.split("_")[1])-1) + "\t" + str(orf.split("_")[1]) + "\t" + str(orf),
                    #                        from_string=True)
                    if str(orf) not in done:
                        # fixes bug where if the read maps to 0 end up with -1 as a position
                        if int(orf.split("_")[1]) == 0:
                            read_feature = BedTool("MN908947.3" + "\t" + str(int(orf.split("_")[1])) + "\t" + str(
                                orf.split("_")[1]) + "\t" + str(orf), from_string=True)
                        else:
                            read_feature = BedTool("MN908947.3" + "\t" + str(int(orf.split("_")[1]) - 1) + "\t" + str(orf.split("_")[1]) + "\t" + str(orf),from_string=True)
                        orf_bed_object = orf_bed_object.cat(read_feature,postmerge=False)
                        done.append(str(orf))


    f.close()
    # orf_bed_object=orf_bed_object.sort().merge(c=4,o="distinct")
    return total_counts,orf_bed_object


def summarised_counts_per_orf(total_counts,orf_bed_object):
    """
    summarise counts per ORF

    sumarise the counts per ORF
    :param total_counts: the total counts dictionary created by calculate_normalised_counts
    :param orf_bed_object: the orf bed file object
    :return: a final dictionary of counts and norm counts per ORF
    """
    result = {}
    for orf in orf_bed_object:
        if orf.name not in result:
            result[orf.name] = {}
            result[orf.name]["gRPHT"] = 0
            result[orf.name]["amplicons"] = []
            result[orf.name]["gRNA_count"] = 0
            if "novel" in orf.name:
                for quality in ["LQ", "HQ"]:
                    result[orf.name]["nsgRNA_" + quality + "_count"] = 0
            else:
                for quality in ["LLQ", "LQ", "HQ"]:
                    result[orf.name]["sgRNA_" + quality + "_count"] = 0

        for amplicon in total_counts:
            if orf.name in total_counts[amplicon]["gRPHT"]:
                result[orf.name]["gRPHT"] += total_counts[amplicon]["gRPHT"][orf.name]
                result[orf.name]["amplicons"].append(str(amplicon))
                result[orf.name]["gRNA_count"] += len(total_counts[amplicon]["gRNA"])
            if "novel" in orf.name:
                for quality in ["LQ", "HQ"]:
                    if orf.name in total_counts[amplicon]["nsgRNA_" + quality]:
                        result[orf.name]["nsgRNA_" + quality + "_count"] += len(
                            total_counts[amplicon]["nsgRNA_" + quality][orf.name])

                    for metric in ["nsgRPHT", "nsgRPTg"]:
                        qmetric = metric + "_" + quality
                        if qmetric not in result[orf.name]:
                            result[orf.name][qmetric] = 0
                        if orf.name in total_counts[amplicon][qmetric]:
                            result[orf.name][qmetric] += total_counts[amplicon][qmetric][orf.name]

            else:
                for quality in ["LLQ", "LQ", "HQ"]:
                    if orf.name in total_counts[amplicon]["sgRNA_" + quality]:
                        result[orf.name]["sgRNA_" + quality + "_count"] += len(total_counts[amplicon]["sgRNA_" + quality][orf.name])

                    for metric in ["sgRPHT", "sgRPTg"]:
                        qmetric = metric + "_" + quality
                        if qmetric not in result[orf.name]:
                            result[orf.name][qmetric] = 0
                        if orf.name in total_counts[amplicon][qmetric]:
                            try:
                                result[orf.name][qmetric] += total_counts[amplicon][qmetric][orf.name]
                            except:
                                result[orf.name][qmetric] = "NA"
    return result

def output_summarised_counts(mapped_reads,result,outfile_counts,outfile_counts_novel):
    """
    output the summarised counts from summarised_counts_per_orf

    :param mapped_reads: mapped read count
    :param result: the result dictionary created by summarised_counts_per_orf
    :param outfile_counts: the outfile for the counts
    :param outfile_counts_novel: the outfile for the novel counts
    """
    with open(outfile_counts,"w") as f:
        header = ["sample", "orf", "mapped_reads", "amplicons","gRNA_count", "sgRNA_HQ_count", "sgRNA_LQ_count", "sgRNA_LLQ_count", "gRHPT", "sgRPTg_HQ", "sgRPTg_LQ", "sgRPTg_LLQ",
                  "sgRPTg_ALL", "sgRPHT_HQ", "sgRPHT_LQ", "sgRPHT_LLQ", "sgRPHT_ALL"]
        f.write(",".join(header)+"\n")
        for orf in result:
            if "novel" not in orf:
                # construct output line
                line = []
                line.append(args.sample)
                line.append(orf)
                line.append(str(mapped_reads))
                line.append("|".join(result[orf]["amplicons"]))
                line.append(str(result[orf]["gRNA_count"]))
                line.append(str(result[orf]["sgRNA_HQ_count"]))
                line.append(str(result[orf]["sgRNA_LQ_count"]))
                line.append(str(result[orf]["sgRNA_LLQ_count"]))
                line.append(str(result[orf]["gRPHT"]))
                line.append(str(result[orf]["sgRPTg_HQ"]))
                line.append(str(result[orf]["sgRPTg_LQ"]))
                line.append(str(result[orf]["sgRPTg_LLQ"]))
                try:
                    sgRPTg_all = sum([result[orf]["sgRPTg_HQ"], result[orf]["sgRPTg_LQ"], result[orf]["sgRPTg_LLQ"]])
                except:
                    sgRPTg_all = "NA"
                line.append(str(sgRPTg_all))
                line.append(str(result[orf]["sgRPHT_HQ"]))
                line.append(str(result[orf]["sgRPHT_LQ"]))
                line.append(str(result[orf]["sgRPHT_LLQ"]))
                try:
                    sgRPHT_all = sum([result[orf]["sgRPHT_HQ"], result[orf]["sgRPHT_LQ"], result[orf]["sgRPHT_LLQ"]])
                except:
                    sgRPHT_all = "NA"
                line.append(str(sgRPHT_all))

                f.write(",".join(line) + "\n")
        f.close()
    # deal with novel sgRNA seperatley
    with open(outfile_counts_novel, "w") as f:
        novel_header = ["sample", "orf", "mapped_reads", "amplicons", "gRNA_count", "nsgRNA_HQ_count", "nsgRNA_LQ_count",
                        "gRHPT", "nsgRPTg_HQ", "nsgRPTg_LQ", "nsgRPTg_ALL", "nsgRPHT_HQ", "nsgRPHT_LQ", "nsgRPHT_ALL"]
        f.write(",".join(novel_header) + "\n")
        for orf in result:
            if "novel" in orf:
                # construct output line
                line = []
                line.append(args.sample)
                line.append(orf)
                line.append(str(mapped_reads))
                line.append("|".join(result[orf]["amplicons"]))
                line.append(str(result[orf]["gRNA_count"]))
                line.append(str(result[orf]["nsgRNA_HQ_count"]))
                line.append(str(result[orf]["nsgRNA_LQ_count"]))
                line.append(str(result[orf]["gRPHT"]))
                line.append(str(result[orf]["nsgRPTg_HQ"]))
                line.append(str(result[orf]["nsgRPTg_LQ"]))
                try:
                    nsgRPTg_all = sum([result[orf]["nsgRPTg_HQ"], result[orf]["nsgRPTg_LQ"]])
                except:
                    nsgRPTg_all = "NA"
                line.append(str(nsgRPTg_all))
                line.append(str(result[orf]["nsgRPHT_HQ"]))
                line.append(str(result[orf]["nsgRPHT_LQ"]))
                try:
                    nsgRPHT_all = sum([result[orf]["nsgRPHT_HQ"], result[orf]["nsgRPHT_LQ"]])
                except:
                    nsgRPTg_all = "NA"
                line.append(str(nsgRPHT_all))

                f.write(",".join(line) + "\n")
        f.close()


def main(args):

    # read input bam file
    inbamfile = pysam.AlignmentFile(args.bam, "rb")
    # get bam header so that we can use it for writing later
    bam_header = inbamfile.header.copy().to_dict()
    # open output bam with the header we just got
    outbamfile = pysam.AlignmentFile(args.output_prefix + "_periscope.bam", "wb", header=bam_header)

    # get mapped reads
    mapped_reads = get_mapped_reads(args.bam)

    # open the orfs bed file
    orf_bed_object = open_bed(args.orf_bed)
    # open the artic primer bed file
    primer_bed_object=read_bed_file(args.primer_bed)
    # set the output reads filename
    # outfile_reads = args.output_prefix + "_periscope_reads.tsv"
    # set the output counts file name

    # add headers to these files
    # file_reads = open(outfile_reads, "w")
    # file_reads.write("sample\tread_id\tposition\tread_length\torf\tscore\tclass\tamplicon\n")

    total_counts = setup_counts(primer_bed_object)
    # for every read let's decide if it's sgRNA or not
    print("Processing " + str(mapped_reads) + " reads", file=sys.stderr)
    for read in tqdm(inbamfile,total=mapped_reads):

        if read.seq == None:
            # print("%s read has no sequence" %
            #       (read.query_name), file=sys.stderr)
            continue
        if read.is_unmapped:
            # print("%s skipped as unmapped" %
            #       (read.query_name), file=sys.stderr)
            continue
        if read.is_supplementary:
            # print("%s skipped as supplementary" %
            #       (read.query_name), file=sys.stderr)
            continue

        # find the amplicon for the read

        amplicons = find_amplicon(read, primer_bed_object)

        total_counts[amplicons["right_amplicon"]]["total_reads"] += 1


        # we are searching for the leader sequence
        search = 'AACCAACTTTCGATCTCTTGTAGATCTGTTCT'

        # search for the sequence
        result = search_reads(read,search)

        # add orf location to result
        result["read_orf"] = check_start(orf_bed_object, read)

        # classify read based on prior information
        read_class = classify_read(read,result["align_score"],args.score_cutoff,result["read_orf"],amplicons)

        # store the attributes we have calculated with the read as tags
        read.set_tag('XS', result["align_score"])
        read.set_tag('XA', amplicons["right_amplicon"])
        read.set_tag('XC', read_class)
        read.set_tag('XO', result["read_orf"])


        # ok now add this info to a dictionary for later processing
        if "sgRNA" in read_class:
            if result["read_orf"] is None:
                result["read_orf"] = "novel_"+str(read.pos)

            if result["read_orf"] not in total_counts[amplicons["right_amplicon"]][read_class]:
                total_counts[amplicons["right_amplicon"]][read_class][result["read_orf"]] = []
            total_counts[amplicons["right_amplicon"]][read_class][result["read_orf"]].append(read)
        else:
            total_counts[amplicons["right_amplicon"]][read_class].append(read)

        # write the annotated read to a bam file
        outbamfile.write(read)

    outbamfile.close()
    pysam.index(args.output_prefix + "_periscope.bam")


    # define ORF bed object because we cleared our session
    orf_bed_object = open_bed(args.orf_bed)

    # go through each amplicon and do normalisations
    outfile_amplicons = args.output_prefix + "_periscope_amplicons.csv"
    total_counts,orf_bed_object = calculate_normalised_counts(mapped_reads,total_counts,outfile_amplicons,orf_bed_object)
    # summarise result into ORFs
    result = summarised_counts_per_orf(total_counts,orf_bed_object)
    # output summarised counts
    outfile_counts = args.output_prefix + "_periscope_counts.csv"
    outfile_counts_novel = args.output_prefix + "_periscope_novel_counts.csv"
    output_summarised_counts(mapped_reads,result,outfile_counts,outfile_counts_novel)


if __name__ == '__main__':

    parser = argparse.ArgumentParser(description='periscopre: Search for sgRNA reads in artic network SARS-CoV-2 sequencing data')
    parser.add_argument('--bam', help='bam file',default="The bam file of full artic reads")
    parser.add_argument('--output-prefix',dest='output_prefix',help="Path to the output, e.g. <DIR>/<SAMPLE_NAME>")
    parser.add_argument('--score-cutoff',dest='score_cutoff', help='Cut-off for alignment score of leader (50) we recommend you leave this at 50',default=50)
    parser.add_argument('--orf-bed', dest='orf_bed', help='The bed file with ORF start positions')
    parser.add_argument('--primer-bed', dest='primer_bed', help='The bed file with artic primer positions')
    parser.add_argument('--amplicon-bed', dest='amplicon_bed', help='A bed file of artic amplicons')
    parser.add_argument('--sample', help='sample id',default="SAMPLE")
    parser.add_argument('--tmp',help="pybedtools likes to write to /tmp if you want to write somewhere else define it here",default="/tmp")
    parser.add_argument('--progress', help='display progress bar', default="")


    args = parser.parse_args()

    set_tempdir(args.tmp)

    periscope = main(args)

    if periscope:
        print("all done", file=sys.stderr)
```
