## Supplementary File F19 for "periscope: sub-genomic RNA identification in SARS-CoV-2 Genomic Sequencing Data": Snakefile

```
rule all:
    input:
        # # merged fastq file
        # expand("{output_prefix}.fastq",output_prefix=config.get("output_prefix")),
        # all bam file
        expand("{output_prefix}.bam",output_prefix=config.get("output_prefix")),
        expand("{output_prefix}.bam.bai",output_prefix=config.get("output_prefix")),
        # counts of read types
        expand("{output_prefix}_periscope_counts.csv",output_prefix=config.get("output_prefix")),
        expand("{output_prefix}_periscope_novel_counts.csv",output_prefix=config.get("output_prefix")),
        # amplicon list
        expand("{output_prefix}_periscope_amplicons.csv",output_prefix=config.get("output_prefix")),
        # tagged bam
        expand("{output_prefix}_periscope.bam",output_prefix=config.get("output_prefix")),
        # expand("{output_prefix}_periscope.bam.bai",output_prefix=config.get("output_prefix"))


if config["technology"] == "illumina":

    rule align:
        input:
            fastq = config.get("fastq")
        output:
           expand("{output_prefix}.bam",output_prefix=config.get("output_prefix"))
        params:
            reference = expand("{resources_dir}/{reference_fasta}",resources_dir=config.get("resources_dir"),reference_fasta=config.get("reference_fasta")),
            threads = config.get("threads")
        shell:
            "bwa mem -Y -t {params.threads} {params.reference} {input.fastq} | samtools sort - | samtools view -bh - > {output}"

elif config["technology"] == "ont":

    if config.get('fastq_dir'):
        if config.get('gzipped') == True:
            # collect fastqs
            rule collect_fastqs:
                output:
                    expand("{output_prefix}.{extension}",output_prefix=config.get("output_prefix"),extension=config.get("extension"))
                params:
                    fastq_dir = config.get("fastq_dir"),
                    extension = config.get("extension")
                shell:
                    "zcat {params.fastq_dir}/*.{params.extension} > {output}"
        else:
            # collect fastqs
            rule collect_fastqs:
                output:
                    expand("{output_prefix}.{extension}",output_prefix=config.get("output_prefix"),extension=config.get("extension"))
                params:
                    fastq_dir = config.get("fastq_dir"),
                    extension = config.get("extension")
                shell:
                    "cat {params.fastq_dir}/*.{params.extension} > {output}"


        # map fastqs
        rule align:
            input:
                merged_fastq = expand("{output_prefix}.{extension}",output_prefix=config.get("output_prefix"),extension=config.get("extension"))
            output:
               expand("{output_prefix}.bam",output_prefix=config.get("output_prefix"))
            params:
                reference = expand("{resources_dir}/{reference_fasta}",resources_dir=config.get("resources_dir"),reference_fasta=config.get("reference_fasta")),
                threads = config.get("threads")
            shell:
                "minimap2 -ax map-ont -k 15 -t {params.threads} {params.reference} {input.merged_fastq} | samtools sort - | samtools view -bh - > {output}"

    else:
        rule align:
            input:
                fastq = config.get("fastq")
            output:
               expand("{output_prefix}.bam",output_prefix=config.get("output_prefix"))
            params:
                reference = expand("{resources_dir}/{reference_fasta}",resources_dir=config.get("resources_dir"),reference_fasta=config.get("reference_fasta")),
                threads = config.get("threads")
            shell:
                "minimap2 -ax map-ont -k 15 -t {params.threads} {params.reference} {input.fastq} | samtools sort - | samtools view -bh - > {output}"

# index the bam file
rule index:
    input:
        bam = expand("{output_prefix}.bam",output_prefix=config.get("output_prefix"))
    output:
        expand("{output_prefix}.bam.bai",output_prefix=config.get("output_prefix"))
    shell:
        "samtools index {input.bam}"

#run periscope
rule periscope:
    input:
        bam = expand("{output_prefix}.bam",output_prefix=config.get("output_prefix")),
        bai = expand("{output_prefix}.bam.bai",output_prefix=config.get("output_prefix"))
    output:
        expand("{output_prefix}_periscope.bam",output_prefix=config.get("output_prefix")),
        expand("{output_prefix}_periscope_counts.csv",output_prefix=config.get("output_prefix")),
        expand("{output_prefix}_periscope_amplicons.csv",output_prefix=config.get("output_prefix")),
        expand("{output_prefix}_periscope_novel_counts.csv",output_prefix=config.get("output_prefix"))
        # expand("{output_prefix}_periscope.bam.bai",output_prefix=config.get("output_prefix"))
    params:
        search = expand("{scripts_dir}/search_for_sgRNA_{technology}.py",scripts_dir=config.get("scripts_dir"),technology=config.get("technology")),
        output_prefix = config.get("output_prefix"),
        score_cutoff = config.get("score_cutoff"),
        sample = config.get("sample"),
        orf_bed = expand("{resources_dir}/{orf_bed}",resources_dir=config.get("resources_dir"),orf_bed=config.get("orf_bed")),
        primer_bed = expand("{resources_dir}/{primer_bed}",resources_dir=config.get("resources_dir"),primer_bed=config.get("primer_bed")),
        amplicon_bed =expand("{resources_dir}/{amplicon_bed}",resources_dir=config.get("resources_dir"),amplicon_bed=config.get("amplicon_bed")),
        tmp = config.get("tmp")
    shell:
        "python {params.search} --bam {input.bam} --score-cutoff {params.score_cutoff} --output-prefix {params.output_prefix} --score-cutoff {params.score_cutoff} --sample {params.sample} --orf-bed {params.orf_bed} --primer-bed {params.primer_bed} --amplicon-bed {params.amplicon_bed} --tmp {params.tmp}"
```
