## Supplementary File F19 for "periscope: sub-genomic RNA identification in SARS-CoV-2 Genomic Sequencing Data": variant_expression.py

```
import vcf
import pysam
import argparse
import pandas as pd
from plotnine import *


def check_position_in_bam(bam,position):
    """
    using pysam pileup get the base composition at a position and split counts based on XC read tag

    :param bam: periscpe bam file
    :param position: position of interest
    :return:
    """
    result={}
    for pileupcolumn in bam.pileup("MN908947.3",position,position+1):

        # deal with 0 and 1 based shenanigans
        if pileupcolumn.pos==position-1:
            for pileupread in pileupcolumn.pileups:
                if not pileupread.is_del and not pileupread.is_refskip:
                    # get the read class that periscope assigned previously
                    read_class = pileupread.alignment.get_tag("XC")
                    if read_class not in result:
                        result[read_class] = {}

                    # get the base
                    base = pileupread.alignment.query_sequence[pileupread.query_position]
                    if base not in result[read_class]:
                        result[read_class][base]=0

                    # add to the count of bases for the read class at this position
                    result[read_class][base] += 1


    return result

def main(args):
    # open vcf file
    vcf_reader = vcf.Reader(open(args.vcf, 'r'))
    # open bam file
    bam = pysam.AlignmentFile(args.bam, "rb")


    result = []
    poses = []

    for record in vcf_reader:
        # do the work
        check = check_position_in_bam(bam, record.POS)
        # get the result
        poses.append(record.POS)
        result.append(pd.DataFrame.from_dict(check))

    # make a pandas dataframe with the data
    df = pd.concat(result,keys=poses)

    df.reset_index(inplace = True)
    df['level_0']=df['level_0'].astype(str)
    print(df)

    # melt the dataframe
    df = pd.melt(df,id_vars=['level_0','level_1'])

    # do some stuff to get the order of positions correct for plotting
    pos_list = df['level_0'].value_counts().index.tolist()
    pos_cat = pd.Categorical(df['level_0'], categories=pos_list)
    df = df.assign(pos_cat=pos_cat)

    # plot the result using ggplot
    x=(ggplot(df, aes(x='pos_cat',y='value',color='level_1',fill='level_1'))) + \
      geom_bar(stat='identity',position='fill') + \
      facet_wrap(['variable'],ncol=1)

    x.save(filename=args.output_prefix+"_base_counts.png", height=5, width=5, units='in', dpi=300)

    df = df.rename(columns={"level_0": "position", "level_1": "base"})
    df = df.assign(sample=args.sample)

    # write the result to a file
    df.to_csv(args.output_prefix+"_base_counts.csv",index=False)

    print(x)

if __name__ == '__main__':

    parser = argparse.ArgumentParser(description='periscope: Get frequencies of bases at variant positions in different read classes')
    parser.add_argument('--periscope-bam',dest="bam", help='Bam file from periscope')
    parser.add_argument('--output-prefix',dest='output_prefix', help='Prefix of the output file i.e. <DIRECTORY>/<FILE_PREFIX>')
    parser.add_argument('--vcf', dest='vcf', help='Non-gz compressed VCF file')
    parser.add_argument('--sample', help='Sample identifier')

    args = parser.parse_args()
    main(args)
```
