## Supplementary File F19 for "periscope: sub-genomic RNA identification in SARS-CoV-2 Genomic Sequencing Data": README.md

![alt text](https://github.com/sheffield-bioinformatics-core/periscope/blob/master/periscope.png "periscope")

**v0.0.8 With improved Illumina sgRNA detection**

# periscope

A tool to quantify sub-genomic RNA (sgRNA) expression in SARS-CoV-2 artic network amplicon sequencing data.
Initial classification of reads into sub-genomic or not based on https://www.biorxiv.org/content/10.1101/2020.04.10.029454v1.abstract

# Citing

Please cite our pre-print if you use this tool in any publications:

https://www.biorxiv.org/content/10.1101/2020.07.01.181867v1

# Requirements
periscope runs on MacOS and Linux. We have also confirmed the tool runs under windows 10 unix subsystem.


* conda
* Your raw fastq files from the artic protocol
* Install periscope

# Installation
```
git clone https://github.com/sheffield-bioinformatics-core/periscope.git && cd periscope
conda env create -f environment.yml
conda activate periscope
pip install .
```

# Execution

```
conda activate periscope

periscope \
    --fastq-dir <PATH_TO_DEMUXED_FASTQ> \ (ont only)
    OR
    --fastq <FULL_PATH_OF_FASTQ_FILE(s)> \ (space separated list of fastq files, you MUST use this for Illumina data)
    --output-prefix <PREFIX> \
    --sample <SAMPLE_NAME> \
    --artic-primers <ASSAY_VERSION; V1,V2,V3 or 2kb> \
    --resources <PATH_TO_PERISCOPE_RESOURCES_FOLDER> \
    --technology <SEQUECNING TECH; ont or illumina> \
    --threads <THREADS_FOR_MAPPING> 
```

`output-prefix` will be the directory and start of the filename for the output.

So if you put `./SAMPLE1` for this argument outputs will go in the current working directory prefixed by "SAMPLE1". 

***Note*** - for illumina data please use --fastq <FASTQ_R1>.fastq.gz <FASTQ_R2>.fastq.gz and --technology illumina

## `/tmp` Issues

If you have issues with `tmp` this is because pybedtools writes there. v0.0.3 contains a fix, and you can also specify `--tmp` and redirect this somewhere else

# Pipeline overview

![alt text](https://github.com/sheffield-bioinformatics-core/periscope/blob/master/workflow.png "periscope")<!-- .element height="10%" width="10%" -->
__Figure 1. Workflow Overview__ 

## Pre-Processing

* Collect demutiplexed pass fastqs
* Remap RAW artic protocol reads

## Counting

_This step takes roughly 1minute per 10k reads_
_Our median read count is ~250k and this will take around 25minutes_

* Read bam file
* Filter unmapped and secondary alignments
* Assign amplicon to read (using artic align_trim.py)
* Search for leader sequence
* Assign read to ORF
* Classify read (see Figure 2)
* Normalise a few ways

![alt text](https://github.com/sheffield-bioinformatics-core/periscope/blob/master/read_classification.png "periscope")<!-- .element height="10%" width="10%" -->
__Figure 2. Read Classification Algorithm__ 

## Normalisation

### ONT Data

We have taken two approaches, a global normalisation based on mapped read counts or a local normalisation based on gRNA from the same amplicon.

* gRNA or sgRNA Per 100,000 mapped reads (gRPHT or sgRPHT)
    * We do this per amplicon and sum them in instances where multiple amplicons contribute to the final ORF count
* sgRNA can be normalised to per 1000 gRNA reads from the same amplicon - sgRPTg (normalising for amplicon efficiency differences)
    * There are things you need to note here:
        * multiple amplicons can contribute to reads which support the same sgRNA
        * we normalise on a per amplicon level and then sum these to get an overall normalised count

### Illumina Data

Ilumina data is still a work in progress, as of v0.0.8 you can get raw sgRNA counts and counts normalised to the average coverage around the ORF TRS start site.

It is worth noting this follows a slightly different algorithm, relying instead on soft clipping. The ratoinale here is that illumina data is more accurate therefore we
can detect shorter matches to the leader.

- 


## Outputs:

#### <OUTPUT_PREFIX>.fastq

A merge of all files in the fastq directory specified as input.

#### <OUTPUT_PREFIX>_periscope_counts.csv

The counts of genomic, sub-genomic and normalisation values for known ORFs

#### <OUTPUT_PREFIX>_periscope_amplicons.csv

The amplicon by amplicon counts, this file is useful to see where the counts come from. Multiple amplicons may be represented more than once where they may have contributed to more than one ORF.

#### <OUTPUT_PREFIX>_periscope_novel_counts.csv

The counts of genomic, sub-genomic and normalisation values for non-canonical ORFs

#### <OUTPUT_PREFIX>.bam

minmap2 mapped reads and index with no adjustments made.

#### <OUTPUT_PREFIX>_periscope.bam

This is the original input bam file and index created by periscope with the reads specified in the fastq-dir. This file, however, has tags which represent the results of periscope:

- XS is the alignment score
- XA is the amplicon number
- XC is the assigned class (gDNA or sgDNA)
- XO is the orf assigned

These are useful for manual review in IGV or similar genome viewer. You can sort or colour reads by these tags to aid in manual review and figure creation.


# Extracting Base Frequencies

To examine the composition of bases at variant sites we have provided this code.
```
conda activate periscope

gunzip <ARTIC_NETWORK_VCF>.pass.vcf.gz

<PATH_TO_PERISCOPE>/periscope/periscope/scripts/variant_expression.py \
    --periscope-bam <PATH_TO_PERISCOPE_OUTPUT_BAM> \
    --vcf <ARTIC_NETWORK_VCF>.pass.vcf \
    --sample <SAMPLE_NAME> \
    --output-prefix <PREFIX>
```

#### <OUTPUT_PREFIX>_base_counts.csv

Counts of each base at each position

#### <OUTPUT_PREFIX>_base_counts.png

Plot of each position and base composition

# Running Tests

We provide a sam file for testing the main module of periscope.

reads.sam contains 23 reads which have been manually reviewed for the truth

```
cd <INSTALLATION_PATH>/periscope/tests

pytest test_search_for_sgRNA.py 
```

# Citations

### Long Amplicon Tiling Data

We implemened 2kb amplicon tilling in v0.0.2 from:

**SARS-CoV-2 genomes recovered by long amplicon tiling multiplex approach using nanopore sequencing and applicable to other sequencing platforms**
Paola Cristina Resende, Fernando Couto Motta, Sunando Roy, Luciana Appolinario, Allison Fabri, Joilson Xavier, Kathryn Harris, Aline Rocha Matos, Braulia Caetano, Maria Orgeswalska, Milene Miranda, Cristiana Garcia, André Abreu, Rachel Williams, Judith Breuer, Marilda M Siqueira
bioRxiv 2020.04.30.069039; doi: https://doi.org/10.1101/2020.04.30.069039


Why periscope? SUB-genomic RNA, SUB-marine, periscope.
