## Supplementary File F19 for "periscope: sub-genomic RNA identification in SARS-CoV-2 Genomic Sequencing Data": setup.py

```
from setuptools import setup
from periscope import __version__, _program

setup(
    name='periscope',
    version=__version__,
    packages=['periscope'],
    scripts=['periscope/scripts/Snakefile',
             'periscope/scripts/search_for_sgRNA_ont.py',
             'periscope/scripts/search_for_sgRNA_illumina.py',
             'periscope/scripts/variant_expression.py'
             ],
    url='',
    license='',
    author='Matthew Parker',
    author_email='',
    description='periscope searches for and quanifies sgRNAs in SARS-CoV-2',
    entry_points="""
    [console_scripts]
    {program} = periscope.periscope:main
    """.format(program=_program),
    include_package_data=True,
)
```
