## Supplementary File F19 for "periscope: sub-genomic RNA identification in SARS-CoV-2 Genomic Sequencing Data": test_search_for_sgRNA_illumina.py

```
# we have a sam file of reads from real samples that give examples
# ReadID    ORF Amplicon    Reason
# NB500959:197:H75W3AFX2:4:11403:13826:18844    ORF7a   90  sgRNA
# NB500959:197:H75W3AFX2:1:21312:23610:4695 ORF7a   90  sgRNA - smaller leader
# NB500959:197:H75W3AFX2:1:11308:21480:4281 ORF7a   90  sgRNA - smaller leader
# NB500959:197:H75W3AFX2:2:11212:14327:14989    ORF7a   90   gRNA
# NB500959:197:H75W3AFX2:4:11507:13616:10003    ORF7a   90  sgRNA longest leader
# NB500959:197:H75W3AFX2:2:11306:20696:15959    ORF7a   90  gRNA
# M01996:271:000000000-J3T3D:1:1101:23402:7678 has soem soft clipping but it is minimal - what is this one?
# M01996:271:000000000-J3T3D:1:1106:16147:13747 borderline - probably not a match
# M01996:271:000000000-J3T3D:1:2103:7946:12236 right read really messy but softclipped so matches somewhat to leader

# TODO - I want some amplicon 86 reads that support E
# TODO - I need some reads supporting 7a

# Import all the methods we need
from periscope.scripts.search_for_sgRNA_illumina import search_reads, classify_read, find_amplicon, get_mapped_reads, check_start, open_bed, supplementary_method, extact_soft_clipped_bases

# this is the truth for these reads

truth = {
    "NB500959:197:H75W3AFX2:4:11403:13826:18844": {
        "soft_clip_check": True,
        "class":"sgRNA",
        "align_score":64.0,
        "amplicon": 90,
        "orf": "ORF7a"
    },
    "NB500959:197:H75W3AFX2:1:21312:23610:4695": {
        "soft_clip_check": True,
        "class":"sgRNA",
        "align_score":52.0,
        "amplicon": 90,
        "orf": "ORF7a"
    },
    "NB500959:197:H75W3AFX2:1:11308:21480:4281": {
        "soft_clip_check": True,
        "class": "sgRNA",
        "align_score": 32.0,
        "amplicon": 90,
        "orf": "ORF7a"
    },
    "NB500959:197:H75W3AFX2:2:11212:14327:14989": {
        "soft_clip_check": False,
        "class":"gRNA",
        "align_score": 12.0,
        "amplicon": 90,
        "orf": None
    },
    "NB500959:197:H75W3AFX2:4:11507:13616:10003": {
        "soft_clip_check": True,
        "class":"sgRNA",
        "align_score":64.0,
        "amplicon": 90,
        "orf": "ORF7a"
    },
    "NB500959:197:H75W3AFX2:2:11306:20696:15959": {
        "soft_clip_check": False,
        "class":"gRNA",
        "align_score": 12.0,
        "amplicon": 90,
        "orf": None
    },
    "NB500959:197:H75W3AFX2:1:21109:16155:11370": {
            "soft_clip_check": False,
            "class":"gRNA",
            "align_score": 12.0,
            "amplicon": 90,
            "orf": None
        },
    "NB500959:197:H75W3AFX2:3:21408:12117:17553":{
            "soft_clip_check": False,
            "class":"gRNA",
            "align_score": 12.0,
            "amplicon": 90,
            "orf": None
    },
    "M01996:271:000000000-J3T3D:1:1101:23402:7678":{

            "soft_clip_check": False,
            "class":"gRNA",
            "align_score": 12.0,
            "amplicon": 90,
            "orf": None

    },
    "M01996:271:000000000-J3T3D:1:1101:20572:27704":{

    }


}


import pysam
import os
from artic.vcftagprimersites import read_bed_file


dirname = os.path.dirname(__file__)

def test_mapped_reads():
    mapped_reads = get_mapped_reads("reads.sam")
    assert mapped_reads == 23


def test_check_start():
    inbamfile = pysam.AlignmentFile("reads.sam", "rb")
    filename = os.path.join(dirname, "../../periscope/resources/orf_start.bed")
    bed_object = open_bed(filename)
    for read in inbamfile:
        if read.is_supplementary:
            # print("%s skipped as supplementary" %
            #       (read.query_name), file=sys.stderr)
            continue
        if read.is_secondary:
            # print("%s skipped as supplementary" %
            #       (read.query_name), file=sys.stderr)
            continue
        print(read.query_name)
        orf = check_start(bed_object,read)
        assert orf == truth[read.query_name]["orf"]


def test_supplementary_method():
    inbamfile = pysam.AlignmentFile("reads.sam", "rb")
    for read in inbamfile:

        print("------")
        print(read.query_name)
        # print(read.is_read1)
        # print(read.get_tags())
        print(supplementary_method(read))

def test_extact_soft_clipped_bases():
    inbamfile = pysam.AlignmentFile("reads.sam", "rb")
    result = {}
    count = 0
    for read in inbamfile:
        if read.is_supplementary:
            continue
        if read.is_secondary:
            # print("%s skipped as supplementary" %
            #       (read.query_name), file=sys.stderr)
            continue
        print("------")
        print(read.query_name)
        # print(read.is_read1)
        # print(read.get_tags())
        print(read.cigar)

        test = extact_soft_clipped_bases(read)

        old_result = None

        if read.query_name in result:

            old_result = result[read.query_name]
        if old_result == None:
            result[read.query_name] = test
        elif old_result == False:
            if test == True:
                result[read.query_name] = True

        count+=1

    print(count)

    print(result)
    truth = {
                'M01996:271:000000000-J3T3D:1:1101:23402:7678': False,
                'M01996:271:000000000-J3T3D:1:1106:16147:13747': False,
                'M01996:271:000000000-J3T3D:1:1101:20572:27704': False,
                'NB500959:197:H75W3AFX2:1:11308:21480:4281': True,
                'NB500959:197:H75W3AFX2:1:21109:16155:11370': False,
                'NB500959:197:H75W3AFX2:1:21312:23610:4695': True,
                'NB500959:197:H75W3AFX2:2:11204:7045:4512': True,
                'NB500959:197:H75W3AFX2:2:11212:14327:14989': False,
                'NB500959:197:H75W3AFX2:2:11306:20696:15959': False,
                'NB500959:197:H75W3AFX2:3:21408:12117:17553': False,
                'NB500959:197:H75W3AFX2:4:11403:13826:18844': True,
                'NB500959:197:H75W3AFX2:4:11507:13616:10003': True,
                'NB500959:197:H75W3AFX2:4:11602:22252:8721': True
            }
    assert result == truth


def test_search_reads():

    inbamfile = pysam.AlignmentFile("reads.sam", "rb")
    for read in inbamfile:
        if read.is_supplementary:
            # print("%s skipped as supplementary" %
            #       (read.query_name), file=sys.stderr)
            continue
        if read.is_secondary:
            # print("%s skipped as supplementary" %
            #       (read.query_name), file=sys.stderr)
            continue

        search = 'AACCAACTTTCGATCTCTTGTAGATCTGTTCT'
        result = search_reads(read, search)
        print(read.query_name)
        print(read.flag)
        print(result)
        assert result["align_score"] == truth[read.query_name]["align_score"]


def test_find_amplicon():

    filename = os.path.join(dirname, "../periscope/resources/artic_primers_V3.bed")
    primer_bed_object = read_bed_file(filename)
    inbamfile = pysam.AlignmentFile("reads.sam", "rb")
    for read in inbamfile:
        amplicon = find_amplicon(read,primer_bed_object)["right_amplicon"]
        assert amplicon == truth[read.query_name]["amplicon"]


def test_classify_read():
    inbamfile = pysam.AlignmentFile("reads.sam", "rb")

    filename = os.path.join(dirname, "../periscope/resources/artic_primers_V3.bed")
    primer_bed_object = read_bed_file(filename)

    filename = os.path.join(dirname, "../periscope/resources/orf_start.bed")
    bed_object = open_bed(filename)

    for read in inbamfile:
        print(read.query_name)
        search = 'AACCAACTTTCGATCTCTTGTAGATCTGTTCT'
        search_result = search_reads(read,search)
        amplicons = find_amplicon(read, primer_bed_object)
        orf = check_start(bed_object, read)
        result = classify_read(read,search_result["align_score"],50,orf,amplicons)
        print(result)
        assert result == truth[read.query_name]["class"]


def test_pybedtools():
    import pybedtools
    read_feature = pybedtools.BedTool("MN908947.3" + "\t" + str(0) + "\t" + str(0),from_string=True)
    for bed_line in read_feature:
        assert bed_line.chrom == "MN908947.3"
        assert bed_line.start == 0
        assert bed_line.end == 0
```
