## Supplementary File F19 for "periscope: sub-genomic RNA identification in SARS-CoV-2 Genomic Sequencing Data": test_search_for_sgRNA_ont.py

```
# we have a sam file of reads from real samples that give examples
# ReadID    ORF Amplicon    Reason
# 90b89b24-9c38-456b-a11e-12bf1712bb06  S   71  sgRNA   Good leader match pool 1 amplicon
# fe0e82e6-6239-4dc2-be2f-98a08860b3d7  S   71  sgRNA   Bad leader match (<50)
# 806f2bd2-3a11-4e54-9c2e-51e54aa3e94d  S   72  sgRNA   This is a read which results from amplicon 72 but looks to be an S sgRNA (WHAT HAPPENS TO THIS READ?)
# 25686e6f-b0b8-4bad-9cbe-d8e6de270648  S   71  gRNA
# 960eb82c-aa1d-46ec-8358-2f9cf496c7cc  S   71  gRNA
# 8319b096-6508-4a37-a517-775133f2adfc  None    72  gRNA
# 3ad58c30-b3b1-4ad7-bba6-0921d6a11ab2  None    72  gRNA
# 5125ffa6-2567-4ba2-9f05-e7f8a161aa11  E   87  sgRNA   E support from amplicon 87
# b7b9fac8-8b27-40b8-9d02-8edc59c82f39  M   87  sgRNA   Good match
# 8092cf1e-d132-4abc-ac3d-4ccb22dbb932  M   87  gRNA
# 5da3aeaa-f955-4b9a-89dd-7bdd6e3b77fe  M   87  gRNA
# 90928d27-3c71-45f2-b31c-3d7de36c78b8  ORF6    89  sgRNA   ORF6 supporting read from amplicon 89
# 4d67e504-7d7d-4b11-aea2-3ec606195a1b  ORF6    90  sgRNA   ORF6 supporting read from amplicon 90
# 8a8c2527-72b2-4ca1-9570-f8f00f336de0  ORF6    90  gRNA
# b07a9f12-ff5f-4df0-bf2b-73a072921392  ORF6    89  gRNA
# 1ec155b3-120c-4181-86ef-b31c186d4651  N   93  sgRNA   Bad leader match (classified as gRNA at 50)
# e0d5f838-1aca-44d2-a001-e47d7c0edb06  N   93  sgRNA   Good leader match
# 21af3571-b970-4d1e-ad1c-3f95d1fa150e  N   93  gRNA
# c85c93e5-427d-448d-940b-ea8e6ec2ac6a  N   93  gRNA
# e8981b0c-21fe-4920-80e3-530197f3d15e  novel_20315   67  sgRNA   Novel sgRNA
# 07b675cc-6b19-4ed4-9341-6576ad51957f  None    67  gRNA
# 76aaf579-4754-4bbe-b001-4ac2d6f76533  novel_19548   65  gRNA    listed as novel, but on edge of amplicon
# 1e0b6284-ec08-4451-b0fc-3f7f36e75b31 None    97  gRNA

# TODO - I want some amplicon 86 reads that support E
# TODO - I need some reads supporting 7a

# Import all the methods we need
from periscope.scripts.search_for_sgRNA_ont import search_reads, classify_read, find_amplicon, get_mapped_reads, check_start, open_bed

# this is the truth for these reads

truth = {
    "90b89b24-9c38-456b-a11e-12bf1712bb06": {
        "class":"sgRNA_HQ",
        "align_score":64.0,
        "amplicon": 71,
        "orf": "S"
    },
    "fe0e82e6-6239-4dc2-be2f-98a08860b3d7": {
        "class":"sgRNA_LQ",
        "align_score":46.0,
        "amplicon": 71,
        "orf": "S"
    },
    "806f2bd2-3a11-4e54-9c2e-51e54aa3e94d": {
        "class": "sgRNA_HQ",
        "align_score": 60.0,
        "amplicon": 72,
        "orf": "S"
    },
    "25686e6f-b0b8-4bad-9cbe-d8e6de270648": {
        "class": "gRNA",
        "align_score": 16.0,
        "amplicon": 71,
        "orf": None
    },
    "960eb82c-aa1d-46ec-8358-2f9cf496c7cc": {
        "class": "gRNA",
        "align_score": 14.0,
        "amplicon": 71,
        "orf": None
    },
    "8319b096-6508-4a37-a517-775133f2adfc": {
        "class": "gRNA",
        "align_score": 14.0,
        "amplicon": 72,
        "orf": None
    },
    "3ad58c30-b3b1-4ad7-bba6-0921d6a11ab2": {
        "class": "gRNA",
        "align_score": 14.0,
        "amplicon": 72,
        "orf": None
    },
    "5125ffa6-2567-4ba2-9f05-e7f8a161aa11": {
        "class": "sgRNA_HQ",
        "align_score": 54.0,
        "amplicon": 87,
        "orf": "E"
    },
    "b7b9fac8-8b27-40b8-9d02-8edc59c82f39": {
        "class": "sgRNA_HQ",
        "align_score": 54.0,
        "amplicon": 87,
        "orf": "M"
    },
    "8092cf1e-d132-4abc-ac3d-4ccb22dbb932": {
        "class": "gRNA",
        "align_score": 15.0,
        "amplicon": 87,
        "orf": None
    },
    "5da3aeaa-f955-4b9a-89dd-7bdd6e3b77fe": {
        "class": "gRNA",
        "align_score": 14.799999999999999,
        "amplicon": 87,
        "orf": None
    },
    "90928d27-3c71-45f2-b31c-3d7de36c78b8": {
        "class": "sgRNA_HQ",
        "align_score": 54.0,
        "amplicon": 89,
        "orf": "ORF6"
    },
    "4d67e504-7d7d-4b11-aea2-3ec606195a1b": {
        "class": "sgRNA_HQ",
        "align_score": 56.0,
        "amplicon": 90,
        "orf": "ORF6"
    },
    "8a8c2527-72b2-4ca1-9570-f8f00f336de0": {
        "class": "gRNA",
        "align_score": 12.0,
        "amplicon": 90,
        "orf": None
    },
    "b07a9f12-ff5f-4df0-bf2b-73a072921392": {
        "class": "gRNA",
        "align_score": 14.799999999999997,
        "amplicon": 89,
        "orf": None
    },
    "1ec155b3-120c-4181-86ef-b31c186d4651": {
        "class": "sgRNA_LLQ",
        "align_score": 26.0,
        "amplicon": 93,
        "orf": "N"
    },
    "e0d5f838-1aca-44d2-a001-e47d7c0edb06": {
        "class": "sgRNA_HQ",
        "align_score": 56.0,
        "amplicon": 93,
        "orf": "N"
    },
    "21af3571-b970-4d1e-ad1c-3f95d1fa150e": {
        "class": "gRNA",
        "align_score": 16.0,
        "amplicon": 93,
        "orf": None
    },
    "c85c93e5-427d-448d-940b-ea8e6ec2ac6a": {
        "class": "gRNA",
        "align_score": 14.0,
        "amplicon": 93,
        "orf": None
    },
    "e8981b0c-21fe-4920-80e3-530197f3d15e": {
        "class": "nsgRNA_HQ",
        "align_score": 64.0,
        "amplicon": 67,
        "orf": None
    },
    "07b675cc-6b19-4ed4-9341-6576ad51957f": {
        "class": "gRNA",
        "align_score": 14.0,
        "amplicon": 67,
        "orf": None
    },
    "76aaf579-4754-4bbe-b001-4ac2d6f76533": {
        "class": "gRNA",
        "align_score": 64.0,
        "amplicon": 65,
        "orf": None
    },
    "1e0b6284-ec08-4451-b0fc-3f7f36e75b31": {
        "class": "gRNA",
        "align_score": 14.0,
        "amplicon": 97,
        "orf": None
    }


}


import pysam
import os
from artic.vcftagprimersites import read_bed_file


dirname = os.path.dirname(__file__)

def test_mapped_reads():
    mapped_reads = get_mapped_reads("reads.sam")
    assert mapped_reads == 23


def test_check_start():
    inbamfile = pysam.AlignmentFile("reads.sam", "rb")
    filename = os.path.join(dirname, "../../periscope/resources/orf_start.bed")
    bed_object = open_bed(filename)
    for read in inbamfile:
        orf = check_start(bed_object,read)
        assert orf == truth[read.query_name]["orf"]


def test_search_reads():

    inbamfile = pysam.AlignmentFile("reads.sam", "rb")
    for read in inbamfile:
        search = 'AACCAACTTTCGATCTCTTGTAGATCTGTTCT'
        result = search_reads(read, search)
        assert result["align_score"] == truth[read.query_name]["align_score"]


def test_find_amplicon():

    filename = os.path.join(dirname, "../../periscope/resources/artic_primers_V3.bed")
    primer_bed_object = read_bed_file(filename)
    inbamfile = pysam.AlignmentFile("reads.sam", "rb")
    for read in inbamfile:
        amplicon = find_amplicon(read,primer_bed_object)["right_amplicon"]
        assert amplicon == truth[read.query_name]["amplicon"]


def test_classify_read():
    inbamfile = pysam.AlignmentFile("reads.sam", "rb")

    filename = os.path.join(dirname, "../../periscope/resources/artic_primers_V3.bed")
    primer_bed_object = read_bed_file(filename)

    filename = os.path.join(dirname, "../../periscope/resources/orf_start.bed")
    bed_object = open_bed(filename)

    for read in inbamfile:
        print(read.query_name)
        search = 'AACCAACTTTCGATCTCTTGTAGATCTGTTCT'
        search_result = search_reads(read,search)
        amplicons = find_amplicon(read, primer_bed_object)
        orf = check_start(bed_object, read)
        result = classify_read(read,search_result["align_score"],50,orf,amplicons)
        print(result)
        assert result == truth[read.query_name]["class"]


def test_pybedtools():
    import pybedtools
    read_feature = pybedtools.BedTool("MN908947.3" + "\t" + str(0) + "\t" + str(0),from_string=True)
    for bed_line in read_feature:
        assert bed_line.chrom == "MN908947.3"
        assert bed_line.start == 0
        assert bed_line.end == 0
```
