## Supplementary figures and images for "periscope: sub-genomic RNA identification in SARS-CoV-2 Genomic Sequencing Data"

### periscope.png

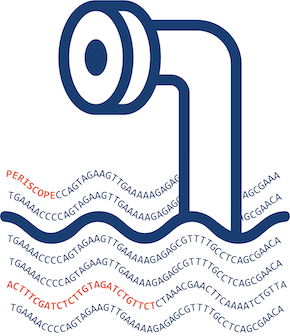

### read_classification.png

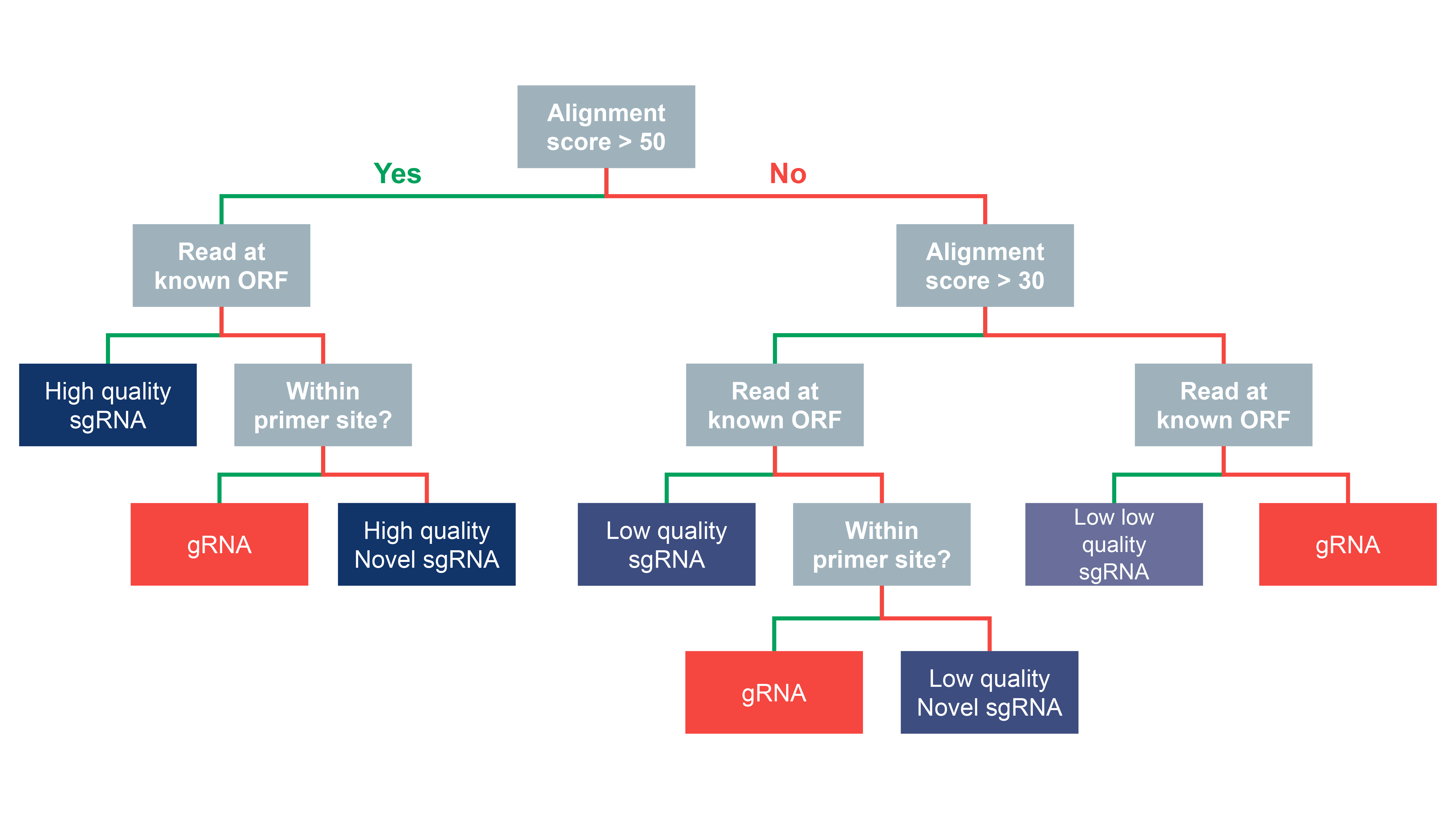

### workflow.png

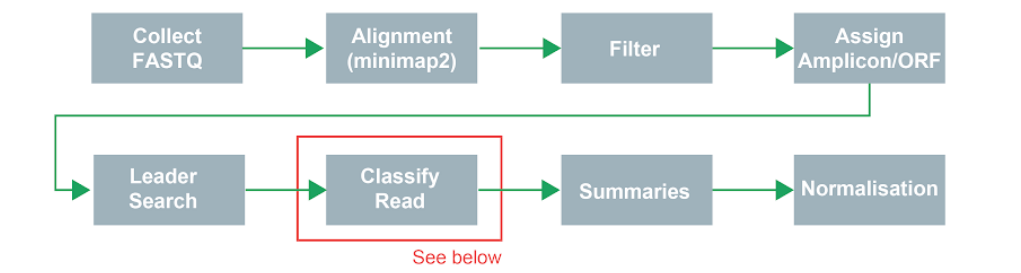
